# Carcinogen-induced DNA adducts amplify APOBEC mutagenesis through nucleotide excision repair

**DOI:** 10.1101/2025.01.18.633716

**Authors:** Cameron Durfee, Bojana Stefanovska, Rongjie Wu, José Córdoba-Caballero, Mahmoud A. Ibrahim, Michael A. Carpenter, Liping Liu, Shuvro P. Nandi, Prokopios P. Argyris, Luca Boscolo-Bielo, Eric C. L. Li, Nuri Alpay Temiz, Yufan Zhou, Erik N. Bergstrom, Christopher D. Mullally, Yaxi Wang, Xingyu Liu, Emilia Barreto Duran, Benjamin Troness, Joshua Proehl, Harshita B. Gupta, Allen J. York, Rachel I. Vogel, Jacob M. Wells, Christopher D. Steele, Isabella Stuewe, Daniel Muldoon, Jun Woo, Mark T. A. Donoghue, Laura A. Lindsey-Boltz, Aziz Sancar, Chaitanya Bandlamudi, Sarat Chandarlapaty, Wei Yang, Anna Malkova, Marcos Díaz-Gay, Ludmil B. Alexandrov, Reuben S. Harris

## Abstract

Extrinsic and intrinsic factors induce mutations in somatic cells that play a key role in cancer development. Tobacco smoke contains carcinogens that contribute to cancers of the lung and head & neck^1,2^. The APOBEC family of DNA cytosine deaminases is a prominent endogenous source of mutation, generating characteristic signatures (SBS2 and SBS13) that often co-occur with the tobacco-associated signature SBS4 in tumors from smokers^3,4^. Here, we challenge the prevailing view that mutational processes act independently and additively. We show that 4-nitroquinoline 1-oxide (NQO), a widely used model for tobacco carcinogenesis, sensitizes cells to APOBEC3B (A3B)-mediated mutagenesis, resulting in a significant increase in oral carcinoma development *in vivo*. NQO exposure markedly amplifies APOBEC-associated SBS2 mutagenesis, without a corresponding increase in NQO-associated SBS4 mutations. Moreover, a large proportion of A3B-catalyzed SBS2 mutations occur as strand-coordinated pairs within 32 nucleotides of each other, supporting a model in which nucleotide excision repair (NER) of NQO-induced DNA adducts generates short single-stranded DNA intermediates that are accessible to APOBEC-catalyzed deamination. These APOBEC mutation pairs—termed *didyma* (Greek for twins)—exhibit distinct genomic features consistent with a NER-initiated mechanism, differing from other types of clustered mutations (*omikli* and *kataegis*). A3B exhibits processive deamination on NER-like tracts *in vitro*. *Didyma* formation in human cell lines and yeast depends on NQO exposure, catalytically active A3B, and functional NER (XPA in human cells and Rad14 in yeast). Computational analyses of lung and head & neck tumor genomes show that both APOBEC mutagenesis and *didyma* are elevated in cancers from smokers compared to non-smokers. Elevated levels are also observed in normal lung tissues from smokers prior to tumor development. Additional tumor types without obvious links to smoking also exhibit *didyma*. Collectively, these results indicate that exogenous DNA adduct-forming carcinogens can amplify mutagenesis by endogenous APOBEC enzymes and that mutational processes can interact synergistically to shape tumor genomes during cancer initiation and progression.

## MAIN

Studies on somatic mutagenesis have been revolutionized by advances in DNA sequencing technologies and computational methods for analyzing large genomic datasets. These advancements have enabled the mapping of somatic mutation patterns imprinted by distinct mutational processes, termed *mutational signatures*. Over 100 single base substitution (SBS) mutational signatures have been described in human cancers, with approximately half assigned to putative mechanistic etiologies that can be broadly categorized as arising from endogenous cellular processes or exogenous environmental factors^3,5^. Major endogenous sources include unavoidable chemical reactions such as spontaneous deamination of 5-methyl-cytosine to thymine (C-to-T in SBS1)^6–8^ and oxidation of guanine to 8-oxoG leading to G-to-T transversions (C-to-A in SBS18)^9^, as well as defects in normal DNA repair processes such as homologous recombination (broad mutational spectrum in SBS3 as well as larger-scale genomic aberrations)^7,10,11^. Exogenous examples include tobacco smoke, ultraviolet (UV) light, and aristolochic acid^3^. Tobacco smoke contains many chemical mutagens but the most potent classes are polycyclic aromatic hydrocarbons (such as benzo[*a*]pyrene) and tobacco-specific nitrosamines (such as *N’*-nitrosonornicotine)^2,12^. Upon metabolic activation, both classes form adducts with guanine nucleobases and, to a lesser extent, with adenines and thymines, leading to the major G-to-T (C-to-A) mutational contribution in SBS4^12,13^. UV light cross-links adjacent pyrimidine bases, which results in error-prone lesion bypass synthesis (A-insertion) by DNA polymerases and C-to-T mutations in SBS7a/b^14^. Aristolochic acid is processed into DNA-reactive compounds that lead to A-to-T transversions (T-to-A in SBS22)^14^. Many other endogenous and exogenous sources of DNA damage also contribute to human mutational signatures. However, regardless of etiology, mutational processes in normal as well as cancerous cells are assumed to occur independently, with additive impact throughout the lifespan of a cell, tissue, or full organism^8,15,16^.

Cellular APOBEC3 enzymes, predominantly APOBEC3B (A3B) and APOBEC3A (A3A)^17–20^, generate SBS2 and SBS13 mutational signatures and represent a common mutational process in human cancer (only following aging-associated processes; *e.g.*, SBS1 and SBS5)^3^. A3B and A3A normally function alongside other family members in innate antiviral immunity^21^. However, it is now clear that approximately 70% of all cancer types are impacted by their mutagenic activity, especially tumors of the bladder, breast, cervix, head & neck, and lung tissues^3,7,22–25^.

A3B and A3A catalyze the hydrolytic deamination of cytosine-to-uracil preferentially in TCA and TCT trinucleotide motifs in single-stranded (ss)DNA^26–28^. The resulting uracil lesions can either template the insertion of adenines during DNA replication (leading to C-to-T in SBS2) or be excised by uracil DNA glycosylase and converted into abasic sites. Abasic sites can mis-template the insertion of adenines during DNA replication (also leading to SBS2) or provoke lesion bypass synthesis by the deoxycytidyl transferase REV1 (leading to C-to-G and C-to-A in SBS13)^18^. Additional processing of abasic sites can result in single- and double-stranded DNA breaks and consequently other types of mutations including insertion/deletion mutations (indels) and larger-scale structural variations^17,19,29^. Most APOBEC3 signature mutations in cancer are dispersed and associated with ssDNA intermediates in DNA replication, transcription, and recombination^30–35^. However, APOBEC3 mutations also occur in clusters, including *omikli*, 2-3 mutations associated with DNA mismatch repair, and *kataegis*, ≥4 strand-coordinated mutations associated with sites of chromosomal DNA breakage/recombination, R-loop accumulation, and extrachromosomal DNA formation^7,30,34,36,37^. Despite the prominent role of APOBEC3 enzymes in cancer mutagenesis, whether and how they interact with other sources of DNA damage remains unexplored.

### Exogenous NQO and endogenous A3B exhibit carcinogenic synergy

Human head & neck cancer is commonly modeled in mice by administering the tobacco carcinogen mimetic 4-nitroquinoline 1-oxide (NQO) in drinking water, which results in visible oral tumors within 16 to 24 weeks^38,39^. Because mutational signatures from tobacco smoke and APOBEC enzymes often occur in the same head & neck and lung tumors^40^, studies here were initiated to test for possible mutagenic interactions between NQO and A3B. This was done by applying the established NQO mutagenesis procedure to wildtype (WT) C57BL/6J animals or to human A3B or A3B-E255A (a catalytically dead mutant) expressing transgenics (workflow in **Fig. 1a**). Human A3B protein levels in these animals (*Rosa26::CAG-L-A3Bi*) approximate those observed in many cancers and result in accelerated tumorigenesis (78-week average penetrance in A3B animals vs 100 in WT animals)^29^. Tissue specimens and sequencing data from these prior experiments are used here for comparison. As anticipated from such long tumor latencies, no oral lesions were evident here at the 32-week timepoint for A3B, A3B-E255A, or WT animals (representative histology, lesion numbers, and images in **Fig. 1b,c,d**).

**Fig. 1.**
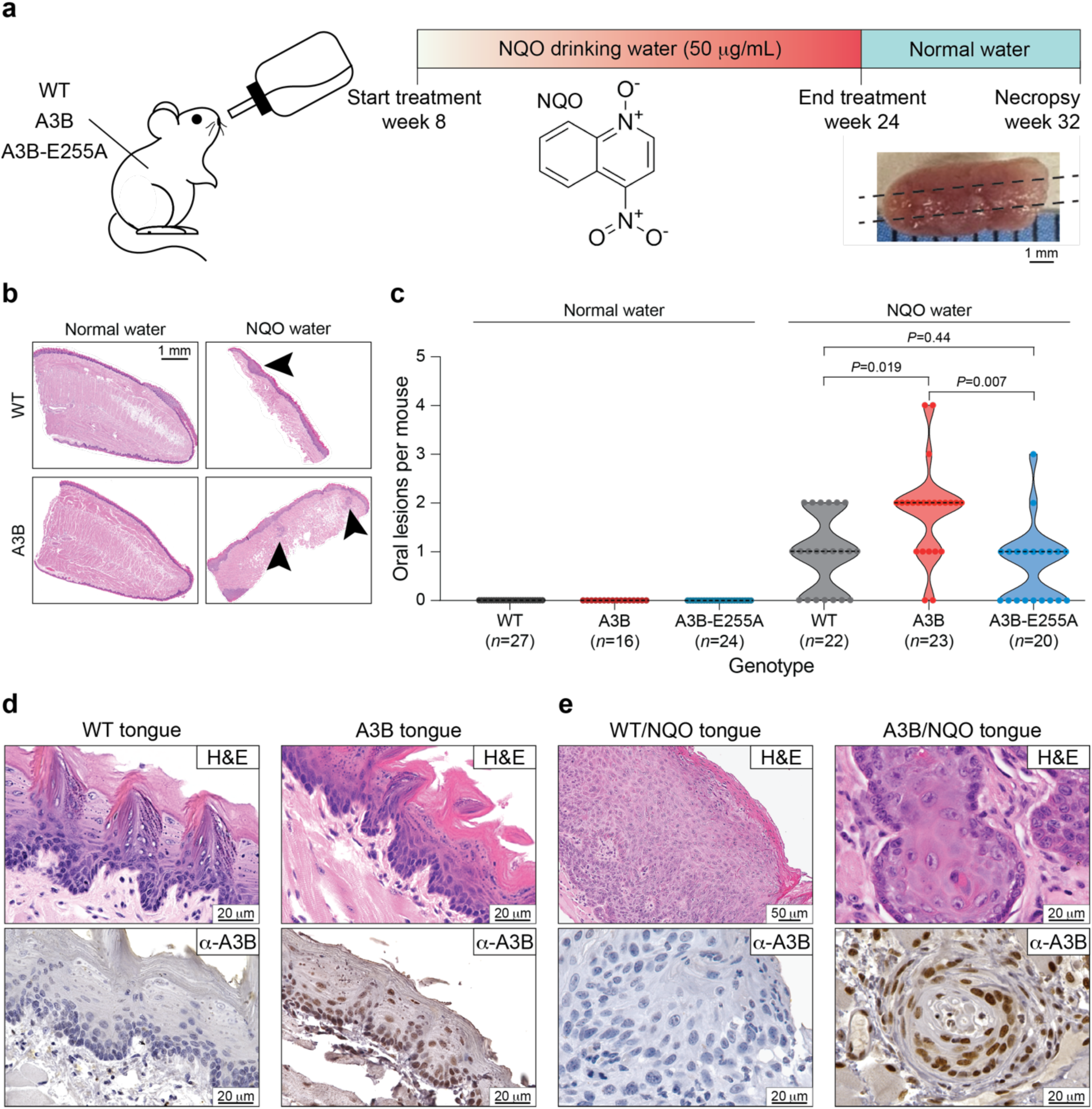
Head & neck tumor formation *in vivo* through A3B and NQO mutagenesis. **(a)** Schematic of the NQO treatment procedure. Animals are treated with NQO for 16 weeks, provided with normal water for 8 weeks, and then sacrificed for analysis including longitudinal sectioning of the tongue and histopathology. **(b)** Representative H&E stained tongue tissues from WT (top) and A3B (bottom) mice after receiving normal water (left) or NQO water (right). The arrowhead in the top right points to an area of epithelial dysplasia, and arrowheads in the bottom right point to invasive SCC (scale=1 mm). **(c)** Quantification of oral lesions in NQO-treated animals (right) in comparison to historic controls provided with normal water (left) with the indicated genotypes. Each dot represents data from an independent animal (dotted lines, median tumor numbers; *P*-values, Poisson regression). (**d**,**e**) H&E (top) and anti-A3B (bottom) stained tongue tissues from WT and A3B mice after receiving normal water (left) or NQO water (right) (scale bars indicated). The lingual surface epithelium in panel-d has no evidence of cytologic atypia. Nuclear A3B staining is strong in basal and spinous cells. Images in panel-e are higher magnifications of lesions from panel-b. The left-side H&E-stained photomicrograph in panel-e demonstrates epithelial dysplastic aberrations, including maturational disorganization, precocious keratinization, and increased nuclear-to-cytoplasmic ratios. The right-side H&E-stained photomicrograph depicts invasive SCC comprising islands and nests of malignant epithelial cells featuring enlarged, hyperchromatic nuclei with macronucleoli infiltrating the fibrous stroma.

In comparison, NQO-treated WT mice exhibit an average of 1 oral mucosal lesion per animal at the 32-week experimental endpoint (representative histology in **Fig. 1b**, lesion numbers in **Fig. 1c**, and representative images in **Fig. 1e**). Oral mucosal lesions are defined here as clinically or microscopically distinct exophytic papillary dysplastic lesions or invasive squamous cell carcinomas (SCCs; combined data from 2 sets of 3 physically separated 4 µm thin sections from each animal’s tongue plus oral cavity; **Methods**). Interestingly, twice as many lesions are evident in the NQO-treated A3B-expressing group, with these animals developing an average of 2 and as many as 4 lesions (representative histology in **Fig. 1b**, lesion numbers in **Fig. 1c**, and representative images in **Fig. 1e**). This synergistic effect requires A3B deaminase activity, because A3B catalytic mutant (E255A)-expressing animals treated with NQO exhibit lesion numbers indistinguishable from NQO-treated WT animals (**Fig. 1c**). A3B catalytic activity also affects the malignant potential of lesions, as evidenced by increased numbers of invasive SCCs in the tongues of A3B/NQO animals but not in A3B-E255A/NQO conditions (**Extended Data Fig. 1a**). However, A3B does not appear to affect individual tumor thickness or the depth of carcinoma invasion (**Extended Data Fig. 1b,c**). Immunohistochemistry (IHC) with a monoclonal antibody that recognizes human A3B^41^ confirms that WT mice lack A3B, whereas animals with both catalytically active and inactive A3B express similar levels of this protein in the nuclear compartment of histologically normal tongue epithelia as well as in oral epithelial dysplastic lesions and invasive SCCs (**Fig. 1d,e**; **Extended Data Fig. 1d,e,f**). Tumors from A3B/NQO mice also manifest elevated γ-H2AX levels indicative of an ongoing DNA damage response (**Extended Data Fig. 1f,g**).

### NQO sensitizes the genome to mutagenesis by A3B

To determine whether the observed tumorigenic synergy is driven by mutagenesis, whole-genome sequencing (WGS) was done for oral tumors from NQO-treated WT, A3B, and A3B-E255A mice alongside matched tails as germline DNA references. This analysis revealed high SBS mutation burdens in tumors from NQO-treated animals (**Fig. 2a**; **Extended Data Fig. 2a,b**; **Supplementary Table 1**). Individual oral tumors show SBS mutation frequencies between 38 and 242 mutations per megabase, which equates to approximately 104,000-659,000 mutations per tumor. These levels are comparable to those observed in highly mutated human tumors^3,42,43^. All NQO-treated tumors contain SBS4 and SBS29 mutations, attributable to this adducting agent and associated in human cancers with tobacco smoking and chewing, respectively^3,14,39^. Most tumors from A3B-expressing animals also show an abundance of C-to-T mutations in TCA and TCT motifs (*i.e*., SBS2). For comparison, tumors isolated from naturally aged WT, A3B, and A3B-E255A animals from our prior studies^29^ do not harbor SBS4 or SBS29 mutations, and only tumors from A3B-expressing animals exhibit SBS2 mutations (**Fig. 2a**; **Extended Data Fig. 2a**; **Supplementary Table 1**).

**Fig. 2.**
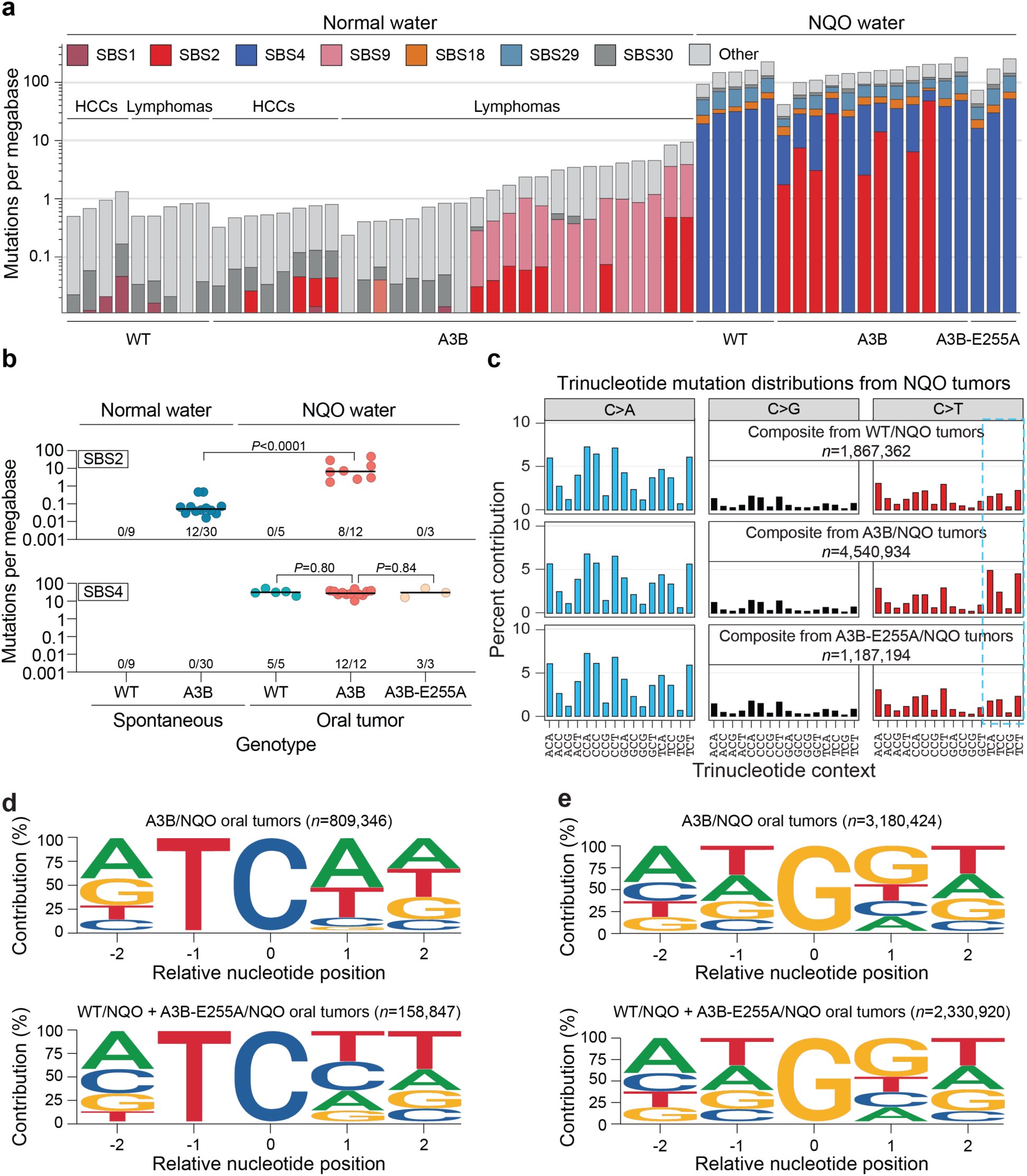
Synergistic increases in APOBEC signature mutations in A3B/NQO tumors. **(a)** Stacked histogram plots of SBS mutation loads in the indicated tumors from historic controls (left) and oral tumors from NQO-treated animals (right). The most prevalent COSMIC SBS mutational signatures are color-coded with APOBEC/SBS2 in red and NQO/SBS4 in blue. The log_10_-scaled Y-axis indicates mutations per megabase. **(b)** APOBEC/SBS2 (top) and NQO/SBS4 (bottom) mutation burdens in individual tumors from animals with the indicated genotypes and treatment conditions. The fractions below each dot plot report the total number of tumors with each mutational signature over the total number of tumors sequenced (horizontal lines, medians; *P*-values, pairwise two-tailed Mann-Whitney U-tests). **(c)** Trinucleotide distributions of all C/G-to-A/T, -G/C, and -T/A mutations in tumors from NQO-treated WT, A3B, and A3B-E255A animals. *n*-values represent the combined total of C/G mutations from each experimental condition. APOBEC signature TC motifs are mutated preferentially in A3B/NQO tumors (blue box). **(d)** Local context of TC-to-TT and -TG mutations in oral tumors from NQO-treated A3B animals (top) in comparison to NQO-treated control animals (WT and A3B-E255A, bottom; *n*=total number of TC context mutations in each group). TC-context mutations in A3B/NQO tumors exhibit a prominent A3B mutation signature with a bias for A or G (purine) at the -2 position and for A or T (W) at the +1 position. **(e)** Local context of G-to-C and G-to-T (G-to-Y) mutations in oral tumors from NQO-treated A3B animals (top) in comparison to NQO-treated control animals (WT and A3B-E255A, bottom; *n*=total number of G-to-Y mutations in each group). The two groups show nearly identical pentanucleotide contexts for NQO-induced G-to-Y mutations with a modest bias for a +1 guanine.

Surprisingly, in NQO oral tumors manifesting an APOBEC mutational signature, the SBS2 mutation burden is an average of ∼100-fold higher than tumors from A3B animals provided with normal drinking water and aged naturally^29^ (SBS2 medians: 6.9 vs 0.05 mutations per megabase; *P*<0.0001 by Mann-Whitney U-test; **Fig. 2b** and summarized in **Supplementary Table 1**). Although this comparison involves historical controls from different tissue types and ages, the markedly elevated SBS2 burden in A3B/NQO tumors is consistent with substantial amplification of APOBEC mutagenesis following NQO exposure.

In contrast, mutational burdens from tobacco-associated signature SBS4 are similarly high in all oral tumors from NQO-treated animals regardless of genotype, indicating that the observed SBS2 mutation increase from A3B constitutes a unique unidirectional synergy (SBS4 medians: 28 vs 32 mutations per megabase for A3B and WT mice, respectively; *P*=0.80 by Mann-Whitney U-test; **Fig. 2b**). In support of this relationship, there is a strong enrichment of C-to-T mutations in A3B-preferred TCW motifs exclusively in A3B/NQO tumors, whereas NQO-induced mutations occur at similar frequencies and local sequence contexts in tumors from all three genotypes (**Fig. 2b,c,d,e**; **Extended Data Fig. 2a,b,c**). NQO-induced G/C-to-T/A transversions have a modest GG dinucleotide bias (**Fig. 2e**). The majority of tumors from A3B/NQO animals also exhibit elevated levels of structural variation and Notch pathway alterations (**Extended Data Fig. 3**), analogous to a subset of human head & neck cancers^42,44^.

Curiously, 4/12 oral SCCs from A3B/NQO animals did not exhibit SBS2. At least one of these is due to a *de novo A3B* minigene mutation causing a somatic E255A substitution and the remainder have yet to be explained. Similar SBS2 penetrance rates were observed in hepatocellular carcinomas and B lymphocyte mouse tumors in our prior studies with naturally aged A3B animals^29^, further suggesting that A3B imposes a selective pressure that can be alleviated, in at least a subset of tumors, by its own mutational inactivation. Therefore, hereafter, our analyses focus on the 8 A3B/NQO tumors that exhibit clear evidence for A3B activity (SBS2).

### *Didyma* - twin APOBEC mutations in A3B/NQO tumors

We next constructed rainfall plots to visualize intermutation distances (IMDs; **Fig. 3a**; **Extended Data Fig. 4a**). A representative oral tumor from an NQO-treated WT animal with 182,789 C-to-T/G mutations (554,429 total SBS mutations) shows that most of these mutations are separated by >1,000 bp with few clustered events. In comparison, an oral tumor from a NQO-treated A3B animal with similar numbers of C-to-T/G mutations (165,128; 331,829 total SBS mutations) also shows predominantly dispersed mutations. However, this A3B/NQO tumor, as well as others from A3B/NQO conditions, also exhibits a striking increase in a new type of paired mutation hereafter called *didyma* (Greek for twins; **Fig. 3a**). We define *didyma* as pairs of strand-coordinated APOBEC signature mutations separated by a short IMD of 1-32nt (yellow dots forming a ladder-like pattern in **Fig. 3a**). APOBEC signature *didyma* comprise an average of 12% of all SBS2 mutations in A3B/NQO tumors and are virtually absent in NQO-treated WT or A3B-E255A tumors, indicating that these mutational events arise from A3B deaminase activity (**Extended Data Fig. 4b,c**). APOBEC signature *didyma* are enriched exclusively in A3B/NQO tumors and, by contrast, opposing-strand APOBEC mutations with the same IMD are not (**Fig. 3b,c**). Interestingly, although APOBEC signature *didyma* occur genome-wide with a slight bias to genic regions in A3B/NQO tumors, they do not appear to associate with transcription or replication (**Supplementary Fig. 1**).

**Fig. 3.**
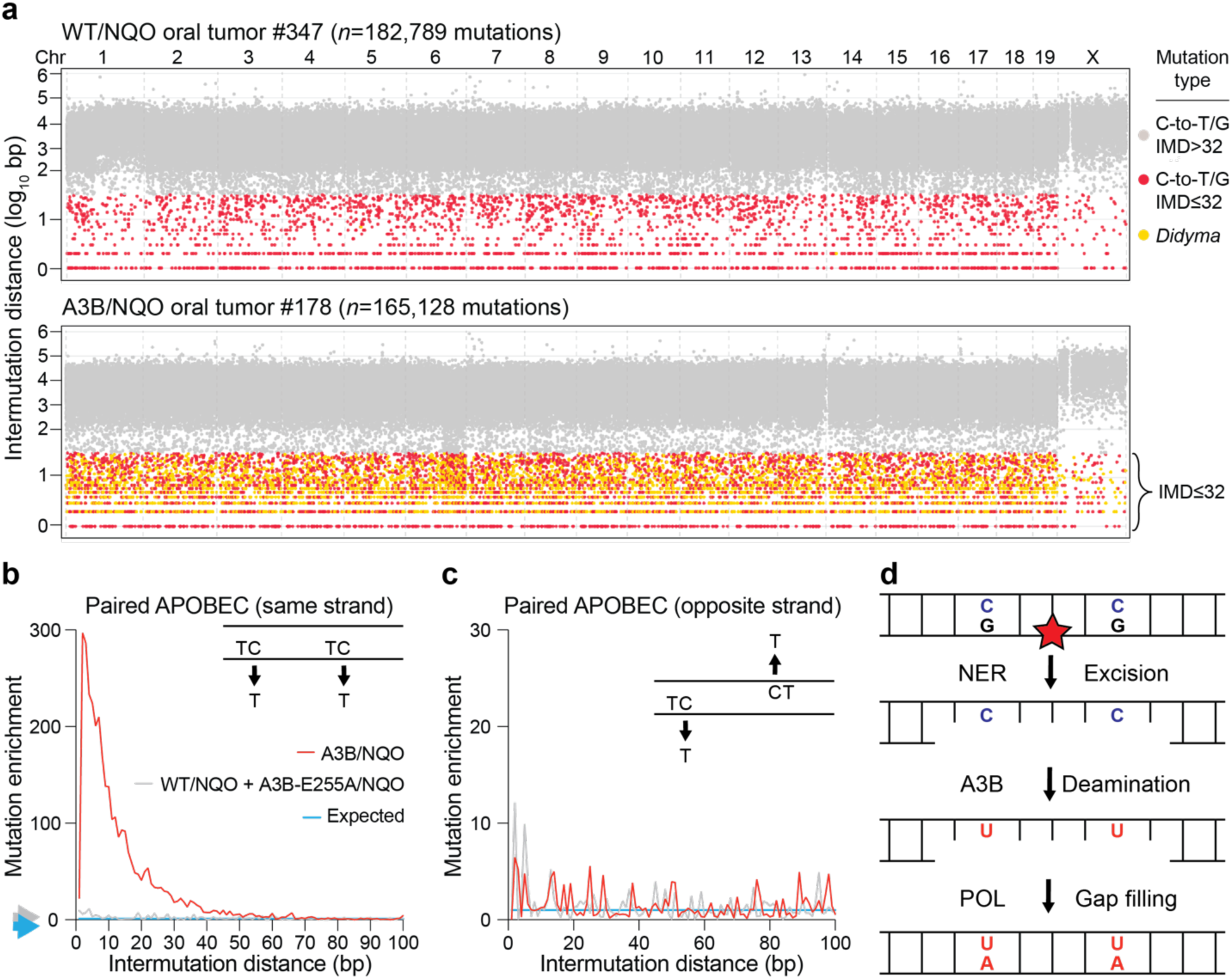
APOBEC *didyma* in tumors from A3B/NQO mice. **(a)** Rainfall plots depicting IMDs for all C-to-T and C-to-G mutations in representative WT/NQO and A3B/NQO tumors. Gray and colored symbols represent mutations with IMD >32bp and ≤32bp, respectively (red = distance between two mutations where one or both is not in an APOBEC-preferred TC motif; yellow = *didyma*, distance between two mutations that are both in APOBEC-preferred TC motifs). *n*-values are total C-to-T/G mutations in each tumor. (**b,c**) Enrichment of twin APOBEC signature mutations in *cis* and *trans*, respectively, in A3B/NQO tumors (*n*=8, red line) vs the control groups combined (*n*=8, gray line). The blue line shows the expected distribution based on simulations. The inset illustrates the queried permutation. **(d)** Schematic of the proposed molecular mechanism. Removal of a DNA adduct such as NQO (red star) by NER creates a 24-32nt ssDNA substrate for C-to-U deamination by A3B. Subsequent gap filling by a DNA polymerase (POL) immortalizes the two U-lesions as APOBEC signature mutations (*didyma*). Downstream uracil repair and/or DNA replication are not shown (expanded model and additional information in **Extended Data** Fig. 7c).

In contrast, although NQO-induced SBS4 mutations are more abundant overall, the distribution of these mutations is similar between conditions (A3B, A3B-E255A, or WT) and they rarely occur in pairs with an IMD <32nt (only ∼3%, **Extended Data Fig. 4d**). In rare instances where SBS4 mutations do occur in pairs, they are not enriched on the same or the opposing DNA strand (**Extended Data Fig. 4e**). NQO-induced G-to-Y mutations are associated with transcription (not replication) and accumulate preferentially on the non-transcribed strand, particularly in highly expressed genes (**Supplementary Fig. 2**), similar to tobacco smoking-associated SBS4 mutations^4,40^. Interestingly, NQO-induced SBS4 mutations are underrepresented on the DNA strand opposite APOBEC signature *didyma* events (**Extended Data Fig. 4f**).

These mutational features, particularly the one-way mutational synergy, the strand-coordinated enrichment of *didyma* within 32bp, and the absence of transcriptional strand asymmetry, combine to suggest a molecular mechanism involving global genome nucleotide excision repair (NER; **Fig. 3d**). In this model, NQO-adducted guanines are removed to yield diagnostic 24-32nt ssDNA tracts, which persist sufficiently long to become substrates for deamination. Subsequent DNA polymerase-mediated gap-filling and strand ligation then immortalize uracil lesions into APOBEC signature mutations (with U’s replaced by T’s anytime thereafter by canonical DNA replication and/or error-free uracil base excision repair). This mechanism is consistent with established NER tract lengths in mammalian cells^45,46^, and studies demonstrating that NQO lesions are preferred substrates for NER^47–50^. Importantly, although *didyma* are a hallmark feature of this mechanism, the short ssDNA tracts created by NER are inevitably also substrates for single A3B-catalyzed C-to-U deamination events, thus explaining the large unidirectional, synergistic, and genome-wide increase in SBS2 in NQO-exposed tumors.

### A3B deaminates cytosines in NER-like ssDNA substrates

To assess whether A3B can act on DNA structures resembling NER intermediates, a series of biochemical experiments was done using purified A3B and defined ssDNA substrates. First, a short ssDNA substrate containing two TCW motifs was used to optimize conditions for single-hit deamination kinetics (**Supplementary Fig. 3a**). Second, A3B processivity was examined using substrates in which two TCW motifs are separated by distances ranging from 5 to 60nt (**Supplementary Fig. 3b**). A3B exhibited comparable processivity across both short (5 to 30nt) and longer distances (up to 60nt), indicating that the unique spacing of *didyma* (<32nt) is unlikely to reflect an intrinsic limitation in enzyme processivity.

Third, to model deamination of NER intermediates, a gapped DNA substrate was designed based on the human *PIK3CA* locus, which contains two APOBEC target motifs and is recurrently mutated in cancer^51–53^ (E542K and E545K; **Fig. 4a,b**). A3B efficiently deaminated each target cytosine individually and also catalyzed dual deamination events within the same substrate, as evidenced by the more-than-additive accumulation of the twin deamination *didyma* product (**Fig. 4b**). Similar results were obtained using an independent gapped substrate (**Supplementary Fig. 3c**).

**Fig. 4.**
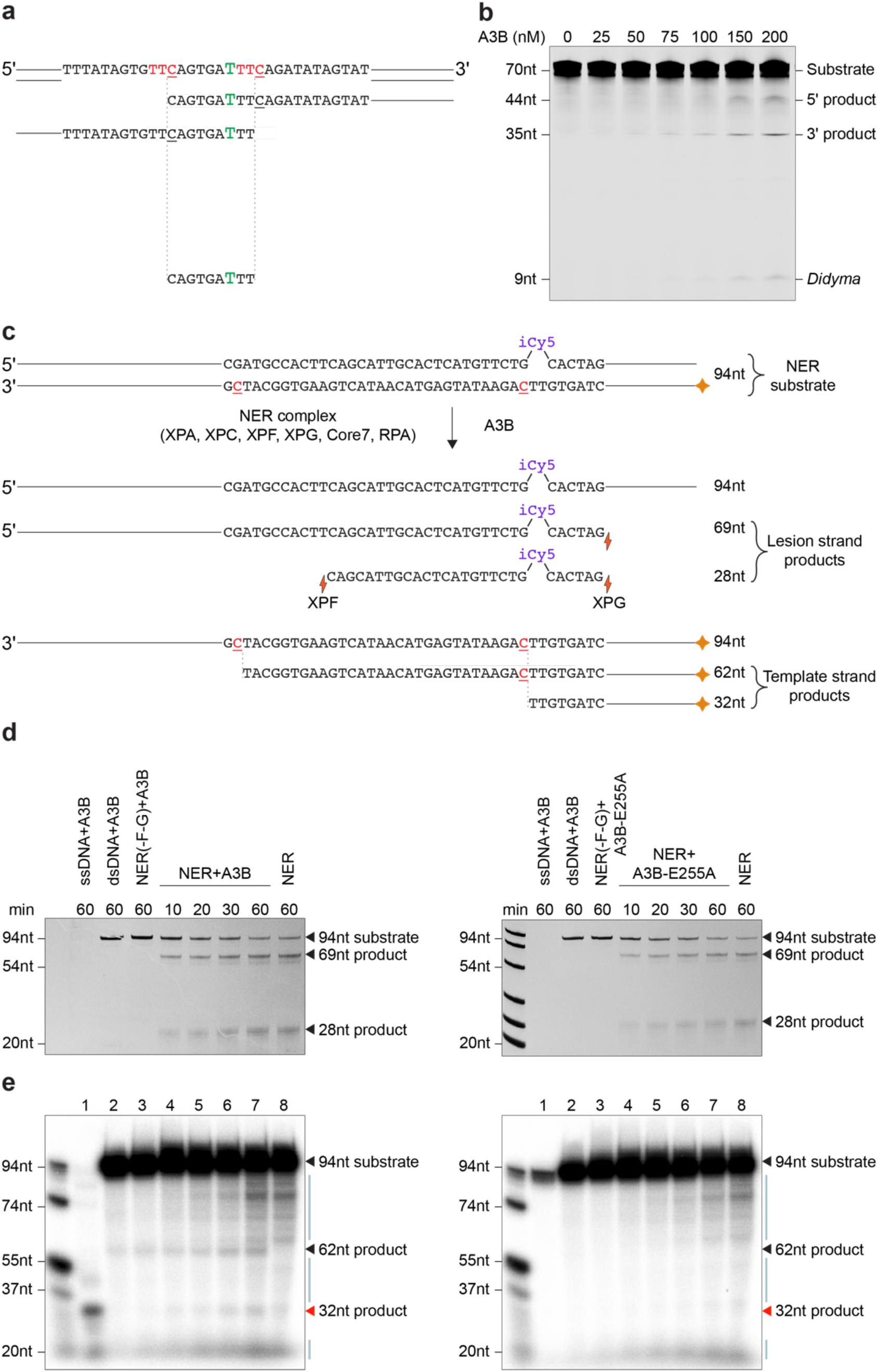
A3B deaminates ssDNA tracts opposite NER-excised damage. **(a)** Schematics of the gapped substrate and three visible deamination products (polarity indicated; green T has a fluorescein label; red bases are *PIK3CA* complementary codons for E542 and E545). **(b)** Gel image of A3B-catalyzed deamination products of the 32nt-gapped substrate depicted in panel-a. **(c)** Schematic of experimental workflow to study A3B deamination during NER. The internal cyanine 5 (iCy5) represents a DNA adduct to initiate NER as well as a fluorescent label for detection. Products mediated by NER and A3B deamination are shown below with nucleotide lengths indicated (orange diamond, ^32^P label for visualization; red underlined C’s are targets for deamination). EndoQ (not shown) cleaves on the 5’ side of dU lesions and liberates the indicated deamination reaction products. (**d**,**e**) Fluoroimages and phosphorimages, respectively, of products following *in vitro* NER and A3B deamination [NER, all NER excision components; NER(-F-G), all NER excision components except XPF and XPG]. The 69nt XPG cleavage product and the ∼28nt dual incision products are labeled. The 32nt deamination product requiring adduct excision by NER is indicated by a red triangle. The 62nt deamination product (black triangle) is also facilitated by NER, likely reflecting substrate breathing or unwinding. The light blue lines to the right of the phosphorimages indicate NER-dependent background cleavage events that accumulate regardless of the presence of A3B (or A3B-E255A; compare lanes 7 and 8).

Fourth, to ask whether A3B can deaminate *bona fide* NER intermediates, a model DNA adduct (Cy5) was incorporated into a DNA duplex substrate with ^32^P-labeled non-lesion strand and incubated *in vitro* with human NER proteins in the presence of A3B or A3B-E255A (**Fig. 4c,d,e**). The Cy5 serves two purposes, as a bulky adduct for NER and as a label for excised strand visualization following denaturing PAGE. EndoQ was included as above to report C-to-U deamination events. As demonstrated^54^, the longer 69nt XPG single incision product and the shorter ∼28nt XPG/XPF dual incision product of the Cy5 strand become evident within 10 min and increase in abundance over time (**Fig. 4d**). This dual-incision reaction is unaffected by inclusion of A3B or A3B-E255A. In comparison, upon phosphorimaging to detect A3B-catalyzed C-to-U deamination events on the non-lesion strand, a 32nt band deamination product of A3B and not A3B-E255A appears with similar kinetics and only in the presence of the full NER excision complex (**Fig. 4e**). Two types of background cleavage events are also evident. The first requires XPG and XPF and is independent of A3B (compare lanes 3 vs 8 in **Fig. 4e**). The second is independent of the NER process: a 62nt deamination product that requires A3B catalytic activity, probably due to substrate breathing. Interestingly, the amount of the 62nt product increases with NER progression, indicating enhanced substrate breathing with the dual incision or unraveling by the NER helicase. Taken together, these results with model gapped DNA substrates and lesion-containing duplex DNA substrates demonstrate that A3B can efficiently deaminate cytosines within ssDNA tracts created by NER.

### XPA and A3B are required for APOBEC signature mutation and *didyma* formation

To test whether NER is required for APOBEC mutagenesis and *didyma* formation in cells, we disrupted *XPA* in human cell lines and the homologous gene, *RAD14*, in yeast. Specifically, we used the human lung adenocarcinoma cell line SW1573^55^, the doxycycline-inducible T-REx-293-A3Bi-eGFP system^56,57^, the U2OS osteosarcoma cell line, and the *Saccharomyces cerevisiae* strain AM7658. Because replication-associated processes can also generate ssDNA with the lagging strand preferentially susceptible to APOBEC deamination^31–33,58^, mutagenesis experiments here were performed under growth-limiting conditions to minimize replication and focus on NER-dependent effects.

*XPA*-null and *rad14*-null cells are hypersensitive to bulky DNA adduct-forming agents such as NQO (**Extended Data Figs. 5 & 6**), consistent with prior reports^59,60^. Here, we uniquely find that this sensitivity is further exacerbated by expression of catalytically active A3B, but not the inactive A3B-E255A mutant, revealing a novel synthetic lethal interaction between A3B activity and unrepaired DNA lesions (**Extended Data Figs. 5 & 6**). NQO sensitivity is also observed in human U2OS cells, which express endogenous A3B protein constitutively (**Extended Data Fig. 5c**). Importantly, *A3B* disruption in this system partly suppresses NQO-sensitivity, indicating that endogenous A3B contributes to the overall synthetic lethal phenotype (**Extended Data Fig. 5c**). In addition, NQO treatment of NHF1 cells triggers an XPA-dependent excision of characteristic fragments ≤32nt (**Supplementary Fig. 3d**), concordant with prior studies with other adducting agents in different cell types^61,62^.

To directly test for NER involvement, parental and *XPA*-null cells expressing A3B, A3B-E255A, or vector control were growth-arrested, exposed to NQO (LD_50_), and analyzed by duplex sequencing (workflow in **Fig. 5a**; SW1573 immunoblots and DNA deaminase activity assay results in **Fig. 5b**). In SW1573 cells, the combination of ectopic A3B expression and NQO treatment yielded 13,954 APOBEC signature SBS2 mutations, and disruption of *XPA* reduced this number by 10-fold (mutation counts are shown normalized per diploid genome in **Fig. 5c**). This dependency is also clearly evident through tri- and penta-nucleotide SBS mutation analyses (**Supplementary Fig. 4a,b,c,d**). Under the same conditions, 686 *didyma* were detected, and *XPA* knockout reduced this number by 95% (*didyma* counts are shown normalized per diploid genome in **Fig. 5d**).

**Fig. 5.**
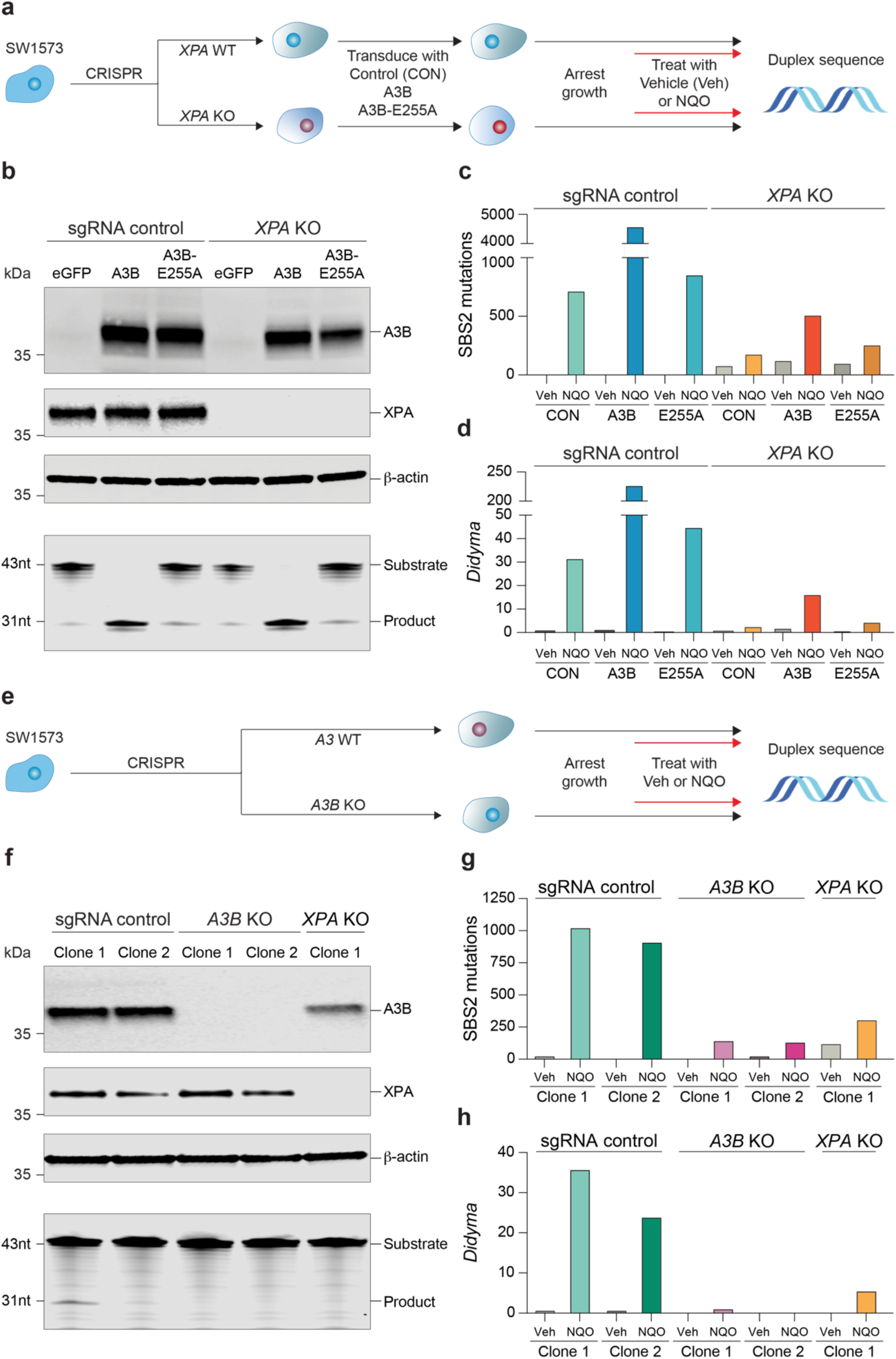
Molecular mechanism of *didyma* formation. **(a)** Schematic of workflow for *XPA* KO experiment. See text and **Methods** for details. **(b)** Immunoblots (top) and ssDNA deaminase activity (bottom) of growth-arrested SW1573 cells from each experimental condition depicted in panel-a. (**c**,**d**) SBS2 mutations and *didyma* per diploid genome in *XPA* WT and KO SW1573 cells from the indicated conditions. **(e)** Schematic of workflow for *A3B* KO experiment. See text for details. **(f)** Immunoblot (top) and ssDNA deaminase activity assay (bottom) of SW1573 and two independent *A3B* KO daughter clones. (**g**,**h**) SBS2 mutation and *didyma* counts per diploid genome in *A3B* WT and KO SW1573 cells from the indicated conditions, including data from a parallel *XPA* KO experiment for comparison.

Notably, both APOBEC signature mutations and *didyma* were also observed in NQO-treated vector control cells, and their occurrence was also dependent upon *XPA* (**Fig. 5c,d**). Given that A3B is the only detectable cancer-associated APOBEC enzyme in SW1573 cells (immunoblots in **Fig. 5b,f** and **Extended Data Fig. 5a**; RTqPCR data in **Extended Data Fig. 5d**), these results indicate that endogenous A3B is sufficient to synergize with NQO through an NER-dependent mechanism. In support of this conclusion, APOBEC mutation pentanucleotide contexts are indistinguishable with or without exogenous A3B expression (**Supplementary Fig. 4a,b**). Similar trends were observed with a lower concentration of NQO (LD_10_; **Extended Data Fig. 5e,f**). APOBEC signature *didyma* mutations in SW1573 showed a non-transcribed strand (NTS) bias and no association with DNA replication (**Supplementary Fig. 4e,f**). CRISPR knockout of endogenous *A3B* in SW1573 cells confirmed that this enzyme is required for formation of nearly all NQO-induced APOBEC signature SBS2 mutations and *didyma* (**Fig. 5e,f,g**; **Supplementary Fig. 4c,d**).

Comparable results were obtained in yeast, where human A3B expression combined with NQO exposure resulted in an average of 311 APOBEC signature mutations and 16 *didyma*, both of which required *RAD14* (values normalized per haploid genome in **Extended Data Fig. 6**). These findings are consistent with the conserved generation of short ssDNA excision intermediates during NER^46^. Taken together, the human cancer cell line SW1573 and yeast mutation results demonstrate that both NER and A3B are involved in the formation of synergistic levels of APOBEC mutagenesis and *didyma*, supporting a model in which the repair of bulky DNA adducts generates ssDNA intermediates that can be deaminated by A3B (**Fig. 3d**).

### APOBEC signature mutations and *didyma* are elevated in tumors from smokers

Because tobacco smoke contains multiple DNA adduct-forming carcinogens^2^, a key prediction from our A3B/NQO experiments is that APOBEC signature mutations should be elevated in smokers compared to non-smokers (S and NS, respectively). To test this possibility, we analyzed a large dataset comprising 1,498 lung adenocarcinomas and squamous cell carcinomas sourced from The Cancer Genome Atlas (TCGA), Pan-Cancer Analysis of Whole Genomes (PCAWG), and other publicly available datasets^3,63–67^. Sequencing data from 265 head & neck tumors^44^ were also analyzed. First, we found that the burden of APOBEC signature mutations is nearly double in lung cancers from smokers compared to non-smokers (median: 2.3 vs 1.2 mutations per megabase, respectively; *P*<0.0001 by Mann-Whitney U-test; **Fig. 6a**). This equates to a median of approximately 6,440 APOBEC signature mutations in tumors from smokers versus 3,360 in tumors from non-smokers. Moreover, a strong positive association is evident between the number of mutational events attributable to APOBEC (SBS2+SBS13) and those attributable to smoking (SBS4; r=0.61, 95% CI=0.54-0.67, *P*<0.0001, Spearman’s correlation; **Fig. 6b**). In contrast, no meaningful association is observed between SBS2/13 and aging-associated SBS1, which is found in nearly all lung cancers (**Fig. 6c**).

**Fig. 6.**
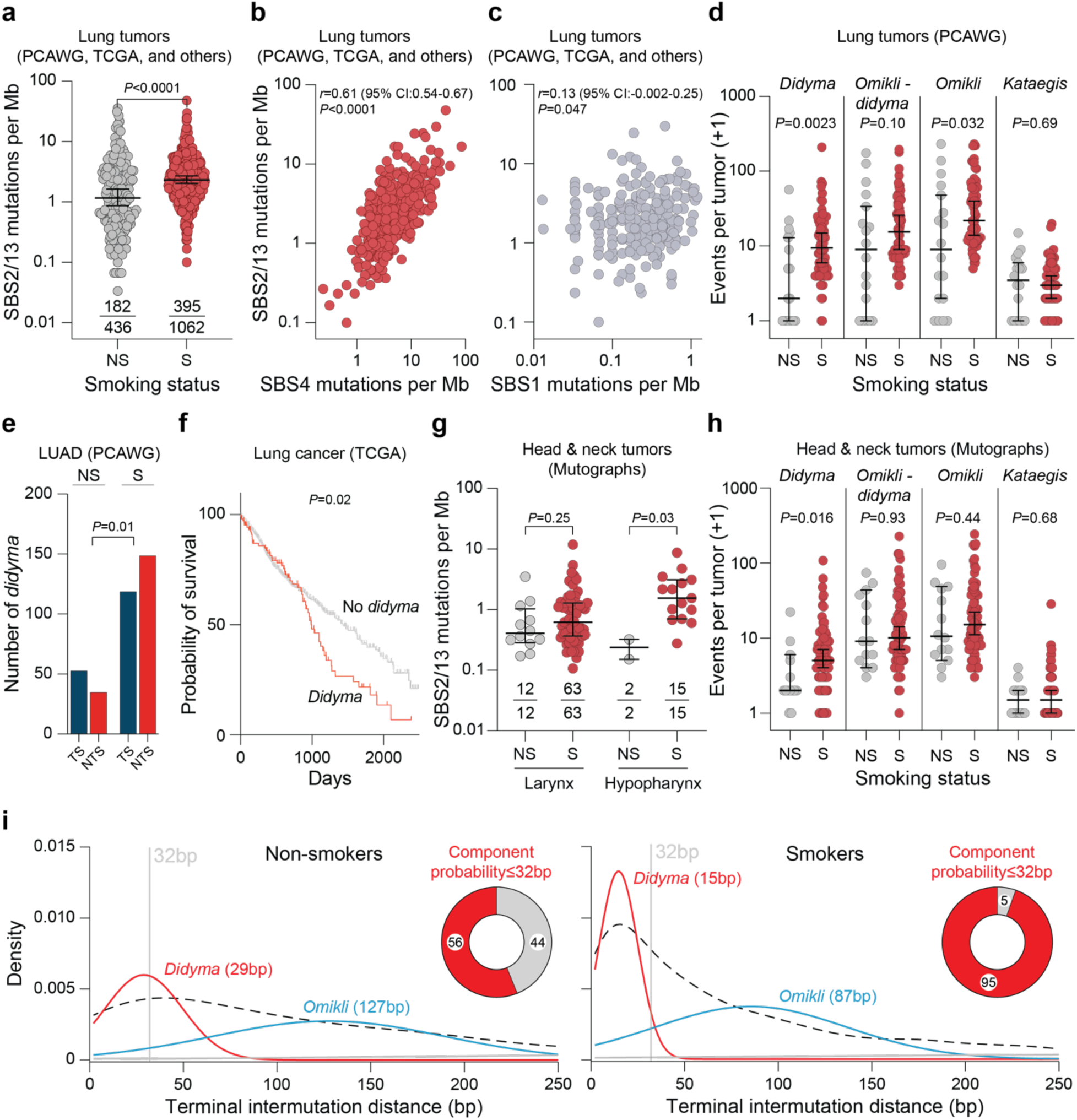
Elevated APOBEC signature mutations and *didyma* in tumors from smokers. **(a)** Quantification of APOBEC signature mutation loads (SBS2/13) in lung adenocarcinomas and SCCs from non-smokers (NS) and smokers (S). Fractions report the total number of tumors with an APOBEC mutational signature over the total number analyzed. The horizontal lines and whiskers represent medians and 95% confidence intervals (*P*-value, two-tailed Mann-Whitney U-test). (**b**,**c**) Scatter plot indicating a direct relationship between APOBEC SBS2/13 mutations and tobacco smoking-associated SBS4 mutations (*n*=395 tumors with both signatures), in comparison to no association between SBS2/13 and SBS1 mutations (*n*=234 tumors with both signatures; *p*- and *r*-values, Spearman’s correlation). **(d)** Quantification of the indicated cluster mutation types in lung tumors from smokers and non-smokers (*n*=66 and *n*=18 tumors, respectively). The horizontal lines and whiskers represent medians and 95% confidence intervals, respectively (*P*-values, pairwise two-tailed Mann-Whitney U-tests). **(e)** Quantification of *didyma* in the indicated transcriptional regions in PCAWG lung adenocarcinomas (LUAD) from smokers and non-smokers (*n*=21 and 8 tumors, respectively; TS, transcribed strand; NTS, non-transcribed strand; *P*-value, two-sided Fisher’s exact test comparing observed mutations in smokers and non-smokers). **(f)** Overall survival of lung cancer patients whose tumors exhibit *didyma* (red, *n*=131 with ≥1 *didyma*) or not (gray, *n*=575 with no *didyma*). Data from TCGA lung adenocarcinomas and lung SCCs (*P*-value, log-rank test). **(g)** Quantification of APOBEC signature mutation loads in head & neck larynx and hypopharynx tumors from non-smokers and smokers. The horizontal lines and whiskers represent medians and 95% confidence intervals (*P*-values, two-tailed Mann-Whitney U-tests). Fractions report the total number of tumors with an APOBEC mutational signature over the total number analyzed. **(h)** Quantification of the indicated cluster mutation types in head & neck larynx and hypopharynx tumors from smokers and non-smokers (*n*=78 and *n*=14 tumors, respectively). The horizontal lines and whiskers represent medians and 95% confidence intervals (*P*-values, pairwise two-tailed Mann-Whitney U-tests). **(i)** Gaussian mixture model of *didyma* and *omikli* distance distributions in non-smokers and smokers using a 3-component-based model, representing terminal IMDs up to 250bp. The 3 components (red, blue, and gray) are overlaid on top of the observed distribution of all *didyma* and *omikli* combined (black dashed line) from the lung and head & neck cancers in panels-d and -h. Inset pie charts represent the probability of clustered mutations within the shortest component having a width ≤32bp. Components are labeled with inferred clustered mutational event and peak value.

We next asked whether tobacco smoke carcinogens are associated with APOBEC mutation clustering, including *didyma*, in lung cancer. This analysis revealed large increases in *didyma* in smokers compared to non-smokers (median: 9 vs 1 *didyma* per tumor, respectively; *P*=0.0023 by Mann-Whitney U-test; **Fig. 6d**). Further consistent with tobacco smoke carcinogens triggering a NER-dependent opening of DNA for *didyma* formation, a non-transcribed strand *didyma* bias is observed exclusively in tumors from smokers (**Fig. 6e**). Moreover, *didyma*-positive lung tumors were associated with poorer overall survival, suggesting potential clinical relevance that warrants further investigation (**Fig. 6f**).

There is also a moderate increase in *omikli*, but not in *kataegis*, in lung tumors (**Fig. 6d**). One explanation is that a subset of *didyma* may have been misclassified previously as *omikli*, which were defined originally as 2-3 clustered APOBEC mutations associated with mismatch repair^36^. This interpretation is supported by sub-analyses where *didyma* are subtracted from *omikli*, with the significance of the latter mutational events diminishing substantially (compare results and statistics for *omikli* versus *omikli* minus *didyma* in **Fig. 6d**). In contrast, mechanistically distinct *kataegis* events occur at similar levels in tumors from both smokers and non-smokers (**Fig. 6d**).

In head & neck tumors, a 7-fold increase in APOBEC signature mutations is apparent in hypopharyngeal tumors from smokers versus non-smokers but not in other tumor subtypes of the head & neck (median: 1.5 vs 0.2 mutations per megabase for SBS2+SBS13, respectively; *P*=0.03 by Mann-Whitney U-test; **Fig. 6g**). Furthermore, elevated *didyma* levels are also evident in tumors from the hypopharynx and larynx of smokers (median: 4 vs 1 *didyma* per sample, respectively; *P*=0.016 by Mann-Whitney U-test), whereas enrichment is not observed for *kataegis* or *omikli* (**Fig. 6h**). The observed synergy might be restricted to certain head & neck tumor sites because, like lung tissue, the hypopharynx is exposed for longer periods of time to mutagens from cigarette smoke after each inhalation^44^. *Didyma* comprise both C-to-T transitions and C-to-G transversions at ratios similar to those observed for single, non-clustered APOBEC signature mutations in lung and head & neck human tumors (**Extended Data Fig. 7a,b**). This result indicates that at least a subset of the excision tract deamination events are processed by uracil DNA glycosylase into abasic sites, which are key intermediates in the formation of C-to-G SBS13 mutations^18^ (expanded model in **Extended Data Fig. 7c**).

### Topology of APOBEC mutation clusters in human cancers

To further resolve the structure of APOBEC mutation clusters, we applied probabilistic models (Gaussian and gamma mixture) to assess cluster width (distance between terminal mutations) and IMDs, respectively, to deconvolute the mutational processes driving clusters with 2-3 APOBEC signature mutations (**Methods**). Decomposing the event width distributions from lung and head & neck tumors of smokers suggests the occurrence of three distinct mutational processes: one generating shorter clusters (up to ∼32bp, *didyma*), one intermediate clusters (up to ∼170bp, *omikli*), and one longer clusters (up to ∼680bp, termed “diffuse *omikli*”) (**Fig. 6i** and **Supplementary Fig. 5a**). Moreover, both probabilistic models suggest that cluster widths and IMDs are shorter in smokers than in non-smokers, with one component confined largely to ≤32bp in tumor specimens from smokers, further consistent with the mechanism described here for *didyma* formation (**Fig. 6i** and **Supplementary Fig. 5a,b,c**). 95% of the shortest component mutations have an IMD ≤32bp in smokers, in comparison to 56% in non-smokers, suggesting a potential overlap in the classification of *didyma* and *omikli* predominantly in the latter group (**Fig. 6i** and **Supplementary Fig. 5a,c**). Expanding the Gaussian mixture model to include the 6 cancer types with the most *didyma* shows that these mutation patterns are consistent across cancers (**Supplementary Fig. 5d**)

### Mutational synergy can precede tumor formation

To determine whether this mutational synergy arises prior to tumorigenesis, we analyzed WGS data from 632 normal human bronchial epithelial cell colonies representing 16 independent subjects^68^. Non-neoplastic lung cells from smokers showed nearly 3-fold higher burdens of dispersed APOBEC signature mutation events (median of 0.2 and 0.07 mutations per Mb, respectively; *P*<0.0001 by Mann-Whitney U-test), as well as a significant increase in *didyma* (*P*=0.0007 by Mann-Whitney U-test; **Extended Data Fig. 8a,b**). Moreover, exposing A3B-expressing mice to NQO drinking water for 8 weeks triggered *didyma* formation in the tongue without visible tumor formation, with no significant increases in *omikli* or *kataegis* (**Extended Data Fig. 8c**). Thus, tobacco smoke mutagens and APOBEC3 enzymes are also capable of combining synergistically in pre-cancer cells.

### Pan-cancer landscape of APOBEC signature mutations and *didyma*

A key prediction of the NER-dependent model proposed here is that *didyma* formation occurs in tumors with active APOBEC enzymes and concurrent exposure to DNA adduct-forming agents in addition to those in tobacco smoke. To address the breadth of these coincident processes, we analyzed WGS data from 26 dìerent cancer types^65,69,70^. As a reference, APOBEC signature-positive lung adenocarcinomas from smokers exhibit >2-fold more *didyma* than non-smokers (medians of 19.5 vs 8, **Fig. 7a**, top panel). Interestingly, localized and metastatic bladder tumors have even more *didyma* (medians of 22 and 32, respectively) despite having similarly high levels of APOBEC signature mutagenesis (**Fig. 7a**, top and middle panels). Although bladder cancer is associated with tobacco smoking, it typically lacks the canonical tobacco-associated SBS4 signature^66,71^. These observations raise the possibility that tissue-specific metabolic processing of tobacco-related compounds may trigger distinct mutational processes in the bladder, such as the mechanism responsible for SBS92^66^, which may also sensitize tumor cells to APOBEC mutagenesis as described here. Another distinct mechanism may be at play in cervical cancer, where neither SBS4 nor SBS92 nor any other known tobacco-associated signature is observed^66^, despite observations of tobacco-derived metabolites occurring in cervical mucus^72^. Consistent with NER driving APOBEC mutagenesis, *didyma* occur more frequently on the non-transcribed strand across cancer types with the highest *didyma* levels (**Fig. 7b** and **Supplementary Fig. 6**; also see **Fig. 6e** above for lung cancer data).

**Fig. 7.**
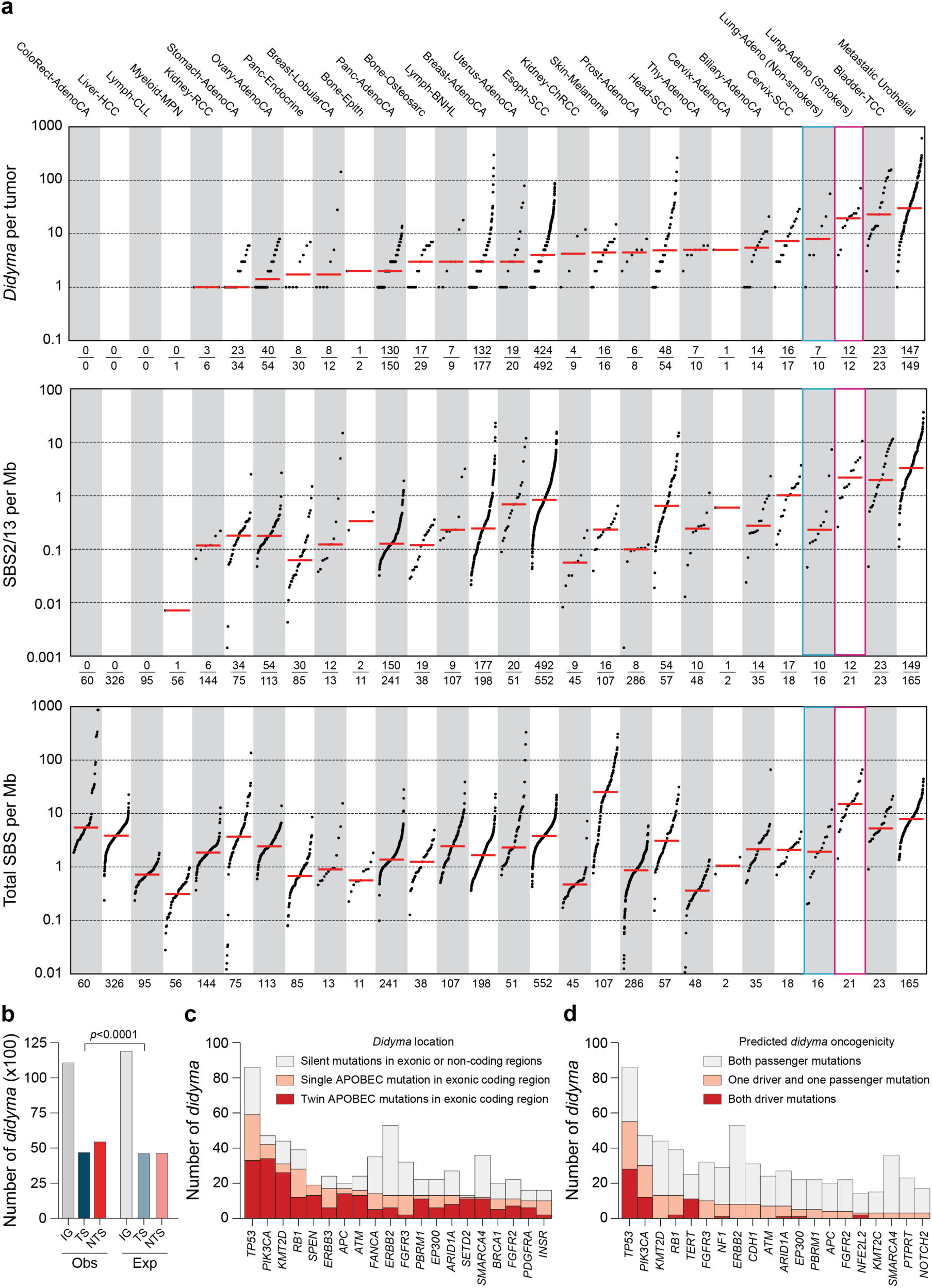
Pan-cancer analysis of APOBEC signature mutations and *didyma*. **(a)** *Didyma* counts, SBS2/13 mutation numbers, and total SBS mutation loads in the indicated tumor types. Horizontal red lines represent median values. Fractions show the number of tumors with *didyma* over the total number of tumors with SBS2/13 mutations, the number of tumors with SBS2/13 mutations over the total number of tumors analyzed, and the total number of tumors of each type. One SBS2/13 data point below 0.001 SBS per Mb is not shown. Data from Fig. 6 lung adenocarcinomas from non-smokers (blue box) and smokers (pink box) are highlighted here for comparison. **(b)** Quantification of aggregated *didyma* mutations in the indicated regions in the top 6 cancer cohorts with the most *didyma* (*n*=281 tumors; IG, intergenic; TS, transcribed strand; NTS, non-transcribed strand). Observed mutations are from tumor data, and expected are from simulations (*P*-value, two-sided Fisher’s exact test). (**c,d**) The top 20 genes with *didyma* in the MSK-IMPACT sequencing panel, ranked by highest to lowest number of coding region changes or predicted driver mutations (clinical annotation by OncoKB).

Skin melanomas, which typically have genomes dominated by mutations associated with unrepaired pyrimidine dimers caused by UV light, also have a significant number of APOBEC signature *didyma* (**Fig. 7a**). To avoid overlap between C-to-T transitions from UV light and APOBEC mutagenesis, we performed a pan-cancer analysis focusing on APOBEC signature SBS13 C-to-G *didyma*, which indicates that a subset of melanomas exhibit *bona fide* APOBEC mutagenesis and, moreover, that UV-triggered NER may sensitize to APOBEC mutagenesis and *didyma* formation (**Extended Data Fig. 9a**). Overall, SBS13 C-to-G *didyma* are apparent nearly all of the same tumor types as SBS2 C-to-T *didyma*, consistent with a common underlying molecular mechanism (**Extended Data Fig. 7c**).

Many individual tumors across other cancer types also exhibit significant numbers of *didyma* (**Fig. 7a**; **Extended Data Fig. 9a**). For instance, the distribution of *didyma* in breast cancer resembles those of esophageal and head & neck cancers, with a significant percentage of individual tumors manifesting high amounts and others low amounts. These results suggest that a subset of these cancer types is exposed simultaneously to both active APOBEC3 deamination and at least one DNA adducting agent. Regardless of tumor type and adducting agent (established or inferred), APOBEC signature mutations within *didyma* invariably occur on the same DNA strand (*i.e*., such rare events are likely NER-dependent and due to processive APOBEC3 activity given the short tract length but could be mistakenly annotated as *omikli*; **Extended Data Fig. 9b,c**). It is also important to note that adult tumor types exhibit a wide range of SBS mutation loads (**Fig. 7a**, bottom panel; also see prior reports^3,73,74^) and that, as evidenced by colorectal tumors with some of the highest SBS mutation burdens, high mutation frequencies alone are insufficient for *didyma* formation.

### *Didyma* target cancer genes and contribute to driver mutations

To begin to gauge the functional relevance of *didyma*, we investigated their occurrence in established cancer genes by leveraging a pan-cancer cohort of 46,073 patients, previously profiled using the FDA-authorized MSK-IMPACT targeted sequencing platform^75^ (**Methods**). These analyses confirmed the human WGS findings above across many cancer types, including lung and head & neck tumors, manifesting APOBEC SBS2/13 signature mutations, coincident SBS2/13 and SBS4 mutation signatures, and APOBEC signature *didyma* (**Extended Data Fig. 10a,b,c**). Numerous cancer driver genes exhibit *didyma* with at least one mutation, and often both, affecting the coding sequence (**Fig. 7c**). Importantly, *didyma* occur in established cancer genes, including *TP53* and *PIK3CA*, and can be comprised of mutations previously annotated as oncogenic drivers (**Fig. 7d**). For instance, one or both mutations of the majority of *TP53* and *PIK3CA didyma* have been reported as driver events^53,76^ (**Extended Data Fig. 10d,e**). PI3K E542K and E545K substitutions are common in breast and head & neck cancers and sometimes part of the same *didyma* event (**Extended Data Fig. 10f**).

## DISCUSSION

DNA damage and mutation processes are generally considered independent, with their effects accumulating additively over time^8,15,16^. Studies here were initiated with mice as a biological *tabula rasa* to investigate oral tumorigenesis driven by mutations from the endogenous mutagen, human A3B, and the exogenous tobacco surrogate, NQO. These *in vivo* experiments revealed an unexpected unidirectional synergy between these very distinct mutagens. Specifically, A3B-inflicted SBS2 mutations increased synergistically in NQO-treated animals, whereas the frequencies of NQO-induced SBS4 mutations remained unchanged regardless of A3B status. This one-way mutational synergy requires A3B deaminase activity, as evidenced by a near-complete absence of SBS2 mutations in otherwise isogenic animals expressing a catalytically inactive protein (A3B-E255A). WGS further revealed unprecedented numbers of APOBEC signature mutations in murine tumors, as well as unique pairs of A3B signature mutations with remarkably short intermutational distances (<32nt). These pairs of APOBEC signature mutations are reminiscent of twins and here named *didyma*. A3B signature *didyma* are strand-coordinated and consistent with a model in which bulky lesions, such as NQO-guanine adducts, are excised by NER, and the resulting exposed single-stranded DNA tract is deaminated by A3B before being converted back into a protected duplex by DNA polymerase-mediated gap-filling (minimal model in **Fig. 3d** and detailed version in **Extended Data Fig. 7c**). This molecular mechanism is supported by biochemical and genetic studies here including, importantly, a requirement of endogenous A3B in human cells and a requirement for the conserved NER initiation factor XPA in human cells and Rad14 in yeast.

### *Didyma* are distinct from *omikli* and *kataegis*

*Didyma* are DNA adduct- and NER-dependent twin APOBEC signature mutations with short (<32nt) IMDs that most likely occur during processive single enzyme-substrate encounters. Although *omikli* and *kataegis* may also arise from single enzyme-substrate encounters, *omikli* tracts are typically longer and associated with mismatch repair^36^, whereas *kataegis* tracts are the longest and associated with recombination repair, R-loop homeostasis, and extrachromosomal DNA pathology^30,34,37^. Although mechanistically distinct, these three mutational processes are challenging to classify through individual DNA sequences. For instance, a *didyma* tract may incur 3 APOBEC signature mutations (*e.g*., **Extended Data Fig. 9b**) and a mismatch repair tract may incur 2, 3, or many more APOBEC signature mutations (*e.g*., ref.^77^). However, when analyzed globally using probabilistic modeling, distinct peaks emerge with the shortest IMD representing APOBEC signature *didyma* (2 strand-coordinated mutations up to ∼32nt), the intermediate *omikli* (2-3 mutations up to ∼200nt), and the longest diffuse *omikli* (2-3 mutations up to ∼800nt; *omikli* described originally in ref.^36^ with distributions refined here; **Fig. 6i**, **Supplementary Fig. 5**, and **Methods**).

Studies here with NQO treatment of human A3B-expressing mice, human cancer cell lines, and yeast reveal a strong synergy at the pathological level through tumor formation and at the molecular level through elevated APOBEC signature mutation and *didyma* formation. This is a striking one-way synergy in which A3B SBS2 mutational events increase synergistically but NQO SBS4 events do not. *Didyma* with SBS13 mutations are not observed in some systems (including the A3B/NQO murine model here) but are clearly evident in human tumors, suggesting differential roles or accessibility of UNG2, REV1, and/or other DNA repair factors (**Extended Data Fig. 7c**). Analogous experiments have yet to be done with other cancer-associated APOBEC family members including A3A, APOBEC1, and AID. However, given the processive nature of these enzymes^78,79^, exclusive preferences for ssDNA substrates^27,30,78^, and the high prevalence of APOBEC signature mutations in the majority of human cancer types^3,6,22,80^ (**Fig. 7a**, middle panel), it is likely that the example detailed here with A3B will constitute the first of many mutational synergies between human DNA deaminases and a wide variety of DNA adducting agents. Indeed, because many human tumors exhibit high levels of APOBEC signature mutations ascribed to A3A and A3B^17–19^, it is likely that A3A will be able to similarly synergize with adducting agents and promote *didyma* formation.

Our findings provide new mechanistic insights into the mutational landscapes of many lung and head & neck cancers. First, we find strongly elevated levels of APOBEC signature SBS2 and SBS13 mutations in smokers compared to non-smokers (**Fig. 6a,g**). This difference was noted but not explained mechanistically in prior studies^4,40^. Second, here we uniquely uncover a strong positive association between APOBEC mutation loads (SBS2/13) and smoking-associated mutation loads (SBS4) in whole-genome and whole-exome sequenced lung cancers from TCGA, PCAWG, and other publicly available sources^63–66^ (**Fig. 6b**). Third, we demonstrate that APOBEC signature SBS2 and SBS13 mutations manifest as *didyma* in lung tumors from smokers more frequently than in non-smokers (**Fig. 6d,h**). Because NER is a universally conserved DNA repair mechanism^61,81,82^, these hallmark *didyma* are likely due to a mechanism in which two processive APOBEC deamination events are inflicted in the 24-32nt ssDNA tract created by NER prior to gap-filling DNA synthesis and ligation (core model in **Fig. 3d** and detailed model in **Extended Data Fig. 7c**). Although the global genomic NER pathway is undoubtedly involved, as *didyma* are evident genome-wide including in many non-expressed regions, the relative contribution of the transcription-coupled NER pathway is less clear as *didyma* are only biased to the non-transcribed strand in the human cell line SW1573 and a subset of human tumor types (**Fig. 6e**, **Fig. 7b**, and **Supplementary Fig. 4e**). Fourth, our studies also show that smoking is strongly associated with APOBEC *didyma* formation, whereas it has little effect on *omikli* and no effect on *kataegis* (**Fig. 6d,h**). Fifth, head & neck cancer data are used to show that APOBEC signature mutations and *didyma* are exacerbated in smokers in comparison to non-smokers (**Fig. 6g,h**). Sixth, we discover similar APOBEC mutation signature and *didyma* enrichments in phenotypically normal lung bronchiolar specimens from smokers in comparison to non-smokers, as well as in A3B/NQO-exposed normal murine oral tissues, indicating that this one-way mutational synergy can occur in normal tissues prior to tumor formation (**Extended Data Fig. 8a,b,c**).

Pan-cancer analyses show that *didyma* can also occur in significant subsets of many different cancer types, most notably bladder cancer, where SBS4 (smoking signature) is rare, strongly suggesting the existence of additional classes of adducting agents that sensitize genomes to APOBEC mutagenesis (**Fig. 7a**). Given the fact that platinum-based chemotherapies are standard-of-care for bladder tumors^83^ and intra-strand platinum adducts are established NER substrates^84,85^, it is possible that the increased numbers of APOBEC mutations and *didyma* in metastatic urothelial cancer may be triggered in part by the chemotherapy itself. Last, but not least, given the breadth of APOBEC mutagenesis in cancer^3,6,22,80^ (**Fig. 7a**), our findings suggest that NER-dependent amplification of APOBEC mutagenesis may extend to additional tissues and DNA adduct-forming agents, providing a framework for identifying previously unrecognized synergistic interactions between mutational processes.

## REFERENCES

1 Sasco, A. J., Secretan, M. B. & Straif, K. Tobacco smoking and cancer: a brief review of recent epidemiological evidence. Lung Cancer 45 **Suppl 2**, S3–9 (2004). 10.1016/j.lungcan.2004.07.998

2 Hecht, S. S. & Hatsukami, D. K. Smokeless tobacco and cigarette smoking: chemical mechanisms and cancer prevention. Nature Reviews 22, 143–155 (2022). 10.1038/s41568-021-00423-4

3 Alexandrov, L. B. et al. The repertoire of mutational signatures in human cancer. Nature 578, 94–101 (2020). 10.1038/s41586-020-1943-3

4 Zhang, T. et al. APOBEC affects tumor evolution and age at onset of lung cancer in smokers. Nat Commun 16, 4711 (2025). 10.1038/s41467-025-59923-8

5 Sondka, Z. et al. COSMIC: a curated database of somatic variants and clinical data for cancer. Nucleic Acids Res 52, D1210–D1217 (2024). 10.1093/nar/gkad986

6 Alexandrov, L. B. et al. Signatures of mutational processes in human cancer. Nature 500, 415–421 (2013). 10.1038/nature12477

7 Nik-Zainal, S. et al. Mutational processes molding the genomes of 21 breast cancers. Cell 149, 979–993 (2012). S0092-8674(12)00528-4

8 Koh, G., Degasperi, A., Zou, X., Momen, S. & Nik-Zainal, S. Mutational signatures: emerging concepts, caveats and clinical applications. Nature Reviews 21, 619–637 (2021). 10.1038/s41568-021-00377-7

9 Shibutani, S., Takeshita, M. & Grollman, A. P. Insertion of specific bases during DNA synthesis past the oxidation-damaged base 8-oxodG. Nature 349, 431–434 (1991). 10.1038/349431a0

10 Zamborszky, J. et al. Loss of BRCA1 or BRCA2 markedly increases the rate of base substitution mutagenesis and has distinct effects on genomic deletions. Oncogene 36, 746–755 (2017). 10.1038/onc.2016.243

11 Setton, J. et al. Long-molecule scars of backup DNA repair in BRCA1- and BRCA2-deficient cancers. Nature 621, 129–137 (2023). 10.1038/s41586-023-06461-2

12 Peterson, L. A. Formation, repair, and genotoxic properties of bulky DNA adducts formed from tobacco-specific nitrosamines. J Nucleic Acids 2010 (2010). 10.4061/2010/284935

13 Kucab, J. E. et al. A compendium of mutational signatures of environmental agents. Cell 177, 821–836 e816 (2019). 10.1016/j.cell.2019.03.001

14 Nik-Zainal, S. et al. The genome as a record of environmental exposure. Mutagenesis 30, 763–770 (2015). 10.1093/mutage/gev073

15 Tomasetti, C., Li, L. & Vogelstein, B. Stem cell divisions, somatic mutations, cancer etiology, and cancer prevention. Science 355, 1330–1334 (2017). 10.1126/science.aaf9011

16 Lee-Six, H. et al. The landscape of somatic mutation in normal colorectal epithelial cells. Nature 574, 532–537 (2019). 10.1038/s41586-019-1672-7

17 Carpenter, M. A. et al. Mutational impact of APOBEC3A and APOBEC3B in a human cell line and comparisons to breast cancer. PLoS Genet 19, e1011043 (2023). 10.1371/journal.pgen.1011043

18 Petljak, M. et al. Mechanisms of APOBEC3 mutagenesis in human cancer cells. Nature 607, 799–807 (2022). 10.1038/s41586-022-04972-y

19 DeWeerd, R. A. et al. Prospectively defined patterns of APOBEC3A mutagenesis are prevalent in human cancers. Cell Rep 38, 110555 (2022). 10.1016/j.celrep.2022.110555

20 Sanchez, A. et al. Mesoscale DNA features impact APOBEC3A and APOBEC3B deaminase activity and shape tumor mutational landscapes. Nat Commun 15, 2370 (2024). 10.1038/s41467-024-45909-5

21 Harris, R. S. & Dudley, J. P. APOBECs and virus restriction. Virology 479-480C, 131–145 (2015). 10.1016/j.virol.2015.03.012

22 Burns, M. B. et al. APOBEC3B is an enzymatic source of mutation in breast cancer. Nature 494, 366–370 (2013). 10.1038/nature11881

23 Burns, M. B., Temiz, N. A. & Harris, R. S. Evidence for APOBEC3B mutagenesis in multiple human cancers. Nat Genet 45, 977–983 (2013). 10.1038/ng.2701

24 Roberts, S. A. et al. An APOBEC cytidine deaminase mutagenesis pattern is widespread in human cancers. Nat Genet 45, 970–976 (2013). 10.1038/ng.2702

25 Nik-Zainal, S. et al. The life history of 21 breast cancers. Cell 149, 994–1007 (2012). S0092-8674(12)00527-2

26 Kouno, T. et al. Crystal structure of APOBEC3A bound to single-stranded DNA reveals structural basis for cytidine deamination and specificity. Nat Commun 8, 15024 (2017). 10.1038/ncomms15024

27 Shi, K. et al. Structural basis for targeted DNA cytosine deamination and mutagenesis by APOBEC3A and APOBEC3B. Nat Struct Mol Biol 24, 131–139 (2017). 10.1038/nsmb.3344

28 Harjes, S. et al. Structure-guided inhibition of the cancer DNA-mutating enzyme APOBEC3A. Nat Commun 14, 6382 (2023). 10.1038/s41467-023-42174-w

29 Durfee, C. et al. Human APOBEC3B promotes tumor development *in vivo* including signature mutations and metastases. Cell Rep Med 4, 101211 (2023). 10.1016/j.xcrm.2023.101211

30 McCann, J. L. et al. APOBEC3B regulates R-loops and promotes transcription-associated mutagenesis in cancer. Nat Genet 55, 1721–1734 (2023). 10.1038/s41588-023-01504-w

31 Haradhvala, N. J. et al. Mutational strand asymmetries in cancer genomes reveal mechanisms of DNA damage and repair. Cell 164, 538–549 (2016). 10.1016/j.cell.2015.12.050

32 Hoopes, J. I. et al. APOBEC3A and APOBEC3B preferentially deaminate the lagging strand template during DNA replication. Cell Rep 14, 1273–1282 (2016). 10.1016/j.celrep.2016.01.021

33 Seplyarskiy, V. B. et al. APOBEC-induced mutations in human cancers are strongly enriched on the lagging DNA strand during replication. Genome Res 26, 174–182 (2016). 10.1101/gr.197046.115

34 Bergstrom, E. N. et al. Mapping clustered mutations in cancer reveals APOBEC3 mutagenesis of ecDNA. Nature 602, 510–517 (2022). 10.1038/s41586-022-04398-6

35 Nguyen, D. D. et al. The interplay of mutagenesis and ecDNA shapes urothelial cancer evolution. Nature 635, 219–228 (2024). 10.1038/s41586-024-07955-3

36 Mas-Ponte, D. & Supek, F. DNA mismatch repair promotes APOBEC3-mediated diffuse hypermutation in human cancers. Nat Genet 52, 958–968 (2020). 10.1038/s41588-020-0674-6

37 Elango, R. et al. Repair of base damage within break-induced replication intermediates promotes kataegis associated with chromosome rearrangements. Nucleic Acids Res 47, 9666–9684 (2019). 10.1093/nar/gkz651

38 Tang, X. H., Knudsen, B., Bemis, D., Tickoo, S. & Gudas, L. J. Oral cavity and esophageal carcinogenesis modeled in carcinogen-treated mice. Clin Cancer Res 10, 301–313 (2004). 10.1158/1078-0432.ccr-0999-3

39 Wang, Z. et al. Syngeneic animal models of tobacco-associated oral cancer reveal the activity of in situ anti-CTLA-4. Nat Commun 10, 5546 (2019). 10.1038/s41467-019-13471-0

40 Alexandrov, L. B. et al. Mutational signatures associated with tobacco smoking in human cancer. Science 354, 618–622 (2016). 10.1126/science.aag0299

41 Brown, W. L. et al. A rabbit monoclonal antibody against the antiviral and cancer genomic DNA mutating enzyme APOBEC3B. Antibodies (Basel) 8 (2019). 10.3390/antib8030047

42 Cancer Genome Atlas, N. Comprehensive genomic characterization of head and neck squamous cell carcinomas. Nature 517, 576–582 (2015). 10.1038/nature14129

43 Caso, R. et al. The underlying tumor genomics of predominant histologic subtypes in lung adenocarcinoma. J Thorac Oncol 15, 1844–1856 (2020). 10.1016/j.jtho.2020.08.005

44 Torrens, L. et al. The complexity of tobacco smoke-induced mutagenesis in head and neck cancer. Nat Genet 57, 884–896 (2025). 10.1038/s41588-025-02134-0

45 Huang, J. C., Svoboda, D. L., Reardon, J. T. & Sancar, A. Human nucleotide excision nuclease removes thymine dimers from DNA by incising the 22nd phosphodiester bond 5’ and the 6th phosphodiester bond 3’ to the photodimer. Proc Natl Acad Sci U S A 89, 3664–3668 (1992). 10.1073/pnas.89.8.3664

46 Hu, J. et al. Nucleotide excision repair in human cells: fate of the excised oligonucleotide carrying DNA damage *in vivo*. J Biol Chem 288, 20918–20926 (2013). 10.1074/jbc.M113.482257

47 Choi, D. H. et al. Hrq1 facilitates nucleotide excision repair of DNA damage induced by 4-nitroquinoline-1-oxide and cisplatin in *Saccharomyces cerevisiae*. J Microbiol 52, 292–298 (2014). 10.1007/s12275-014-4018-z

48 Ide, F. et al. p53 haploinsufficiency profoundly accelerates the onset of tongue tumors in mice lacking the xeroderma pigmentosum group A gene. Am J Pathol 163, 1729–1733 (2003). 10.1016/S0002-9440(10)63531-6

49 Ikenaga, M. & Kakunaga, T. Excision of 4-nitroquinoline 1-oxide damage and transformed in mouse cells. Cancer Res 37, 3672–3678 (1977).

50 Ikenaga, M., Takebe, H. & Ishii, Y. Excision repair of DNA base damage in human cells treated with the chemical carcinogen 4-nitroquinoline 1-oxide. Mutat Res 43, 415–427 (1977). 10.1016/0027-5107(77)90062-8

51 Gupta, A. et al. APOBEC3 mutagenesis drives therapy resistance in breast cancer. Nat Genet 57, 1452–1462 (2025). 10.1038/s41588-025-02187-1

52 Henderson, S., Chakravarthy, A., Su, X., Boshoff, C. & Fenton, T. R. APOBEC-mediated cytosine deamination links PIK3CA helical domain mutations to human papillomavirus-driven tumor development. Cell Rep 7, 1833–1841 (2014). 10.1016/j.celrep.2014.05.012

53 Vasan, N. et al. Double PIK3CA mutations in cis increase oncogenicity and sensitivity to PI3Kalpha inhibitors. Science 366, 714–723 (2019). 10.1126/science.aaw9032

54 Li, E. C. L. et al. Pre-incision structures reveal principles of DNA nucleotide excision repair. Nature 652, 1060–1067 (2026). 10.1038/s41586-026-10122-5

55 Ferrarone, J. R. et al. Genome-wide CRISPR screens in spheroid culture reveal that the tumor suppressor LKB1 inhibits growth via the PIKFYVE lipid kinase. Proc Natl Acad Sci U S A 121, e2403685121 (2024). 10.1073/pnas.2403685121

56 Akre, M. K. et al. Mutation processes in 293-based clones overexpressing the DNA cytosine deaminase APOBEC3B. PloS One 11, e0155391 (2016). 10.1371/journal.pone.0155391

57 Serebrenik, A. A. et al. The deaminase APOBEC3B triggers the death of cells lacking uracil DNA glycosylase. Proc Natl Acad Sci U S A 116, 22158–22163 (2019). 10.1073/pnas.1904024116

58 Morganella, S. et al. The topography of mutational processes in breast cancer genomes. Nat Commun 7, 11383 (2016). 10.1038/ncomms11383

59 Blee, A. M. et al. XPA tumor variant leads to defects in NER that sensitize cells to cisplatin. NAR Cancer 6, zcae013 (2024). 10.1093/narcan/zcae013

60 Bankmann, M., Prakash, L. & Prakash, S. Yeast RAD14 and human xeroderma pigmentosum group A DNA-repair genes encode homologous proteins. Nature 355, 555–558 (1992). 10.1038/355555a0

61. Sancar, A. DNA excision repair. Annu Rev Biochem 65, 43–81 (1996).

62 Scharer, O. D. Nucleotide excision repair in eukaryotes. Cold Spring Harbor perspectives in biology 5, a012609 (2013). 10.1101/cshperspect.a012609

63. Cancer Genome Atlas Research, N. Comprehensive genomic characterization of squamous cell lung cancers. Nature 489, 519–525 (2012). 10.1038/nature11404

64. Cancer Genome Atlas Research, N. Comprehensive molecular profiling of lung adenocarcinoma. Nature 511, 543–550 (2014). 10.1038/nature13385

65 Consortium, I. T. P.-C. A. o. W. G. Pan-cancer analysis of whole genomes. Nature 578, 82–93 (2020). 10.1038/s41586-020-1969-6

66 Islam, S. M. A. et al. Uncovering novel mutational signatures by *de novo* extraction with SigProfilerExtractor. Cell Genom 2, None (2022). 10.1016/j.xgen.2022.100179

67 Ellrott, K. et al. Scalable open science approach for mutation calling of tumor exomes using multiple genomic pipelines. Cell Syst 6, 271–281 e277 (2018). 10.1016/j.cels.2018.03.002

68 Yoshida, K. et al. Tobacco smoking and somatic mutations in human bronchial epithelium. Nature 578, 266–272 (2020). 10.1038/s41586-020-1961-1

69 Moody, S. et al. Mutational signatures in esophageal squamous cell carcinoma from eight countries with varying incidence. Nat Genet 53, 1553–1563 (2021). 10.1038/s41588-021-00928-6

70 Priestley, P. et al. Pan-cancer whole-genome analyses of metastatic solid tumours. Nature 575, 210–216 (2019). 10.1038/s41586-019-1689-y

71 Lawson, A. R. J. et al. Extensive heterogeneity in somatic mutation and selection in the human bladder. Science 370, 75–82 (2020). 10.1126/science.aba8347

72 Prokopczyk, B., Cox, J. E., Hoffmann, D. & Waggoner, S. E. Identification of tobacco-specific carcinogen in the cervical mucus of smokers and nonsmokers. J Natl Cancer Inst 89, 868–873 (1997). 10.1093/jnci/89.12.868

73 Cornish, A. J. et al. The genomic landscape of 2,023 colorectal cancers. Nature 633, 127–136 (2024). 10.1038/s41586-024-07747-9

74 Vogelstein, B. et al. Cancer genome landscapes. Science 339, 1546–1558 (2013). 10.1126/science.1235122

75 Cheng, D. T. et al. Memorial Sloan Kettering-Integrated Mutation Profiling of Actionable Cancer Targets (MSK-IMPACT): A hybridization capture-based next-generation sequencing clinical assay for solid tumor molecular oncology. J Mol Diagn 17, 251–264 (2015). 10.1016/j.jmoldx.2014.12.006

76 Efe, G. et al. Mutant p53: evolving perspectives. Genes Dev 40, 4–25 (2026). 10.1101/gad.353408.125

77 Palacio, T. et al. DNA mismatch repair mediated by Mlh1-Pms1 endonuclease-catalyzed mispair excision. Proc Natl Acad Sci U S A 122, e2528670122 (2025). 10.1073/pnas.2528670122

78 Ito, F., Fu, Y., Kao, S. A., Yang, H. & Chen, X. S. Family-wide comparative analysis of cytidine and methylcytidine deamination by eleven human APOBEC proteins. J Mol Biol 429, 1787–1799 (2017). 10.1016/j.jmb.2017.04.021

79 Chelico, L., Pham, P., Calabrese, P. & Goodman, M. F. APOBEC3G DNA deaminase acts processively 3’ --> 5’ on single-stranded DNA. Nat Struct Mol Biol 13, 392–399 (2006). 10.1038/nsmb1086

80 Roberts, S. A. & Gordenin, D. A. Clustered and genome-wide transient mutagenesis in human cancers: Hypermutation without permanent mutators or loss of fitness. Bioessays 36, 382–393 (2014). 10.1002/bies.201300140

81 Hanawalt, P. C. Transcription-coupled repair and human disease. Science 266, 1957–1958 (1994).

82 Lans, H., Hoeijmakers, J. H. J., Vermeulen, W. & Marteijn, J. A. The DNA damage response to transcription stress. Nat Rev Mol Cell Biol 20, 766–784 (2019). 10.1038/s41580-019-0169-4

83 Ning, W., Chang, P., Zheng, J. & Chen, W. Advances in immunotherapy for bladder cancer and clinical practice of next-generation sequencing. Frontiers in Immunology 16, 1684597 (2025). 10.3389/fimmu.2025.1684597

84 Zhang, C., Xu, C., Gao, X. & Yao, Q. Platinum-based drugs for cancer therapy and anti-tumor strategies. Theranostics 12, 2115–2132 (2022). 10.7150/thno.69424

85 Wang, D., Hara, R., Singh, G., Sancar, A. & Lippard, S. J. Nucleotide excision repair from site-specifically platinum-modified nucleosomes. Biochemistry 42, 6747–6753 (2003). 10.1021/bi034264k

