## Extended Data Figs 1-10 and Methods for "Carcinogen-induced DNA adducts amplify APOBEC mutagenesis through nucleotide excision repair"

**Extended Data Figures 1-10** for

### Carcinogen-induced DNA adducts amplify APOBEC mutagenesis through nucleotide excision repair

Cameron Durfee[1,*,†], Bojana Stefanovska[1,2,*], Rongjie Wu[3,4,5,*], José Córdoba-Caballero[6,*], Mahmoud A. Ibrahim[1,*], Michael A. Carpenter[1,2,#], Liping Liu[1,#], Shuvro P. Nandi[3,4,5,#], Prokopios P. Argyris[7,#], Luca Boscolo-Bielo[8,#], Eric C. L. Li[9], Nuri Alpay Temiz[10,11], Yufan Zhou[1], Erik N. Bergstrom[3,4,5], Christopher D. Mullally[1], Yaxi Wang[1], Xingyu Liu[1], Emilia Barreto Duran[1,2], Benjamin Troness[1], Joshua Proehl[1], Harshita B. Gupta[1], Allen J. York[1,2], Rachel I. Vogel[11,12], Jacob M. Wells[1], Christopher D. Steele[3,4,5], Isabeflla Stuewe[3,4,5], Daniel Muldoon[8], Jun Woo[8], Mark T. A. Donoghue[8], Laura A. Lindsey-Boltz[13], Aziz Sancar[13], Chaitanya Bandlamudi[8], Sarat Chandarlapaty[8], Wei Yang[9], Anna Malkova[1], Marcos Díaz-Gay[3,4,5,6], Ludmil B. Alexandrov[3,4,5], Reuben S. Harris[1,2]

[1] *Department of Biochemistry and Structural Biology, University of Texas Health San Antonio, San Antonio, TX, USA, 78229*
[2] *Howard Hughes Medical Institute, University of Texas Health San Antonio, San Antonio, TX, USA, 78229*
[3] *Department of Cellular and Molecular Medicine, University of California San Diego, La Jolla, CA, USA, 92093*
[4] *Department of Bioengineering, University of California San Diego, La Jolla, CA, USA, 92093*
[5] *Moores Cancer Center, University of California San Diego, La Jolla, CA, USA, 92093*
[6] *Digital Genomics Group, Cancer Genomics Program, Spanish National Cancer Research Center (CNIO), Madrid, Spain, 28029*
[7] *Department of Oral and Maxillofacial Diagnostic Sciences, Division of Oral Pathology, University of Florida College of Dentistry, Gainesville, FL, USA, 32610*
[8] *Memorial Sloan Kettering Cancer Center, New York, NY, USA, 10065*
[9] *Laboratory of Molecular Biology, National Institute of Diabetes and Digestive and Kidney Diseases, National Institutes of Health, Bethesda, MD, USA, 20892*
[10] *Institute for Health Informatics, University of Minnesota, Minneapolis, MN, USA, 55455*
[11] *Masonic Cancer Center, University of Minnesota, Minneapolis, MN, USA, 55455*
[12] *Department of Obstetrics, Gynecology, and Women's Health, University of Minnesota, Minneapolis, MN, USA, 55455*
[13] *Department of Biochemistry and Biophysics, University of North Carolina, Chapel Hill, NC, USA, 27599*
† *Present address: Stanford Cancer Institute, Stanford, CA, USA, 94305*

[*] Equal primary contributions
[#] Equal secondary contributions

**Content:** Extended Data Figs. 1-10

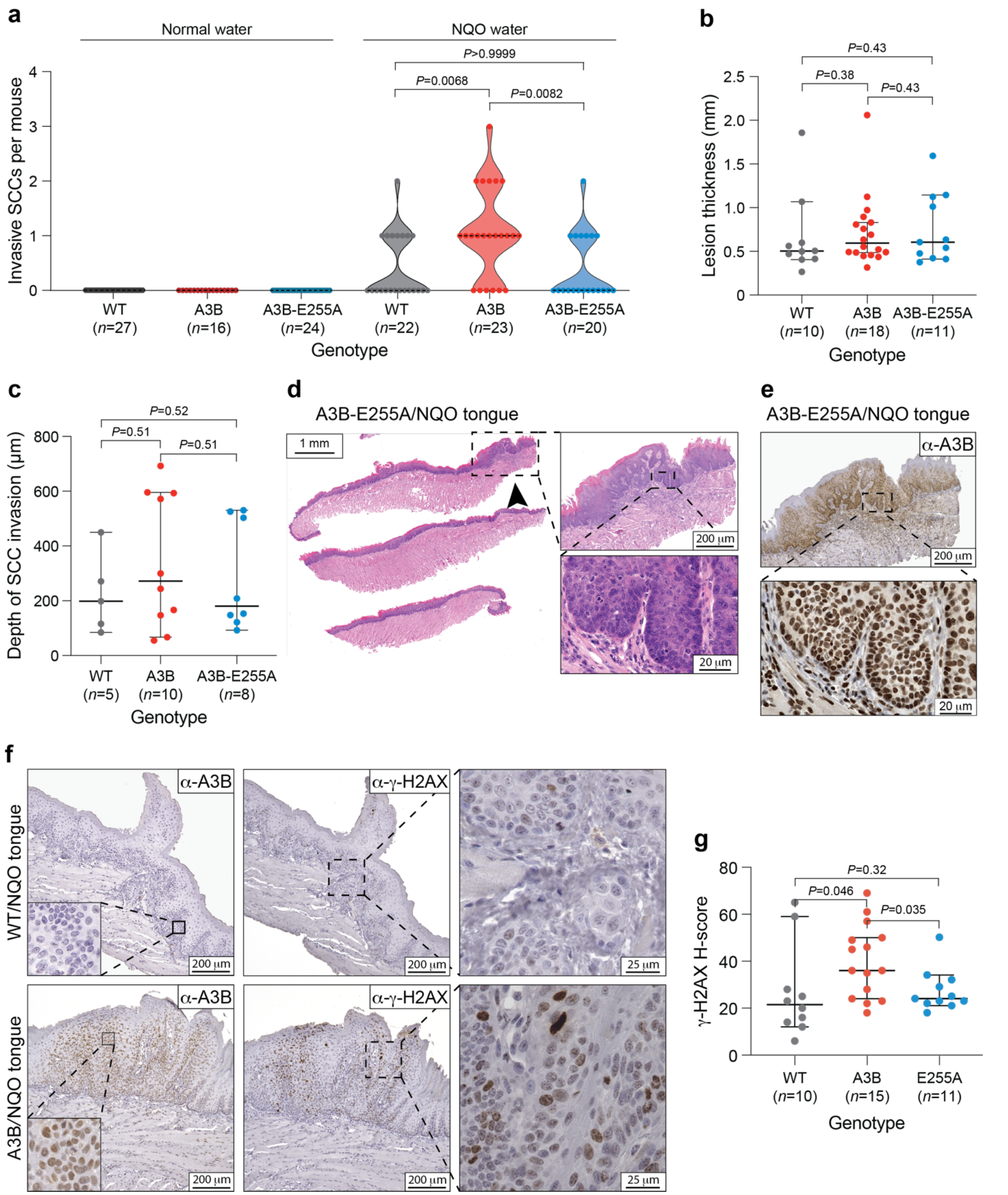
a
Normal water
NQO water
P>0.9999
P=0.0068
P=0.0082
Invasive SCCs per mouse
WT (n=27)
A3B (n=16)
A3B-E255A (n=24)
WT (n=22)
A3B (n=23)
A3B-E255A (n=20)
Genotype
b
P=0.43
P=0.38
P=0.43
Lesion thickness (mm)
WT (n=10)
A3B (n=18)
A3B-E255A (n=11)
Genotype
c
P=0.52
P=0.51
P=0.51
Depth of SCC invasion (μm)
WT (n=5)
A3B (n=10)
A3B-E255A (n=8)
Genotype
d
A3B-E255A/NQO tongue
1 mm
200 μm
20 μm
e
A3B-E255A/NQO tongue
α-A3B
200 μm
20 μm
f
WT/NQO tongue
A3B/NQO tongue
α-A3B
α-γ-H2AX
200 μm
25 μm
g
P=0.32
P=0.046
P=0.035
γ-H2AX H-score
WT (n=10)
A3B (n=15)
E255A (n=11)
Genotype

**Extended Data Fig. 1. Additional histopathology of oral lesions.**

(**a**) Quantification of oral invasive SCCs in NQO-treated animals (right) in comparison to historic controls provided with normal water (left). Each dot represents SCC quantification from an independent animal, and the dotted lines represent medians (*P*-values, pairwise two-tailed Mann-Whitney U-tests).

(**b**) Quantification of lesion thickness of the exophytic papillary high-grade epithelial dysplasias that developed in the oral cavity of animals with indicated genotypes. The horizontal lines and whiskers represent medians and 95% confidence intervals (*P*-values, pairwise two-tailed Mann-Whitney U-tests).

(**c**) Quantification of the depth of invasion of each invasive SCC that developed in the oral cavity of animals with indicated genotypes. The horizontal lines and whiskers represent medians and 95% confidence intervals (*P*-values, pairwise two-tailed Mann-Whitney U-tests).

(**d**) Representative H&E staining of a tongue from an A3B-E255A mouse, with an arrow indicating an area of high-grade epithelial dysplasia. The dysplastic area is enlarged 5x and 50x to the right, with scale bars indicated.

(**e**) A3B-E255A staining of an adjacent section of the tongue lesion shown in panel-d.

(**f**) Representative IHC images of high-grade epithelial dysplasias from WT (left) and A3B (right) mice treated with NQO. Lesions from WT animals stain negative for A3B and γ-H2AX, whereas those from A3B animals stain strongly. Nuclear A3B is most evident in the top right inset image and γ-H2AX in the bottom right panel (serial sections of the same oral SCC).

(**g**) H-score quantification of γ-H2AX staining of high-grade oral epithelial dysplasias from mice with the indicated genotypes. The horizontal lines and whiskers represent medians and 95% confidence intervals (*P*-values, pairwise two-tailed Mann-Whitney U-tests).

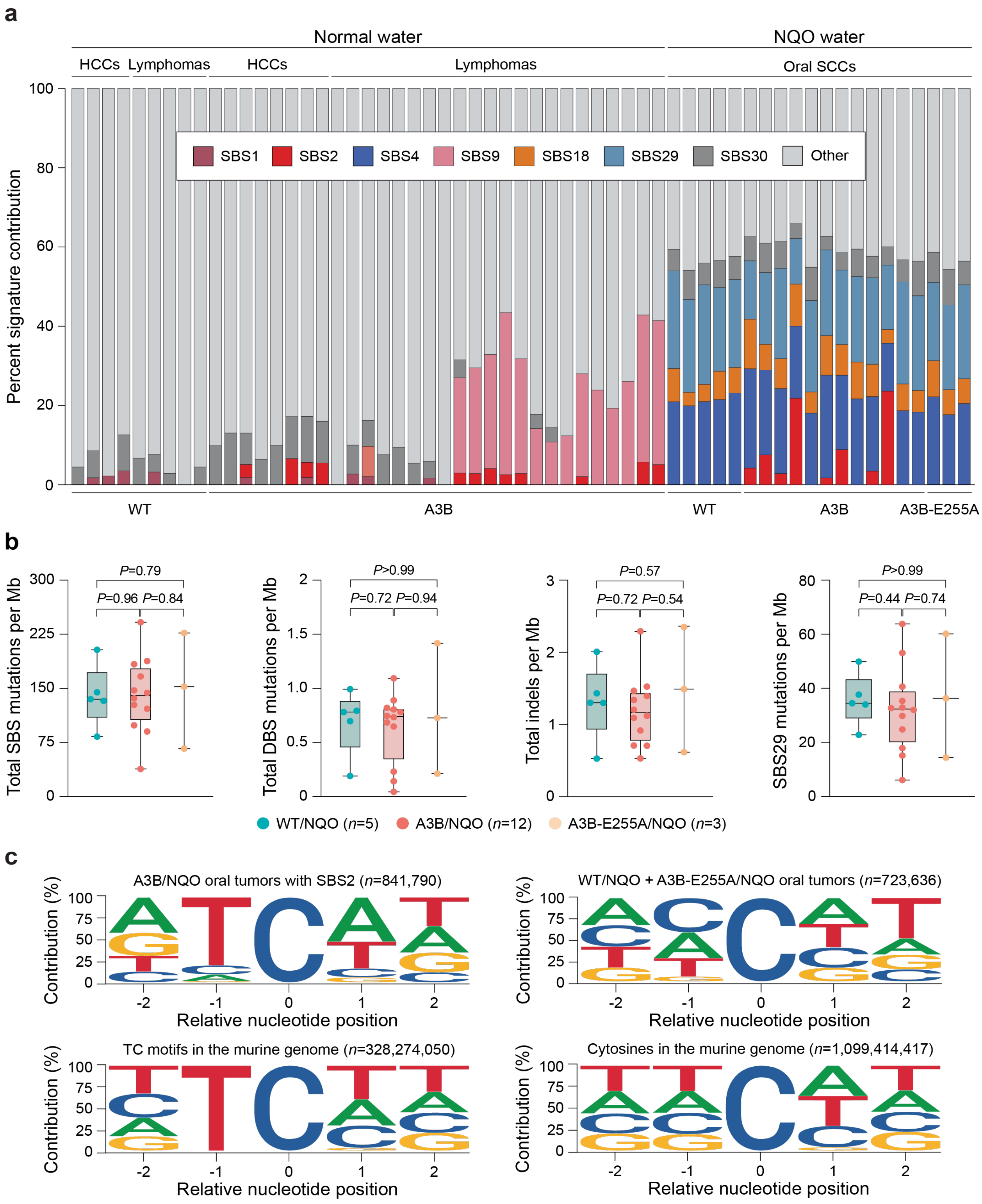
a
Normal water
NQO water
HCCs
Lymphomas
HCCs
Lymphomas
Oral SCCs
Percent signature contribution
100
80
60
40
20
0
SBS1
SBS2
SBS4
SBS9
SBS18
SBS29
SBS30
Other
WT
A3B
WT
A3B
A3B-E255A
b
Total SBS mutations per Mb
P=0.79
P=0.96 P=0.84
Total DBS mutations per Mb
P>0.99
P=0.72 P=0.94
Total indels per Mb
P=0.57
P=0.72 P=0.54
SBS29 mutations per Mb
P>0.99
P=0.44 P=0.74
WT/NQO (n=5)
A3B/NQO (n=12)
A3B-E255A/NQO (n=3)
c
A3B/NQO oral tumors with SBS2 (n=841,790)
WT/NQO + A3B-E255A/NQO oral tumors (n=723,636)
TC motifs in the murine genome (n=328,274,050)
Cytosines in the murine genome (n=1,099,414,417)
Contribution (%)
Relative nucleotide position

**Extended Data Fig. 2. Additional data on mutations in A3B/NQO tumors.**

(**a**) An alternative depiction of the data in Fig. 2a with the percentage of each SBS mutational signature illustrated. The labeling scheme and presentation order are the same as those in Fig. 2a.

(**b**) Quantification of the indicated classes of mutation in oral tumors from mice treated with NQO (SBS, single base substitutions; DBS, double base substitutions; Indels, insertion/deletions; tobacco/NQO-associated SBS29). Blue data points represent tumors from WT/NQO mice, red from A3B/NQO mice, and yellow from A3B-E255A/NQO mice. The horizontal line within each boxplot denotes the median and each box extends from the 25th to 75th percentiles (*P*-values, pairwise two-tailed Mann-Whitney U-tests).

(**c**) Pentanucleotide context of C-to-T mutations in 8 oral tumors from NQO-treated A3B animals with SBS2 (top left) in comparison to tumors from NQO-treated controls (WT and A3B-E255A, top right). For comparison, pentanucleotide context of TC motifs and C nucleobases in the murine genome are shown below. A3B/NQO tumors predominantly exhibit C-to-T mutations in TC motifs, whereas NQO control tumors show a slight bias toward CC motifs (GG on the opposite strand). *n*-values represent the total number of C-to-T mutations in each group.

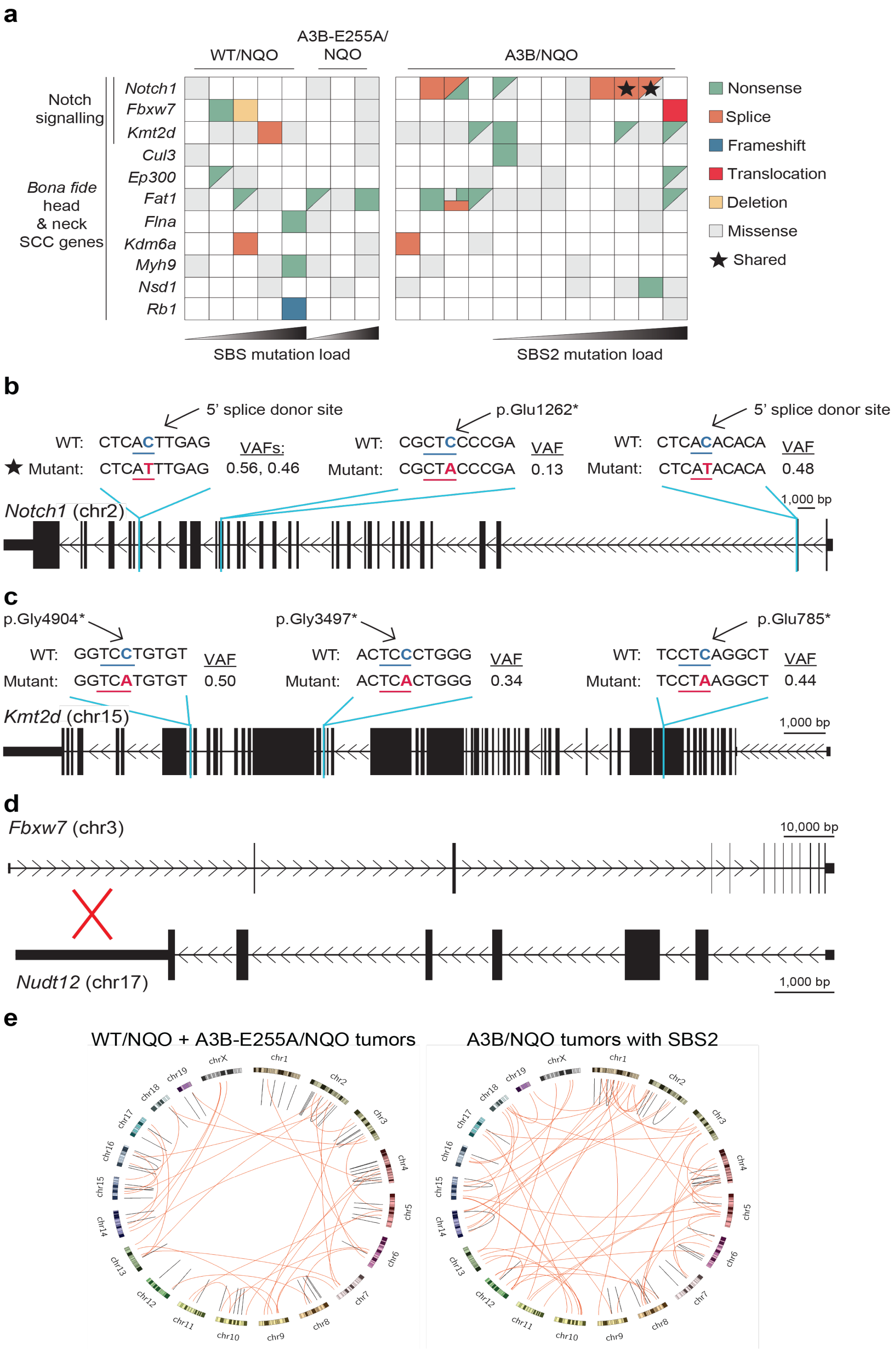
a
WT/NQO
A3B-E255A/
NQO
A3B/NQO
Notch
signalling
Bona fide
head
& neck
SCC genes
Notch1
Fbxw7
Kmt2d
Cul3
Ep300
Fat1
Flna
Kdm6a
Myh9
Nsd1
Rb1
Nonsense
Splice
Frameshift
Translocation
Deletion
Missense
Shared
SBS mutation load
SBS2 mutation load
b
5' splice donor site
WT: CTCACTTGAG
Mutant: CTCATTTGAG
VAFs:
0.56, 0.46
p.Glu1262*
WT: CGCTCCCCGA
Mutant: CGCTACCCGA
VAF
0.13
5' splice donor site
WT: CTCACACACA
Mutant: CTCATACACA
VAF
0.48
Notch1 (chr2)
1,000 bp
c
p.Gly4904*
WT: GGTCCTGTGT
Mutant: GGTCATGTGT
VAF
0.50
p.Gly3497*
WT: ACTCCCTGGG
Mutant: ACTCACTGGG
VAF
0.34
p.Glu785*
WT: TCCTCAGGCT
Mutant: TCCTAAGGCT
VAF
0.44
Kmt2d (chr15)
1,000 bp
d
Fbxw7 (chr3)
10,000 bp
Nudt12 (chr17)
1,000 bp
e
WT/NQO + A3B-E255A/NQO tumors
A3B/NQO tumors with SBS2

**Extended Data Fig. 3. Notch signaling pathway alterations in A3B/NQO tumors.**

(**a**) Oncoprint representation of mutations in tumors from the indicated NQO-treated animals. The 11 genes listed here are *bona fide* human head & neck cancer genes. Each mutation type is indicated by a different color. The black star highlights a *Notch1* splice-donor site mutation that occurred independently in two different tumors.

(**b**,**c**) Schematics of the *Notch1* and *Kmt2d* genes showing predicted high-impact mutations from panel-a (scale=1,000bp). The WT sequence is shown on top, aligned to each mutant sequence on the bottom. Variant allele frequencies (VAFs) are also shown to the right of each mutation. The black star highlights a splice-site mutation that occurred in two independent tumors (also evidenced by different VAFs).

(**d**) Schematic of the reciprocal translocation between *Fbxw7* and *Nudt12*. This translocation is predicted to disrupt *Fbxw7* expression by separating promoter region sequences from the majority of the gene body.

(**e**) Circos plots of structural variations occurring in control oral tumors (WT/NQO and A3B-E255A/NQO; *n*=8; left) and A3B/NQO oral tumors with SBS2 (*n*=8; right). Red lines represent translocations between the indicated chromosomes. Black lines represent intrachromosomal events (inversions, deletions, and duplications).

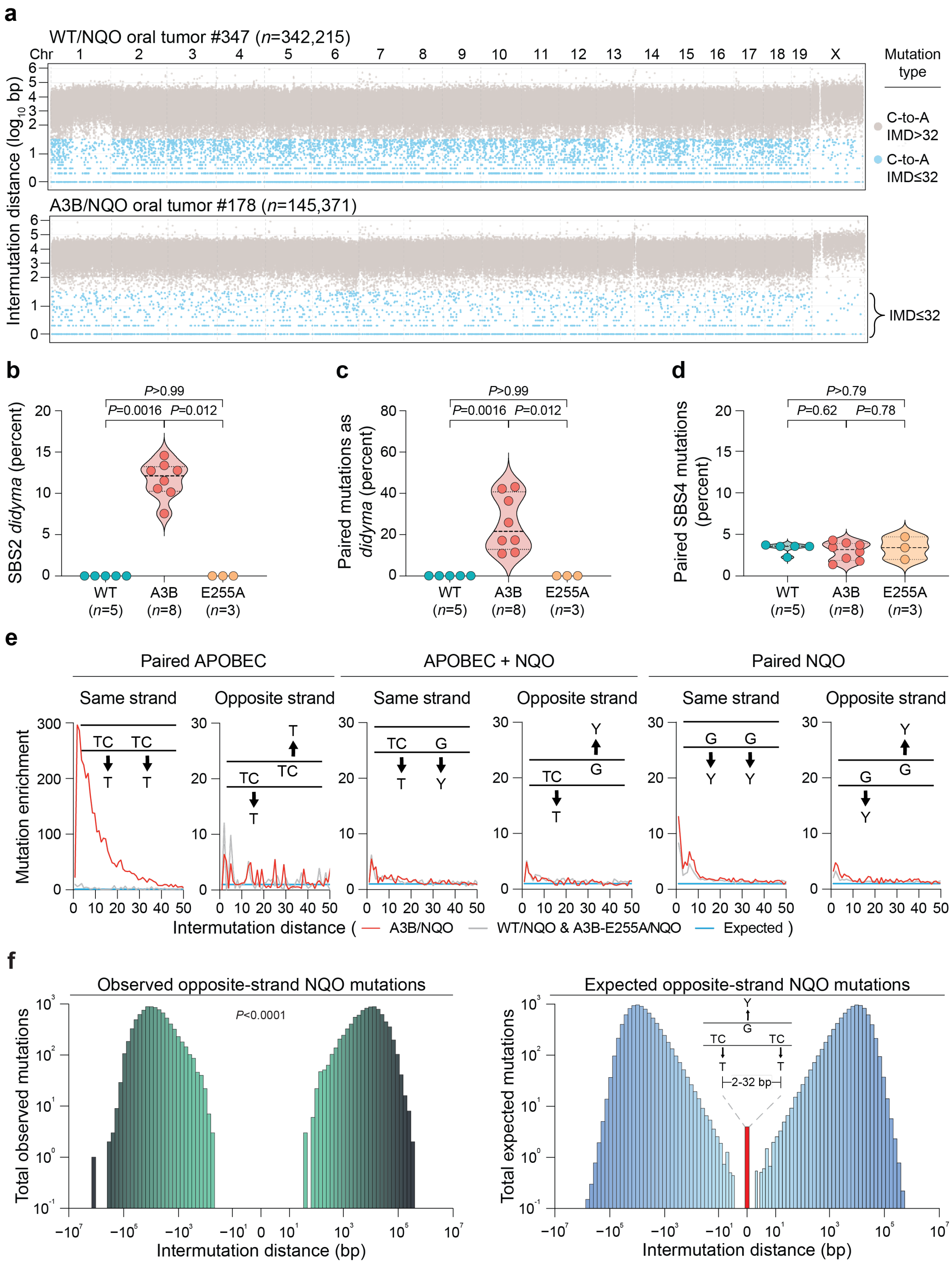

a
WT/NQO oral tumor #347 (n=342,215)
Chr 1 2 3 4 5 6 7 8 9 10 11 12 13 14 15 16 17 18 19 X
Mutation type
C-to-A IMD>32
C-to-A IMD≤32
A3B/NQO oral tumor #178 (n=145,371)
IMD≤32
Intermutation distance (log10 bp)
b
SBS2 didyma (percent)
P>0.99
P=0.0016 P=0.012
WT (n=5)
A3B (n=8)
E255A (n=3)
c
Paired mutations as didyma (percent)
P>0.99
P=0.0016 P=0.012
d
Paired SBS4 mutations (percent)
P>0.79
P=0.62 P=0.78
e
Paired APOBEC
APOBEC + NQO
Paired NQO
Same strand
Opposite strand
Mutation enrichment
Intermutation distance ( A3B/NQO WT/NQO & A3B-E255A/NQO Expected )
f
Observed opposite-strand NQO mutations
P<0.0001
Total observed mutations
Expected opposite-strand NQO mutations
Total expected mutations
2-32 bp
Intermutation distance (bp)

**Extended Data Fig. 4. Additional data on murine tumor mutations.**

(**a**) Rainfall plots depicting IMDs of C/G-to-A/T mutations from representative WT/NQO and A3B/NQO tumors. Gray and light blue symbols represent SBS mutations with IMD >32 and ≤32bp, respectively. *n*-values are total C-to-A mutations in each tumor.

(**b**,**c**,**d**) Quantification of the percentage of SBS2 *didyma* (panel-b), the contribution of *didyma* to all paired mutations with IMD≤32 in oral tumors from NQO-treated mice with the indicated genotypes (panel-c), and the percentage of paired NQO-context variations with IMD≤32 comprising all SBS4 mutations (panel-d). *n*-values for each group are shown. Dashed horizontal lines represent medians and interquartile ranges (*P*-values, pairwise two-tailed Mann-Whitney U-tests).

(**e**) Enrichment values for mutational pairs over IMD distances 1 to 50bp in A3B/NQO oral tumors with SBS2 (*n*=8; red line) or control oral tumors (WT/NQO and A3B-E255A/NQO; *n*=8; gray line) with inset schematics illustrating the queried permutation. The blue line shows the expected distribution based on simulations.

(**f**) Comparisons of observed versus expected positions of NQO-context G-to-Y mutations opposite APOBEC signature *didyma* events computed on a per-sample basis (*n*=17,229 *didyma* events in 8 A3B/NQO oral tumors; *P*-value, one-sided Fisher's exact test). The "zero" position NQO-context mutations are any events occurring between the two APOBEC signature mutations that define each *didyma* (represented by the inset schematic); all other distances are actual bp from the 5' and 3' mutational events that make up each *didyma* (*i.e.*, $\log_{10}$ increments each direction from 1 to $10^7$).

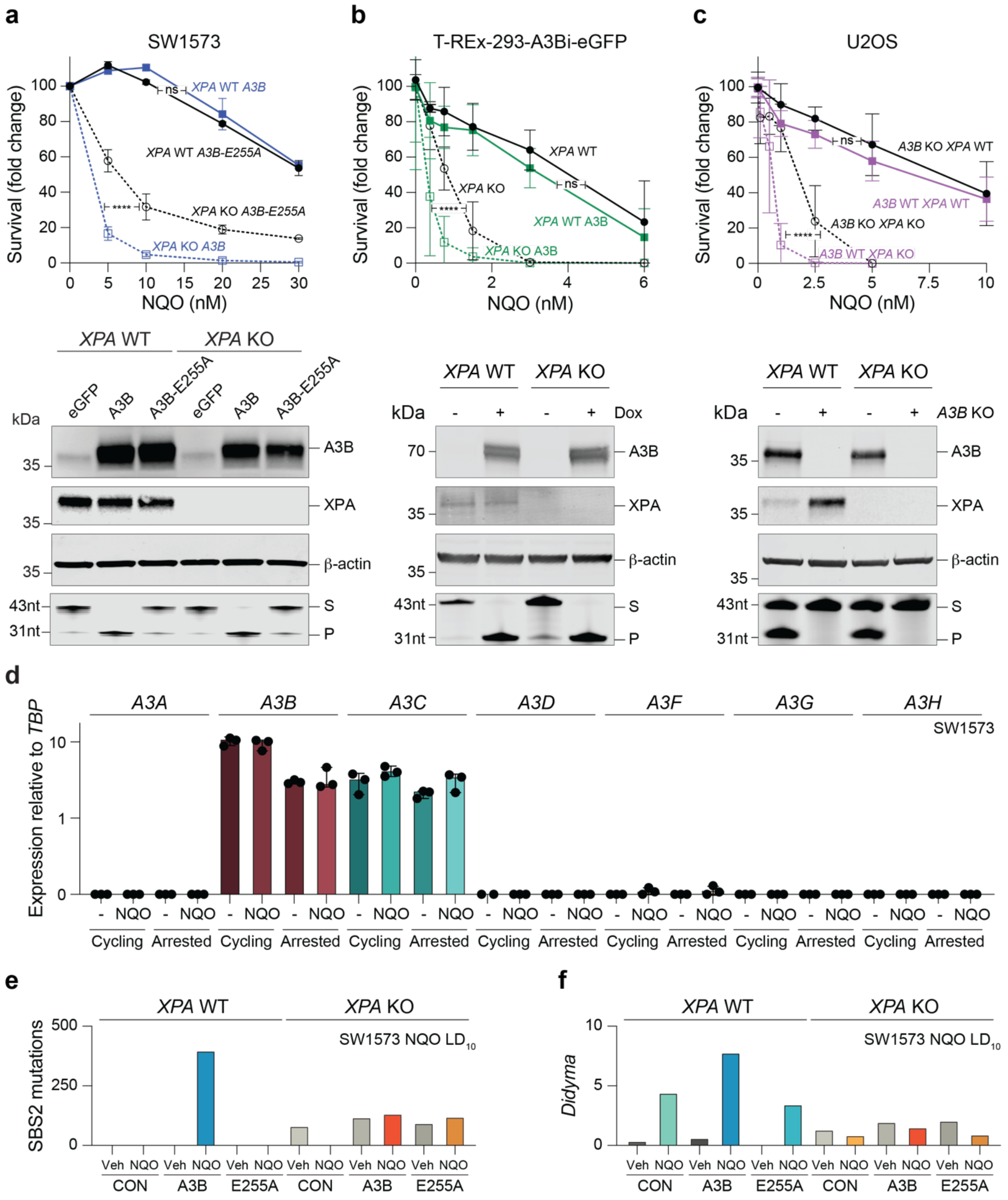
a
SW1573
Survival (fold change)
NQO (nM)
XPA WT A3B
XPA WT A3B-E255A
XPA KO A3B-E255A
XPA KO A3B
ns
****
b
T-REx-293-A3Bi-eGFP
XPA WT
XPA KO
XPA WT A3B
XPA KO A3B
c
U2OS
A3B KO XPA WT
A3B WT XPA WT
A3B KO XPA KO
A3B WT XPA KO
XPA WT
XPA KO
kDa
eGFP
A3B
A3B-E255A
Dox
A3B KO
A3B
XPA
β-actin
43nt
31nt
S
P
d
A3A
A3B
A3C
A3D
A3F
A3G
A3H
SW1573
Expression relative to TBP
NQO
Cycling
Arrested
e
SW1573 NQO LD10
SBS2 mutations
Veh
CON
E255A
f
Didyma

**Extended Data Fig. 5. Validation studies for human cell lines used here.**

(**a,b,c**) Survival curves, immunoblots, and DNA deaminase activity assays for the indicated cell lines. For NQO treatment of SW1573-derived lines, each data point is the mean +/- SD of 2 technical replicates (*P*-values, 0.39 and <0.0001 for *XPA* WT and *XPA* KO, respectively; extra sum-of-squares F-test). For T-REx-293-A3Bi-eGFP lines, A3B-eGFP was induced by treatment with 1µg/mL doxycycline for 24h, and each data point is the average of 2 independent experiments, each done in technical triplicate (*P*-values, 0.34 and <0.0001 for *XPA* WT and *XPA* KO, respectively; extra sum-of-squares F-test). For U2OS-derived lines, each data point is the average of 2 independent experiments, each done in technical triplicate (*P*-values, 0.52 and <0.0001 for *XPA* WT and *XPA* KO, respectively; extra sum-of-squares F-test). Immunoblots and DNA deaminase activity results are shown below for each set of cell lines (S, substrate; P, product).

(**d**) *APOBEC3* family member mRNA expression in cycling or growth-arrested SW1573 cells following 5µM NQO or DMSO (-) treatment. The horizontal lines and whiskers represent medians and 95% confidence intervals.

(**e**,**f**) SBS2 mutations and *didyma* per diploid genome of SW1573 cells from the indicated conditions ($LD_{10}$ of NQO).

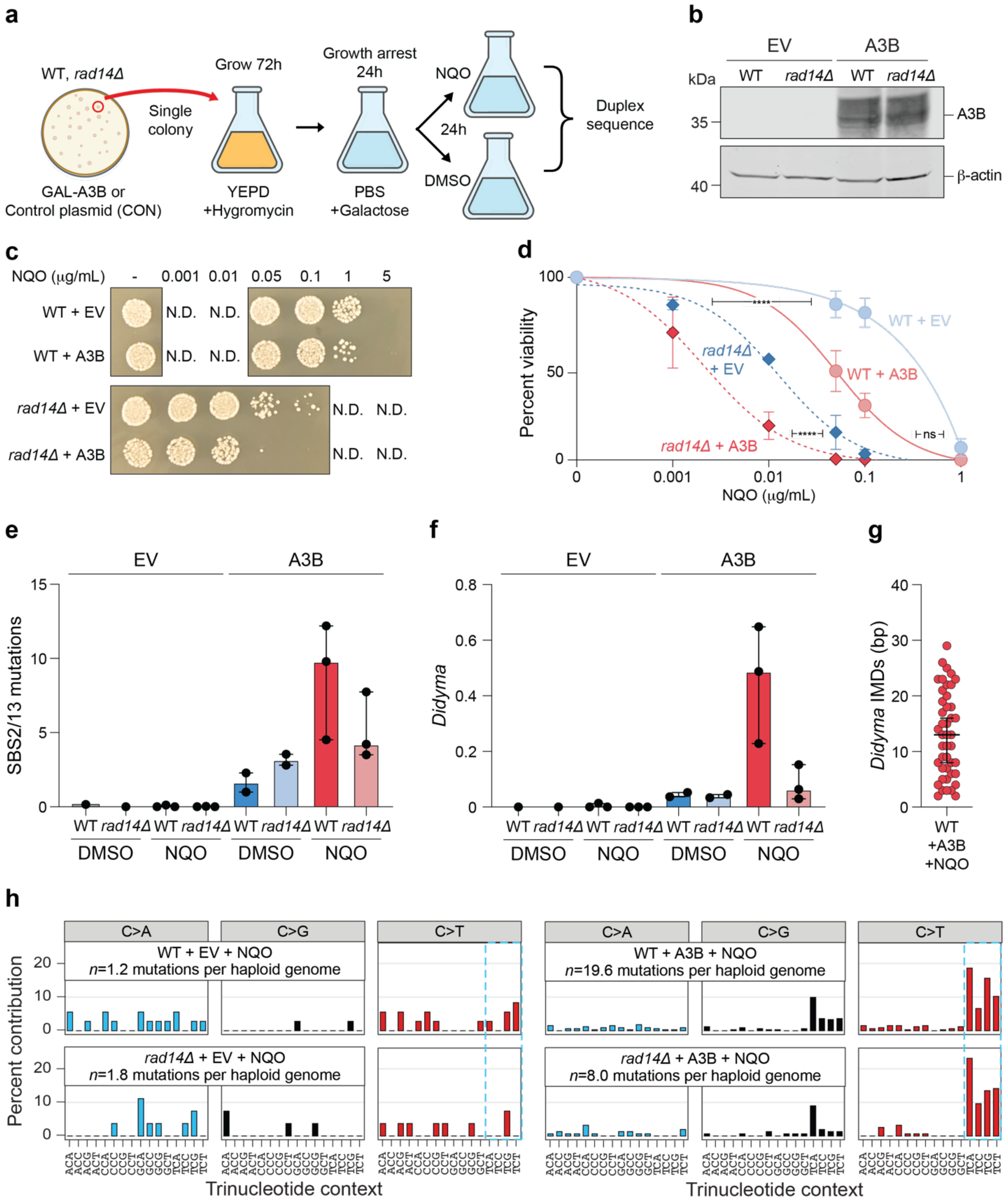
a
WT, rad14Δ
Single colony
Grow 72h
Growth arrest 24h
NQO
24h
DMSO
Duplex sequence
GAL-A3B or Control plasmid (CON)
YEPD +Hygromycin
PBS +Galactose
b
EV
A3B
kDa
WT rad14Δ WT rad14Δ
35
40
A3B
β-actin
c
NQO (μg/mL) - 0.001 0.01 0.05 0.1 1 5
WT + EV
WT + A3B
rad14Δ + EV
rad14Δ + A3B
N.D.
d
Percent viability
NQO (μg/mL)
WT + EV
WT + A3B
rad14Δ + EV
rad14Δ + A3B
****
ns
e
EV
A3B
SBS2/13 mutations
WT rad14Δ WT rad14Δ WT rad14Δ WT rad14Δ
DMSO NQO DMSO NQO
f
EV
A3B
Didyma
WT rad14Δ WT rad14Δ WT rad14Δ WT rad14Δ
DMSO NQO DMSO NQO
g
Didyma IMDs (bp)
WT +A3B +NQO
h
C>A C>G C>T
WT + EV + NQO
n=1.2 mutations per haploid genome
rad14Δ + EV + NQO
n=1.8 mutations per haploid genome
WT + A3B + NQO
n=19.6 mutations per haploid genome
rad14Δ + A3B + NQO
n=8.0 mutations per haploid genome
Percent contribution
Trinucleotide context

**Extended Data Fig. 6. Rad14 mediates A3B/NQO mutagenic synergy and *didyma* formation in yeast.**

(**a**) Schematic of workflow for growth-arrested yeast experiments.

(**b**) Immunoblots of whole-cell extracts from yeast strains with the indicated genotypes following 24h growth arrest.

(**c**) Qualitative images of WT and *rad14Δ* yeast following growth arrest, galactose addition (A3B induction), 24h NQO treatment, and outgrowth on YEPD agar plates (30°C for 48h).

(**d**) Quantification of the viability of the yeast shown in panel-c by determining colony-forming units on YEPD plates. Each data point is the mean +/- SD of $n$=3 independent experiments. Viability was determined as the percent of surviving colonies relative to those before NQO treatment ($P$-values, 0.0005 for *rad14Δ*+A3B compared to *rad14Δ*+EV, 0.0002 for *rad14Δ* +EV compared to WT+EV, and 0.08 for WT+A3B compared to WT+EV, extra sum-of-squares F test).

(**e,f**) Quantification of SBS2/13 mutations and *didyma* per haploid genome for the indicated yeast genotypes following growth arrest, A3B induction, NQO treatment, and duplex sequencing. The horizontal lines and whiskers represent medians and 95% confidence intervals.

(**g**) IMDs between the two APOBEC signature mutations comprising each *didyma* in growth-arrested WT yeast expressing A3B and treated with NQO (cumulative from 3 experiments).

(**h**) Trinucleotide distributions of all SBS mutations detected in the indicated conditions (representative experiment).

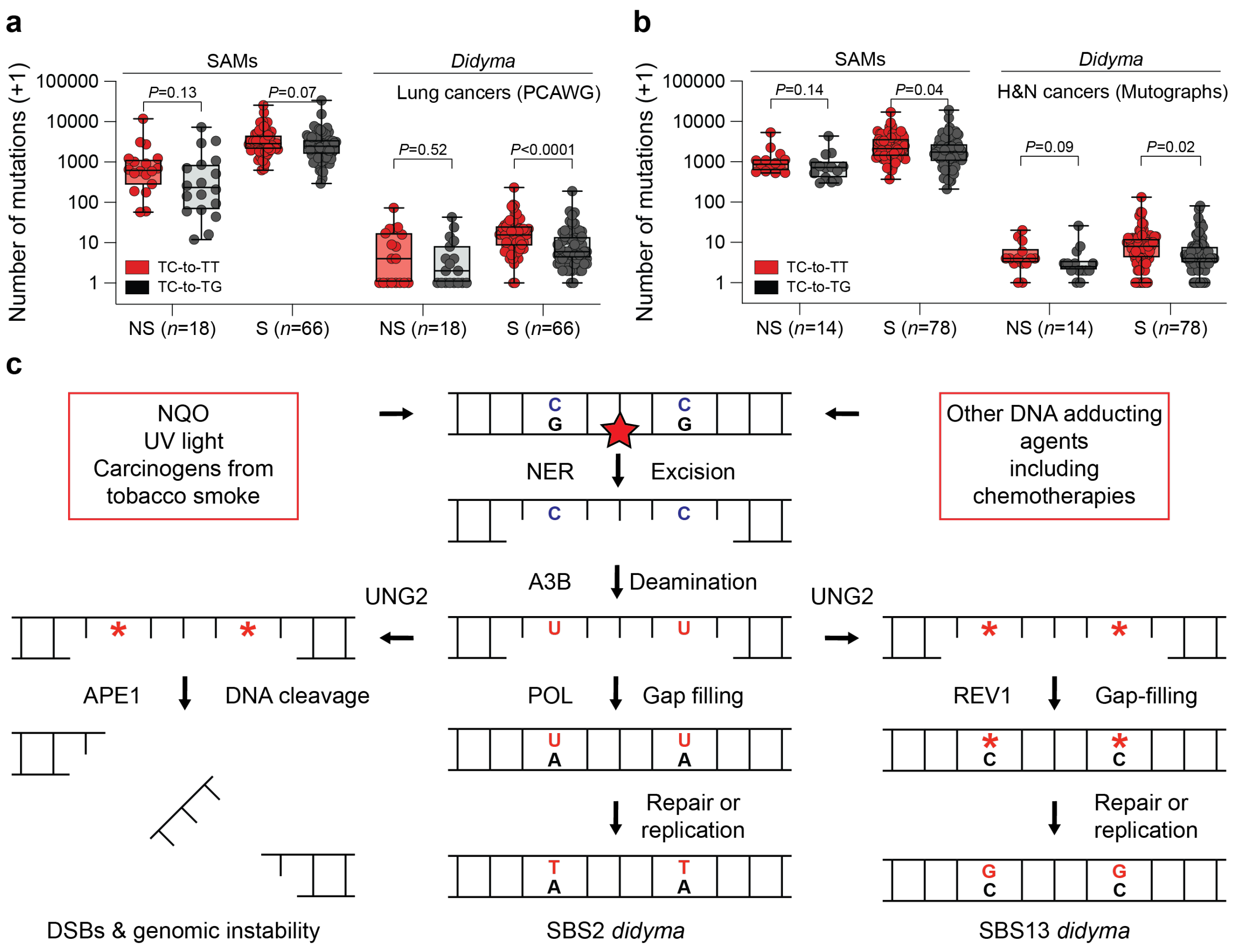
a
SAMs
Didyma
Lung cancers (PCAWG)
Number of mutations (+1)
100000
10000
1000
100
10
1
P=0.13
P=0.07
P=0.52
P<0.0001
TC-to-TT
TC-to-TG
NS (n=18)
S (n=66)
NS (n=18)
S (n=66)
b
SAMs
Didyma
H&N cancers (Mutographs)
Number of mutations (+1)
P=0.14
P=0.04
P=0.09
P=0.02
TC-to-TT
TC-to-TG
NS (n=14)
S (n=78)
NS (n=14)
S (n=78)
c
NQO
UV light
Carcinogens from
tobacco smoke
Other DNA adducting
agents
including
chemotherapies
NER
Excision
A3B
Deamination
UNG2
UNG2
APE1
DNA cleavage
POL
Gap filling
REV1
Gap-filling
Repair or
replication
Repair or
replication
DSBs & genomic instability
SBS2 didyma
SBS13 didyma

**Extended Data Fig. 7. Mutational features of *didyma* in tumor and normal tissues.**

(**a,b**) The number of APOBEC signature events, including single APOBEC mutations (SAMs) and *didyma*, that are transitions (TC-to-TT) or transversions (TC-to-TG) in lung and head & neck cancers, respectively, from non-smokers and smokers (*n*-values indicated for each group). The horizontal line within the boxplot denotes the median, and each box extends from the 25$^{th}$ to 75$^{th}$ percentiles (*P*-values, pairwise two-tailed Mann-Whitney U-tests).

(**c**) The central pathway is the same as that shown in Fig. 3d with NER creating a ssDNA gap susceptible to APOBEC-catalyzed deamination, with an additional step of repair or DNA replication immortalizing the uracil lesions as SBS2 *didyma*. The excised ssDNA strand with the bulky adduct (red star) is degraded (not illustrated). Following A3B-catalyzed C-to-U deamination, the resulting DNA uracils template the insertion of adenines during gap-filling DNA synthesis, resulting in SBS2 *didyma*. The pathway offshoot to the left depicts uracil excision by UNG2, DNA cleavage by APE1, and double-strand break formation, which would require repair by homologous recombination or potentially result in larger-scale chromosomal aberrations and elevated genomic instability. The pathway offshoot to the right depicts uracil excision by UNG2, abasic site formation (red asterisk), and error-prone gap-filling with REV1-catalyzed incorporation of cytosines opposite each abasic site. A subsequent round of repair or DNA replication will immortalize the misincorporated cytosines as SBS13 C-to-G *didyma*. Adenine insertion opposite the abasic sites can also result in SBS2 C-to-T *didyma* (not illustrated). Additional steps are not shown including competition with RPA for binding to single-stranded DNA, nucleolytic or helicase enlargement of the ssDNA tract, and alternative gap-filling DNA polymerases including POL-η (XP-V).

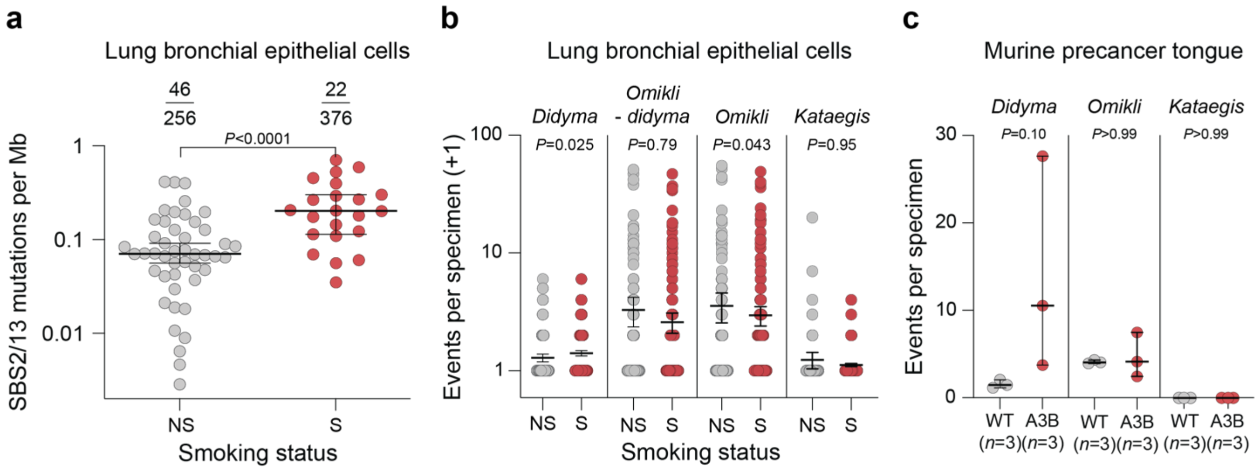


**Extended Data Fig. 8. Adducting agents increase *didyma* in normal tissues.**

(**a**) Quantification of APOBEC signature mutation loads in pathologically normal lung bronchial epithelial specimens from smokers (S) and non-smokers (NS). The horizontal lines and whiskers represent medians and 95% confidence intervals (*P*-values, two-tailed Mann-Whitney U-test). The fractions report the total number of samples with an APOBEC mutation signature over the total number analyzed.

(**b**) Quantification of APOBEC-associated mutational events in normal lung bronchial epithelial tissue from smokers (S) and non-smokers (NS; *n*=375 and 206, respectively). The horizontal lines and whiskers represent means and 95% confidence intervals (*P*-values, pairwise two-tailed Mann-Whitney U-tests). The Y-axis shows events + 1 to enable plotting on a $\log_{10}$ scale.

(**c**) Quantification of *didyma*, *omikli*, and *kataegis* from duplex sequencing of phenotypically normal tongue tissue from mice treated for 8 weeks with NQO. The horizontal lines and whiskers represent medians and 95% confidence intervals (*P*-values, pairwise two-tailed Mann-Whitney U-tests).

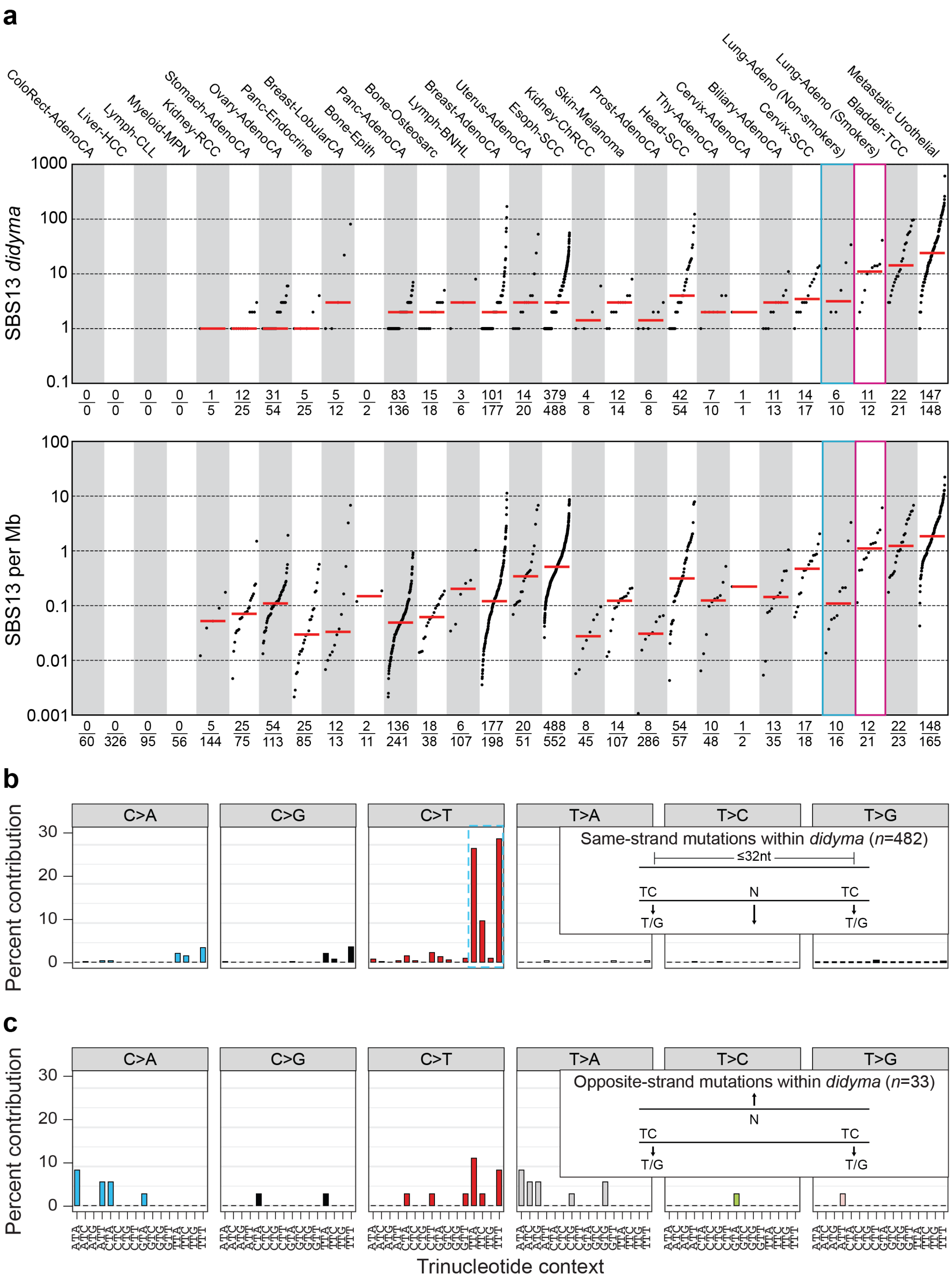
a
ColoRect-AdenoCA
Liver-HCC
Lymph-CLL
Myeloid-MPN
Kidney-RCC
Stomach-AdenoCA
Ovary-AdenoCA
Panc-Endocrine
Breast-LobularCA
Bone-Epith
Panc-AdenoCA
Bone-Osteosarc
Lymph-BNHL
Breast-AdenoCA
Uterus-AdenoCA
Esoph-SCC
Kidney-ChRCC
Skin-Melanoma
Prost-AdenoCA
Head-SCC
Thy-AdenoCA
Cervix-AdenoCA
Biliary-AdenoCA
Cervix-SCC
Lung-Adeno (Non-smokers)
Lung-Adeno (Smokers)
Bladder-TCC
Metastatic Urothelial
SBS13 didyma
SBS13 per Mb
b
Percent contribution
C>A
C>G
C>T
T>A
T>C
T>G
Same-strand mutations within didyma (n=482)
≤32nt
TC
N
TC
T/G
T/G
c
Opposite-strand mutations within didyma (n=33)
Trinucleotide context

**Extended Data Fig. 9. Additional characteristics of *didyma* across cancer types.**

(**a,b**) Quantification of SBS13 *didyma* and APOBEC signature SBS13 mutations in the indicated tumor types (order identical to Fig. 6). A SBS13 *didyma* has at least one APOBEC signature C-to-G mutation and often two (this definition only excludes twin APOBEC signature C-to-T mutations). Horizontal red lines are medians. Fractions show the number of SBS13 *didyma* over the number of tumors with SBS13, and the number of SBS13-positive tumors over the total number of tumors analyzed. Three data points below 0.001 SBS per Mb and not shown in panel-b.

(**c**,**d**) Trinucleotide distributions of SBS mutations occurring on the same strand or the opposite strand, respectively, within APOBEC signature *didyma* events in the 26 cancer types analyzed here (*n*-values in insets). This analysis indicates that some *didyma* may be *tridyma* (triplets) and that APOBEC signature mutations on the DNA strand opposing a *didyma* event are rare.

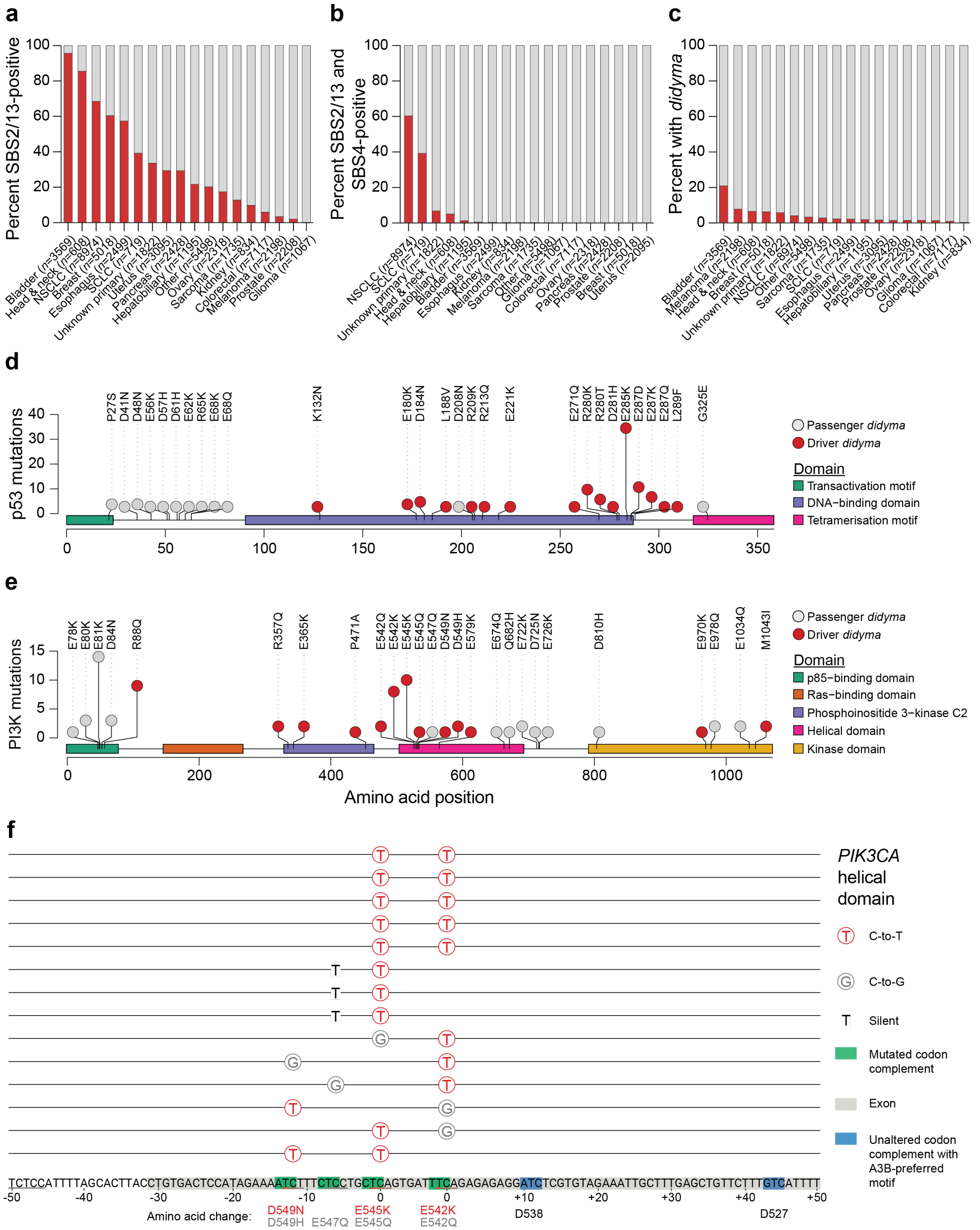
a
Percent SBS2/13-positive
Bladder (n=3569)
Head & neck (n=608)
NSCLC (n=8974)
Breast (n=5018)
Esophagus (n=2499)
SCLC (n=719)
Unknown primary (n=1822)
Uterus (n=3095)
Pancreas (n=2428)
Hepatobiliary (n=1195)
Other (n=5498)
Ovary (n=2318)
Sarcoma (n=1735)
Kidney (n=834)
Colorectal (n=7117)
Melanoma (n=2198)
Prostate (n=2208)
Glioma (n=1067)
b
Percent SBS2/13 and SBS4-positive
c
Percent with didyma
d
p53 mutations
P27S D41N D48N E56K D57H D61H E62K R65K E68K E68Q K132N E180K D184N L188V D208N R209K R213Q E221K E271Q R280K R280T D281H E285K E287D E287K E287Q L289F G325E
Passenger didyma
Driver didyma
Domain
Transactivation motif
DNA−binding domain
Tetramerisation motif
e
PI3K mutations
E78K E80K E81K D84N R88Q R357Q E365K P471A E542Q E542K E545K E545Q E547Q D549N D549H E579K E674Q Q682H E722K D725N E726K D810H E970K E978Q E1034Q M1043I
p85−binding domain
Ras−binding domain
Phosphoinositide 3−kinase C2
Helical domain
Kinase domain
Amino acid position
f
PIK3CA helical domain
C-to-T
C-to-G
Silent
Mutated codon complement
Exon
Unaltered codon complement with A3B-preferred motif
TCTCCATTTTAGCACTTACCTGTGACTCCATAGAAAATCTTTCTCCTGCTCAGTGATTTCAGAGAGAGGATCTCGTGTAGAAATTGCTTTGAGCTGTTCTTTGTCATTTT
Amino acid change:
D549N D549H E547Q E545K E545Q E542K E542Q D538 D527

**Extended Data Fig. 10. Oncogenic potential of *didyma*.**

(**a,b,c**) Quantification of the percentage of tumor specimens with SBS2/13 mutations, coincident SBS2/13 and SBS4 mutations, and *didyma*, respectively, from the MSK-IMPACT cohort. The data in each panel are ordered by descending SBS2/13 mutation prevalence (*n*-values represent the total number of specimens for each tumor type).

(**d,e**) Graphical representation of amino acid changes caused by *didyma* in P53 and PI3K, respectively. The Y-axis shows the number of times each substitution has been observed across cancers in the MSK-IMPACT cohort (precise amino acid changes are shown above each data point). The oncogenicity of each substitution (determined by OncoKB annotation) is indicated by color (red, driver; gray, passenger).

(**f**) Schematic of *didyma* in the E542 and E545 exon of the *PIK3CA* gene. These *didyma* comprise twin TC-to-TT transition mutation events (SBS2) but also TC-to-TG transversions (SBS13), with both coding and silent changes.

#### Methods

Mice

Mice were housed at the University of Minnesota Twin Cities and University of Texas Health San Antonio animal facilities in specific pathogen-free conditions at an ambient temperature of 24℃ under a standard 12h light/dark cycle. Standard breeding and husbandry for cancer studies, as well as NQO treatments, were included in protocols reviewed and approved by Institutional Animal Care and Use Committees (IACUC protocols 2201-39748A and 20220024AR, respectively).

B6.*Rosa26::CAG-LSL-A3Bi* mice and B6.*Rosa26::CAG-LSL-A3Bi-E255A* mice have been described[29] and deposited in Jackson Laboratory (Jax #038176; RRID:IMSR_JAX:038176 and #038177; RRID:IMSR_JAX:038177, respectively). These animals were crossed with B6.C-Tg(CMV-cre)1Cgn/J mice (Jax #006054) to excise the transcription STOP cassette (*i.e.*, reduce *loxP-STOP-loxP* to a single *loxP* site by Cre-mediated recombination). Subsequent crosses with WT C57BL/6 animals yielded the experimental cohorts described here including WT littermates for the control cohort. Mice were genotyped for the *Rosa26*, *Rosa26::CAG-L-A3Bi*, and *Rosa26::CAG-L-A3Bi*-E255A alleles using the following PCR conditions: 1) 95°C for 30 seconds; 2) 68°C for 30 seconds; 3) 72°C for 1min; 4) repeat steps 1 - 3 11 times; 5) 95°C for 30 seconds; 6) 68°C for 30 seconds; 7) 72°C for 1min; 8) repeat cycles 5 – 7 25 times. Primers are as follows: *Rosa26* forward: 5'-AGCACTTGCTCTCCCAAAGTC; *Rosa26* reverse: 5'-CACCTGTTCAATTCCCCTGC; *CAG-L-A3Bi* forward: 5'-CGTGCTGGTTATTGTGCTGT; and *CAG-L-A3Bi* reverse: 5'-TCCGCTCCATCGGATTTCTG.

Mice were genotyped in parallel for Cre and *Interleukin 2* (housekeeping control) using the following PCR conditions: 1) 94°C for 3min; 2) 94°C for 30 seconds; 3) 51.7°C for 1min; 4) 72°C for 1min; 5) repeat steps 2 - 4 35 times; 6) 72°C for 3min. Primers are as follows: *Cre* forward: 5'-GCGGTCTGGCAGTAAAAACTATC; *Cre* reverse: 5'- GTGAAACAGCATTGCTGTCACTT; *Interleukin-2* forward: 5'- CTAGGCCACAGAATTGAAAGATCT; and *Interleukin-2* reverse: 5'-GTAGGTGGAAATTCTAGCATCATCC.

Master mixes for all reactions consisted of final concentrations of 1x Taq buffer (Denville Scientific, C788T65), 1 mM dNTPs (Thermo Scientific, R0182), 0.3 units of Taq DNA polymerase (Thermo Scientific, EP0402), 1 µM of each primer (Integrated DNA Technologies), and 25 ng of genomic DNA.

Human cell lines

SW1573 cell lines were obtained from ATCC (CRL-2170) and maintained in Leibovitz's L-15 medium (Gibco, 11415114) supplemented with 10% FBS (Biowest, 058N24) and were maintained at 37°C using free gas exchange with atmospheric air. NHF1[86] was maintained in DMEM medium (Gibco, 11965), supplemented with 10% FBS (Biowest, 058N24). U2OS cells were obtained from ATCC (HTB-96) and maintained in RPMI medium (Gibco, 11875-093), supplemented with 10% FBS (Biowest, 058N24). Expi293F cells were obtained from Invitrogen (Gibco, A14527) and maintained in Expi293™ Expression Medium (Gibco, A1435101) at 37°C with 8% $CO_2$ shaking at 125 rpm. T-REx-293 cells (Invitrogen, R71007) were engineered to inducibly express A3Bi-eGFP[56,57]. HEK293T cells (ATCC, CRL-3216) and T-REx-293-A3Bi-eGFP cells were maintained in RPMI (Gibco, 11875093) supplemented with 10% FBS (Biowest, 058N24). NHF1, U2OS and T-REx-293 cells were maintained at 37°C with 5% $CO_2$. Cells were maintained under Mycoplasma free conditions and tested quarterly using the MycoAlert Mycoplasma Detection Kit (Lonza, LT07).

Yeast strains

The *Saccharomyces cerevisiae* yeast strain AM7658 used here was derived from the haploid strain AM3422[87], and has the following genotype: *MATa bar1Δ trp1Δ ura3Δ leu2-3,112 ade2Δ lys2::HIS4 cup1Δ yhr054cΔ cup2Δ V29616::CUP1 V34205-ADE2-lys2Δ3' hmlΔ::ADE3 ade3::GAL::HO-URA3 his4::TRP1-lys2Δ5'-NATMX his3Δ mfa1::MFA-HIS3 rpl28Q38K.* The *RAD14* gene was deleted in AM7658 at its native locus by transformation with a PCR-derived *KANMX* cassette flanked by 100bp of sequences homologous to regions surrounding the open reading frame[88], and the resulting *rad14::KANMX* strain was named AM7711. AM7658 and AM7711 were transformed either with a galactose-inducible A3B expression plasmid (*GAL1::A3B*) with a codon-optimized *A3B* sequence[89] for expression in yeast, or with the corresponding empty vector control.

Rich medium (yeast extract-peptone-dextrose; YEPD) contained 1% (w/v) yeast extract, 2% (w/v) peptone, and 2% (w/v) glucose, supplemented with 8 mL of 5% (w/v) adenine per liter and adjusted to pH 5.5. Solid media contained 2.5% (w/v) agar. All yeast cultures were grown at 30°C with agitation for liquid cultures.

Oral tumor induction and analysis

Animals of each genotype were enrolled randomly at 8 weeks of age for treatment with NQO water. NQO powder (Sigma-Aldrich, N8141) was dissolved in 100% DMSO to create a 5 mg/mL stock solution, which was subsequently diluted in water to 50 µg/mL for administration to animals. NQO water was provided continuously from week 9 to week 24, and all animals were switched to normal water for weeks 25-32. At 32 weeks of age, animals were sacrificed by $CO_2$ asphyxiation, subjected to necropsy and pathological examination, and surgically dissected for collection of tongue, oral soft tissues, esophagus, duodenal tissues, and tails. Half of each tissue was used for genomic DNA preparation, and the remainder was fixed overnight in 10% buffered formalin (10% formalin, 90% distilled water, 5 mM $Na_2HPO_4$). Tongues were trisected, embedded in paraffin blocks, stained using hematoxylin & eosin (H&E) as described below, and subsequently analyzed by a board-certified oral and maxillofacial pathologist under fully blinded conditions. Oral lesions were quantified by considering clinically or microscopically distinct exophytic papillary high-grade epithelial dysplasias and invasive SCCs in the oral cavity only, including tumors of the tongue, buccal, or labial mucosa. In addition to the oral cavity, the esophagus and duodenum of each animal were also harvested and histopathologically examined for epithelial lesions. As anticipated, lesions were confined to the oral cavity and the esophagus. Lesion thickness and depth of invasion were measured using the Keyence BZ-X800 Analyzer software. Lesion thickness was determined by measuring the distance from the apical surface of the epithelium (keratin layer) to the basal cell layer. Depth of invasion was quantified by measuring the distance from the basal cell layer of normal adjacent-to-tumor epithelium to the deepest edge of invading carcinoma nests.

Hematoxylin & eosin (H&E) staining

Formalin-fixed paraffin-embedded (FFPE) tissues were sectioned into 4 µm slices and mounted onto positively charged adhesive glass slides. Slides were subsequently baked at 60°C for 20min, washed using xylene 3 times for 5min, immersed in a series of graded alcohols (100% x 2, 95% x 1, and 80% x 1) for 2min each, and rinsed in tap water for 5min for deparaffinization and rehydration. Slides were stained with hematoxylin for 5min, rinsed in tap water for 30 seconds, subsequently submerged in an acid solution and 60 seconds in ammonia water. Slides were then washed with tap water for 10min, immersed in 80% ethanol for 1min, counterstained with eosin for 1min, dehydrated in graded alcohols (as above but inverted in increasing concentrations) followed by xylene, and coverslipped with Cytoseal Mountant (Thermo Scientific, 23-244257). High-resolution digital images were acquired using a Keyence all-in-one fluorescence microscope BZ-X800, and contrast and brightness were adjusted uniformly across images to optimize visualization.

Immunohistochemical staining

Immunohistochemistry (IHC) was done as described[29,90,91]. FFPE tissues were sectioned into 4 μm slices and mounted on positively charged adhesive slides. Tissue slices were baked at 65°C for 20min, then immersed in CitriSolv (Decon Labs, 1601) for 5min each followed by graded alcohol washes as in the preceding section and a 5min tap water rinse for deparaffinization and rehydration. 1x Reveal Decloaker (BioCare Medical, RV1000M) at pH 6.0 was used for epitope retrieval, steaming encased slides for 35min with a subsequent 30min off the steamer. Slides were then rinsed with running tap water for 5min followed by submersion in Tris-buffered saline with 0.1% Tween 20 (TBST) for 5min. Endogenous peroxidase activity was stifled with a 10min soak in 3% $H_2O_2$ diluted in TBST and successive tap water rinse for 5min. Nonspecific binding was blocked using a 15min soak in Background Sniper (BioCare Medical, BS966H). Ensuing primary antibody incubation was carried out at 4°C overnight using primary antibody diluted in 10% Background Sniper in TBST. Primary antibodies used for detection were directed against A3B (5210-87-13; RRID: AB _2721202)[41] at a 1:500 dilution and γ-H2AX Ser139 (Cell Signaling, 9718; RRID: AB_2118009) at a 1:200 dilution. Directly after overnight incubation, samples were rinsed with TBST for 5min and then incubated using Novolink Polymer (Leica Biosystems, RE7200-CE) for 30min to visualize the rabbit IgG primary antibodies. Signal was developed by application of the Novolink DAB substrate kit (Leica Biosystems, RE7140-CE) for 5min, rinsed with tap water for 5min, and counterstained with Mayer’s hematoxylin solution (Electron Microscopy Sciences, 50-317-94) for 10min. Finally, slides were washed with tap water for 10min and dehydrated in graded alcohols and CitriSolv, then cover-slipped with Permount mounting media (Thermo Scientific, SP15-100). High-resolution digital images were acquired using a Keyence all-in-one fluorescence microscope BZ-X800, and contrast and brightness were adjusted uniformly across images to optimize visualization while preserving signal fidelity. Histoscore (H-score) calculations were performed using the QuPath software[92] nuclear staining algorithm, which detects nuclear staining in cells within a manually demarcated lesional area. Lesional areas were identified based on H&E-stained slides. H-score was calculated for each lesion using the linear formula: H-score = 1x(%weak-positive cells) + 2x(%moderate-positive cells) + 3x(%strong-positive cells).

DNA extraction

Genomic DNA was extracted from fresh frozen oral tumors and matched normal tails using the DNeasy Blood and Tissue Kit (Qiagen, 69506). Tissues were homogenized using Qiashredder columns (Qiagen, 79656) and genomic DNA was prepared according to the manufacturer's instructions. Genomic DNA from cell lines was extracted using the same procedure and eluted in Buffer EB (Qiagen, 19086).

Whole-genome sequencing

100 ng genomic DNA from each murine oral tumor was used for WGS library preparation using the NEBNext Ultra II FS DNA Library Prep Kit for Illumina (New England Biolabs, E7805L). The genomic DNA was broken with an enzymatic fragmentation reaction that simultaneously repairs ends and adds dA-tails, and then each reaction was subsequently cleaned up using KAPA Pure Beads to ensure a uniform library insert size. The library was then amplified using 5 PCR cycles: 1) 98°C for 30sec; 2) 98°C for 10sec; 3) 65°C for 75sec; 4) repeat steps 2 and 3 thrice; 5) 65°C for 5min. The final DNA sequencing library was cleaned up using KAPA Pure Beads and quantified using an Invitrogen Qubit 3 Fluorometer and an Agilent TapeStation. Libraries were normalized to 10 nM and pooled at equimolar concentrations and sequencing on an Illumina NovaSeq 6000 Sequencing System to approximately 30x coverage with 150 bp paired-end sequencing. Following the sequencing run, sample demultiplexing was performed on instrument to generate FASTQ files for each sample.

Somatic mutation calling

Mouse WGS paired reads were trimmed with Trimmomatic v0.40-rc1[93] and then aligned to the mouse genome mm10 using BWA v0.7.17-r1188[94]. PCR duplicates were removed by MarkDuplicates module of GATK v4.2.6.1[95]. Reads were locally realigned around indels using RealignerTargetCreator and the IndelRealigner module of GATK3 v3.8-1-0-gf15c1c3ef. Single base substitutions and small indels were called relative to the matched normal tissues individually using Mutect2 module of GATK v4.2.6.1[96], MUSE v2.0[97], Strelka2[98], and VarScan v2.4.6[99]. Single base substitutions and small indels identified by ≥ two mutation callers were accepted as true mutations to reduce false positives. These candidate mutations were additionally filtered by requiring at least 3 reads supporting the mutation, a minimum of 10 reads at each variant site, and a variant allele frequency (VAF) over 0.05. These filtered calls were used for downstream analyses below. SnpEff was used to determine which SBS mutations and indels resulted in high- or moderate-impact mutations in genes[100].

Structural variation calling

Somatic structural variations were detected by comparing tumor to matched normal tissues and implementing four independent programs including: Manta with a minimum somatic score of 40[101]; SvABA v1.1.0[102]; DELLY[103]; and GRIDSS v2.13.2 with a quality score higher than 500[104]. Structural variations that were observed within 100 bp of each other in at least two of these algorithms were used for downstream analyses. Circos plots of structural variations were generated by Galactic Circos[105].

Mutational signature analysis

Mutational landscapes from mouse tumors were plotted using MutationalPatterns R package[106]. Known signatures from COSMICv3.4 were assigned utilizing a two pass non-negative least squares fitting where a user-defined cut-off (0.015 in this study) is applied to remove low contribution signatures after the first pass using the SigAssignR package (https://github.com/temizna/SigAssignR). Mutational signatures in human lung tumors were assigned using SigProfilerAssignment (v0.1.9) to decompose the SBS mutational signatures extracted in the original publication[66] into known signatures present in COSMICv3.4[5]. SBS mutational signature activities derived from the original publication were used for head and neck tumors[44]. For normalization of mutation burdens between different samples, we assumed 2,723 megabases (Mb) to be sequenced from mouse whole-genome sequencing, 2,800 Mb for human whole-genome sequencing, and 30 Mb sequenced for human whole-exome sequencing. For the human lung cancer datasets, a large portion lacked clinical metadata and therefore self-reported smoking status was not annotated. The existence of the tobacco smoking signature SBS4 in these samples was used to distinguish data from smokers (S) and non-smokers (NS).

Mutational context assignment

Pentanucleotide contexts were extracted using MutationalPatterns[106]. Genome-wide distributions of pentanucleotides were calculated using the mm10 genome. The mutation frequencies of each pentanucleotide context were adjusted using the genome-wide distributions of the pentanucleotides.

Rainfall plots of IMDs between mutations

For each mutation, the IMD from the preceding mutation was calculated, sans the first mutation on each chromosome. Mutations were then parsed into two groups: (i) C-to-T and C-to-G context; or (ii) C-to-A context. Mutations falling within the mm10 ENCODE blacklisted regions[107] were excluded from the analysis to avoid inconsistent mutation calling from problematic genomic areas, including telomeres and the Y chromosome. *Didyma* in mouse tissues were classified by grouping all somatic paired APOBEC-context mutations (TC-to-TT and TC-to-TG) within 32 nucleotides of each other, then ensuring 100% read concordance (*e.g.*, both mutations must be present on every single read with at least one mutation to be considered a *didyma*) from corresponding bam files. Any paired SNVs that had less than 100% read concordance were considered two separate APOBEC mutations.

IMD simulation and paired mutation calculations

Clustered mutations were extracted from detected somatic mutations of each individual sample as described[108]. Briefly, high confidence somatic mutations called from 2 of 4 different mutation callers were combined, and SigProfilerSimulator v1.1.6[109] was used to simulate a background mutation distribution on every chromosome with strand asymmetry and genic location taken into consideration. SigProfilerClusters v1.2.3[108] was used to determine sample-dependent maximum IMD, capturing 90% of mutations below IMD as unlikely to occur by chance ($q$-value $< 0.01$). Genome-wide imbalanced mutation distributions were further corrected on mutations by applying an additional regional IMD cut-off based on real and simulated mutation numbers within a 1 Mb size sliding genomic window. Maximum VAF difference with a cut-off of 0.1 was used to finalize clustered mutations, ensuring that clustered mutation events occurred in the same cells.

Paired mutations were defined based on mutational context. For APOBEC mutations, these were defined as C-to-T and C-to-G mutations in a TC context. C-to-A mutations were excluded from the quantification of APOBEC mutations and signature because they are largely eclipsed by the G/C-to-T/A mutation contribution from NQO.

For NQO, analyses of paired mutations included G-to-T and G-to-C single-base substitutions (except for G-to-C mutations in a GA context, which were excluded to avoid potential overlap with APOBEC). Mutations were classified as occurring on either the same strand or opposite strands based on the strand orientation of the reference nucleotides. Mutation enrichment was defined as the number of observed mutations divided by the number of simulated mutations. Simulated mutations were treated as the baseline expectation and therefore have an enrichment equal to 1. Here, the observed paired mutations were those that occurred in the collected tumors. Simulated paired mutations were derived from simulated mutation distribution imputed at random across each chromosome in the genome as described above. One hundred rounds of iterative simulation were performed, and the average of each type of simulated mutation pair was used for each enrichment analysis. For every IMD, the number of observed mutations was divided by the number of simulated mutations, and then values were averaged for all A3B/NQO samples or combined WT/NQO and A3B-E255A/NQO samples.

Transcription and replication topography analysis

Mutations from mouse oral tumors were assigned genomic regions based on the GRCm38.p4 genome assembly. Mutations from SW1573 cells were assigned to genomic regions based on GRCh38.p14, and mutations from PCAWG lung tumors were assigned using GRCh37.87. Transcriptomes of murine SCCs from NQO-treated and of normal tongue samples were obtained from a prior study[110]. Transcript expression was normalized using transcripts per million (TPM) to allow within-sample comparisons across genes and to correct for gene length effects, then expression of each gene was averaged and binned into quartiles or a fifth group for no expression. For mouse data, NQO and *didyma* mutations were assigned into quartiles based on gene expression from published transcriptome data[110]. Mutation burden for genic and intergenic analyses was normalized by the respective megabases within those regions, while the total transcript length of genes within each quartile was used to normalize burdens within expression quartiles. The transcribed strand bias or replication strand bias for all samples and substitution types was computed using SigProfilerTopography[111].

Computing the proximity of mutations nearest to *didyma*

For every A3B/NQO oral tumor, all read-concordant *didyma* were considered and treated as index *didyma*. Genic locations of observed index *didyma* were transposed into simulation positions (from SigProfilerClusters v1.2.3[108], as above) to assign distances to the nearest expected mutation. For observed mutation analyses, the IMD from the APOBEC mutation on the outside of the index *didyma* to the nearest SNV of each class was computed using absolute distance. Following the initial IMD assignment, distances were assigned a negative value if they are to the 5' of the index *didyma*, and positive if they are to the 3', using the 5'-TC-containing target strand as a reference. Mutations grouped into the "zero" bins include those that occur between the two APOBEC mutations comprising a single index *didyma*. Whereas VAF and read concordance were used to define observed *didyma*, proximal NQO mutation analyses here were restricted to features captured in the simulated datasets and therefore did not incorporate these parameters. Instead, NQO mutations are based only on the chromosome and position from 100 iterative simulations per tumor, and the "expected" IMDs are the average from all simulations. Statistical analyses were performed by conducting a Fisher's exact test to compare observed and the average expected NQO mutations in putative *didyma*-associated NER tracts. This includes all mutations in the zero bins representing mutations within didyma, as well as mutations in the flanking genomic area, which could represent an NER tract (*e.g.*, if the *didyma* mutations are 5 bases apart, the 13 bases upstream and downstream of the two *didyma* mutations are considered to represent the complete 32-nucleotide NER tract).

Protein Purification

A3BmycHis was expressed in Expi293F cells transfected with PEI MAX (Polysciences, 24765) at a 5:1 ratio in OptiMEM (Gibco, 31985062). 20h post-transfection, the cells were supplemented with glucose to 40 mM, valproic acid to 4.4 mM, and sodium propionate to 6.1 mM. Cells were harvested by centrifugation 3 days post-transfection and lysed in Lysis Buffer (25 mM Tris-Cl pH 8.0, 5% glycerol, 500 mM sodium chloride, 5 mM magnesium chloride, 20 mM imidazole, 0.1% IGEPAL CA-630, 50 mM L-arginine, Roche cOmplete EDTA-free protease inhibitors). Cells were then sonicated to disrupt released nucleic acids using a Branson Sonifier

450 for at least two rounds of 2min at power 5 and 50% duty cycle. RNaseA (Millipore Sigma, R5503) was added to 100 ug/mL along with 9400 units of Salt Active Ultra Nuclease (Yeasen, 20159ES60) and the lysate was incubated 1h at 37°C. The lysate was centrifuged at 16,000g for 30min at room temperature, the supernatant was collected, sodium chloride was adjusted to 1 M, and TCEP was added to 0.5 mM. The lysate was purified over Qiagen Ni-NTA Superflow resin (1018124), washed twice with Wash Buffer (25 mM Tris-Cl pH 8.0, 300 mM sodium chloride, 0.1% Triton X-100, 40 mM imidazole, 10% glycerol). Purified A3BmycHis was eluted in Elution Buffer (50 mM Tris-Cl pH 8.0, 50 mM L-arginine, 50 mM L-glutamic acid, 300 mM sodium chloride, 0.1% Triton X-100, 400 mM imidazole, 20% glycerol, 0.5 mM TCEP). The resulting protein was analyzed by UV spectroscopy and by denaturing PAGE to verify purity and quantity. The protein was then dialyzed into Storage Buffer (50 mM Tris-Cl pH 8.0, 50 mM L-arginine, 50 mM L-glutamic acid, 300 mM sodium chloride, 0.1% Triton X-100, 20% glycerol, 0.5 mM TCEP), quantified, aliquoted, and stored at -80°C.

Recombinant human nucleotide excision repair (NER) proteins including XPA, XPC trimer (XPC, RAD23B and CETN2), XPF/ERCC1, XPG, RPA and Core7 of TFIIH (XPB, XPD-Flag, p8, p62, p34, p52 and His-p44) were expressed and purified as described[112,113].

Deaminase assays

The appropriate concentration of A3BmycHis to result in ~5% deamination of oligonucleotide substrates (sequences in **Supplementary Table 2** and depicted in figure schematics) was determined by serially diluting A3BmycHis protein in reaction buffer (25 mM Tris-Cl pH 8.0, 50 mM sodium chloride, 10 mM magnesium chloride) and incubating for 15min at 37°C with 800 nM oligonucleotide substrates. Subsequent experiments were performed at a concentration of 25 nM A3BmycHis. Reactions were stopped by heating to 95°C for 5min followed by treatment with 1 μM EndoQ for 10min at 65°C. EndoQ cuts to the 5' side of uracil lesions in ssDNA[114]. Reactions were then mixed with 2x DNA PAGE loading buffer (1x TBE, 80% formamide, xylene cyanol, bromophenol blue, supplemented with 20 mM EDTA) along with SDS and Proteinase K (to final concentrations of 1% and 1 mg/mL, respectively) to remove protein:DNA complexes, heated to 95°C for 5min, and products were separated by 20% TBE-Urea PAGE. Gels were imaged on a LICORbio Odyssey M and quantified using Empiria Studio (version 2.3.0.154). Processivity Factor was determined as described[79]. The fraction of double deaminations was calculated as the product of the two sums, that is, all 5' deaminations (5' single deaminations plus double deaminations) multiplied by all 3' deaminations (3' single deaminations plus double deaminations). PF=dd/[(5'+dd)*(3'+dd)].

Deamination assays for whole cell lysates in HED buffer [25 mM HEPES, 5 mM EDTA, 10% glycerol, 1 mM DTT, and 1x cOmplete protease inhibitor (Roche, 11697498001)] from T-REx-293-A3Bi-eGFP, SW1573 and U2OS cells were performed by incubating samples at 37°C for 2 h, adding 4 pmol of a 3'-fluorescein–labeled oligonucleotide substrate (5'-ATTATTATTATTCTAATGGATTTATTTATTTATTTATTTATTT-fluorescein), 0.025 U uracil DNA glycosylase (UDG), 1× UDG reaction buffer (NEB, M0280), and 1.75 U RNase A. Following incubation, samples were treated with 100 mM NaOH and heated to 95°C for 10min to induce strand cleavage at abasic sites. Reaction products were resolved on 15% denaturing TBE–urea polyacrylamide gels to separate cleaved products from intact substrate. Fluorescent signals were detected using LICORbio Odyssey M.

<u>NER dual incision assays *in vitro*</u>

The A3B deamination experiment was performed in a 20 µl NER dual incision reaction as described[54]. Briefly, 5 nM m94_iCy5 substrate with a 5′-$^{32}$P-label on the non-Cy5 complementary strand was preincubated with 5 nM XPC at 37°C for 5 min. Then 10 nM Core7, 15 nM XPA, 10 nM RPA, 5 nM XPF/ERCC1, 5 nM XPG, and 5 nM wildtype A3B or catalytically inactive A3B-E255A mutant were added for an additional 5 min incubation at 37°C. Reaction was initiated by adding 3 mM ATP and 7.5 mM $MgCl_2$. After 10, 20, 30 and 60 minutes incubation, 1 µM EndoQ was added and incubated at 65°C for 15 minutes to cleave deaminated DNA. The reaction was then supplemented with 1 µl Proteinase K (Roche, 5mg/ml) and incubated at 55°C for 15 min to digest proteins. For analysis of NER dual-incision products, 2 µl of each reaction was mixed with TBE-urea loading buffer, heated at 95˚C for 5 min, resolved on a 15% polyacrylamide TBE-urea gel, and Cy5 fluorescence was scanned by Typhoon FLA 9500 Phosphor Imager. The remaining reaction mixture was analyzed on a separate 15% TBE-urea gel to detect $^{32}$P-labeled APOBEC3B deamination products. The gel was exposed to a phosphor imaging screen overnight and scanned using the Typhoon FLA 9500 Phosphor Imager.

<u>Generation of knockout cell lines</u>

The *A3B* knockout clone of U2OS was generated as described[30]. The *XPA* knockout clone of NHF1 has been reported[115]. All other knockout clones were generated by non-viral delivery of Cas9 ribonucleoprotein (RNP) complexes, in which Cas9 protein (Invitrogen, A36496) was complexed with a gRNA. For *XPA* knockout in T-REx-293-A3Bi-eGFP, SW1573, and U2OS cells, the gRNA targeted exon 2 of *XPA* (5'-GCCCCAAAGATAATTGACAC). For *A3B* knockout in SW1573 cells, a single gRNA targeting a sequence shared by *A3A* exon 4, *A3B* exon 7, and *A3G* exon 7 was used (5'- TGGCCCGCAGCCTCCCACTC)[116,117]. Cells were seeded and, the following day, transfected with RNPs using Lipofectamine CRISPRMAX Cas9 transfection reagent (Invitrogen, CMAX00003) according to the manufacturer's protocol. At 48 h after transfection, cells were pooled and seeded into 96-well plates at 0.5 cells per well. In parallel, control lines were generated using an identical protocol with RNPs containing a gRNA that targets no gene in the human genome (Invitrogen, A35526). Control populations were expanded sans single-cell cloning, except for those generated in parallel with the SW1573 *A3* edited clones.

*XPA* knockout clones were screened via immunoblot and validated by Sanger sequencing, as described below. To validate on-target editing, genomic DNA was extracted as described above and ~400 bp flanking the Cas9 cleavage site was amplified by PCR using the following conditions: 1) 95°C for 30sec; 2) 98°C for 10sec; 3) 60°C for 20sec; 4) 72°C for 30 sec, 5) repeat steps 2 - 4 35 times; 6) 72°C for 5min. Primers are as follows: Forward: 5'-GGTAACATACAGGCTTACCT and Reverse: 5'-GTACATCAGCCACAAGTTAA. Master mixes for all reactions consisted of final concentrations of 1x Phusion HF Buffer (Thermo Scientific, F530S), 1 mM dNTPs (Thermo Scientific, R0182), 0.02 U/μL of Phusion High–Fidelity DNA Polymerase (Thermo Scientific, F530S), 0.5 μM of each primer (Integrated DNA Technologies), and 25 ng of genomic DNA. PCR products were purified using Monarch Spin PCR & DNA Cleanup Kit (NEB, T1130L). Purified PCR products were cloned into the pJET1.2/blunt cloning vector (Thermo Fisher Scientific, K1232) according to the manufacturer's instructions and transformed into chemically competent *E. coli*. For each CRISPR clone, at least 10 independent bacterial colonies were picked, plasmid DNA was isolated by miniprep and subjected to Sanger sequencing (Genewiz). Sequencing chromatograms were analyzed using SnapGene software to identify indels at the target locus and determine allelic composition and detect insertions or deletions. For the T-REx-293-A3Bi-eGFP flow cytometry was also used to verify that the *XPA* knockout clone has a similar level of eGFP expression (and therefore A3B) following doxycycline (dox) induction. Briefly, T-REx-293-

A3Bi-eGFP cells were treated with dox (1µg/mL) or vehicle control for 48 h. Cells were then harvested, resuspended into a single-cell suspension, and subjected to flow cytometric analysis. A minimum of 10,000 events were recorded per sample to assess GFP fluorescence intensity.

*A3B* knockout clones were screened initially via immunoblot. *A3A*, *A3B*, and *A3G* loci were then assessed individually by PCR using the following forward and reverse primer pairs: *A3A* (5′-TAGGTGCCACCCCGAT and 5′-AGTGTTTGTGGAAACTCTTGC), *A3B* (5′-TGTCCCTTTTCAGAGTTTGAGTA and 5′-TGGAGCCAATTAATCACTTCAT) and *A3G* (5′-ATCAAGGAAGATGTCAGGAGGGGCT and 5′-GTTGAATGGAGCAAAGGGTGAG). Deletions spanning multiple genes were assessed by junction PCR using *A3A-B* (5'-TAGGTGCCACCCCGAT and 5'-TGGAGCCAATTAATCACTTCAT), *A3B-G* (5'-TGTCCCTTTTCAGAGTTTGAGTA and 5'-GTTGAATGGAGCAAAGGGTGAG), and *A3A-G* (5'-TAGGTGCCACCCCGAT and 5'-GTTGAATGGAGCAAAGGGTGAG) primer pairs. Master mixes for all reactions consisted of final concentrations of 1x Taq RED Master Mix (Genesee Scientific, 42137), 0.5 µM of each primer (Integrated DNA Technologies), and 25 ng of genomic DNA. PCR products were purified using Monarch Spin PCR & DNA Cleanup Kit (NEB, T1130L). Purified PCR products were cloned into the pJET1.2/blunt cloning vector and genotyped by Sanger sequencing as described above. The two SW1573 clones used in this study are both *A3B*-null due to heterozygous deletions (Δ*A3A-B* and Δ*A3B-G*).

Cellular excision assay

The cellular excision assay was performed as described[115] to detect NQO-induced excised oligos. An equal number of NHF1 cells were either untreated, treated with 5µM NQO, or irradiated with 20 $J/m^2$ UVC and incubated 2h at 37°C to allow for repair. Cells were lysed by the Hirt procedure and low molecular weight DNA in the supernatant was mixed with a 50-mer internal control oligo for 3'-end labeling with [$\alpha$-$^{32}$P]-adenosine 5′-triphosphate and separated on a DNA sequencing gel along with the indicated size markers.

Colony formation assays

Colony formation assays were performed using T-REx-293-A3Bi-eGFP, SW1573, and U2OS cells. For T-REx-293-A3Bi-eGFP *XPA* WT and *XPA* KO cells, 500 cells per well were plated for vehicle-treated controls and 750 cells per well for dox-treated conditions to account for

A3B-associated growth suppression. A3B expression was induced at the time of seeding by addition of dox (1 ng/mL). Cells were simultaneously treated with vehicle or serial dilutions of NQO prepared in 0.01% DMSO. For SW1573 (*XPA* WT and *XPA* KO; eGFP control, A3B or A3B-E255A overexpression) and U2OS (*XPA* WT/*A3B* WT, *XPA* WT/*A3B* KO, *A3B* WT/*XPA* KO, and A3B/XPA double KO) cells, 500 cells per well were plated and treated the following day with vehicle or serial dilutions of NQO (0.01% DMSO). All cells were cultured for 12-14 days (or 16-18 days for SW1573). Colonies were fixed with 70% ethanol for 10min and stained with crystal violet solution (0.5% crystal violet in 20% methanol; Sigma-Aldrich, C6158). Plates were air-dried prior to imaging using Licor Odyssey M imager. Colony numbers were manually counted using ImageJ v1.54r. Data points were excluded as outliers only when predefined criteria indicated that they were unlikely to represent the biological effect of interest and were more likely to result from technical error.

*Didyma* assays in human cells

*XPA* WT and *XPA* KO SW1573 cells were transduced with retroviral vectors to drive expression of eGFP, A3B or A3B-E255A[118]. Retroviral particles were generated by transiently transfecting HEK293T cells with either the muLV-MND-mEGFP-P2A-T2A-PuroR, muLV-MND-A3x3B-P2A-T2A-PuroR or muLV-MND-A3x3B-E255A-P2A-T2A-PuroR expression plasmids, together with the appropriate packaging and envelope plasmids. Plasmid DNA was combined at a 3:2:1 mass ratio of expression, packaging, and envelope plasmids, respectively, and transfection was performed using LT-1 reagent at an LT-1:total DNA ratio of 3:1. Viral supernatants were collected 72h after transfection and passed through a 0.45-µm filter. Cells were infected in the presence of polybrene (4 µg/mL) and treated 48h post-infection with 2 µg/mL puromycin (Gibco, A1113803) to select for successful transduction. Following selection, expression of A3B and A3B-E255A was confirmed by immunoblot, and functional activity was verified using a DNA deaminase activity assay (as described above).

For *didyma* analysis, SW1573 cells were seeded at 0.7 x $10^6$ cells per well in 6-well plates four days prior to NQO treatment to allow cells to reach contact inhibition and undergo arrest. Cell cycle distribution was assessed to confirm growth arrest, as described below. Growth arrested SW1573 cells were treated for 48h with cell line-specific concentrations of NQO, as described below. Following treatment, genomic DNA was extracted as described above and subjected to

UDSeq, as described below.

Growth arrest was validated using flow cytometric analysis of 5-ethynyl-2'-deoxyuridine (EdU) incorporation using the Click-iT Plus EdU Alexa Fluor 647 kit (Thermo Fisher Scientific, C10635) and Propidium Iodide (PI)-based DNA content assessment. Briefly, cells were incubated with 10 μM EdU for 1h at 37°C. Following labeling, cells were harvested and 1 x $10^6$ cells per condition were stained for viability to exclude non-viable cells that could confound PI-based DNA content analysis. Cells were incubated with Zombie Yellow Fixable Viability dye (BioLegend, 423104) in PBS for 40min at 4°C in the dark, followed by washing, fixation, and EdU detection as per the manufacturer's protocol. Total DNA content was determined by PI staining (20 μg/mL) in the presence of RNase A (200 μg/mL). Growth arrest was considered successful when cells showed enrichment of the 2N DNA content population together with minimal EdU incorporation, defined as at least a 90% reduction in EdU-positive cells relative to non-arrested cells analyzed in parallel. Data were acquired using LSRFortessa X-20 (BD Biosciences, 656385) and analyzed using FACSDiva and FlowJo 10.10.0 (BD Biosciences).

NQO treatment concentrations were determined as follows. SW1573 cells were seeded in flat-bottom 96-well plates and growth arrested as described above. After arrest, the cells were treated with media containing serial dilutions of NQO or 0.01% DMSO. 48h later, the CellTiter-Glo (Promega, G7570) luminescent cell viability assay was used according to the manufacturer's instructions to determine $LD_{10}$ and $LD_{50}$ concentrations. Briefly, the media was aspirated and replaced with 100 μL of a solution consisting of 1 part PBS and 1 part CellTiter-Glo reagent. Cells were lysed by orbital shaking for 2min, and then signal was developed by incubation at room temperature for 10min. Then 50 μL of the solution was transferred to white 96-well plates (Costar, 3922) and luminescence was measured using a Tecan Spark multimode microplate reader. In SW1573 cells, $LD_{50}$ concentrations were 37.8 μM for *XPA* WT and 19.5 μM for *XPA* KO cells; $LD_{10}$ concentrations were 7 μM for *XPA* WT and 4 μM for *XPA* KO cells.

Immunoblots

Cells were collected in 100 μL HED buffer (25 mM HEPES, 5 mM EDTA, 10% glycerol, 1 mM DTT, and 1x cOmplete protease inhibitor [Roche, 11697498001]) per 1 million cells. Cells were lysed by snap freezing followed by sonication for 20min in a water bath sonicator and subsequently cleared by centrifugation (16,000 x *g* for 15min). Protein concentration was quantified using a Bradford Assay (Sigma-Aldrich, B6916) at a 1:100 ratio of protein in assay reagent in triplicate. Absorbance at 595 nm wavelength was measured using a Tecan Spark multimode microplate reader. Samples were resuspended in SDS-PAGE loading buffer (62.5 mM Tris-Cl, pH 6.8; 20% glycerol, 7.5% SDS, 5% 2-mercaptoethanol) and then normalized to the same concentrations before immunoblotting and denatured by heating at 95°C for 10min. Protein was collected from yeast cells by harvesting approximately $1 \times 10^8$ cells and centrifuging at 1,000 x *g* for 2min. The pellet was washed once with 500 μL of 20% (w/v) trichloroacetic acid (TCA), followed by centrifugation at 1,000 x *g* for 2min, and the supernatant was discarded. Cells were resuspended in 200 μL of 20% TCA, mixed with 300 μL of acid-washed glass beads, and disrupted by vortexing at 4°C for 8 min. Lysates were collected by pipetting and clarified by centrifugation at 7,000 rpm for 5min, after which the TCA supernatant was discarded. The cells were neutralized by washing with 800 μL of 0.5 M Tris-HCl (pH 8.0) and pelleted again by centrifugation at 4,500 x g for 5min. Pellets were resuspended in 100 μL of 0.5 M Tris-HCl (pH 8.0) and mixed with 100 μL of 2× SDS sample buffer (0.1 M Tris-HCl, pH 6.8, 4% (w/v) SDS, 0.2% (w/v) bromophenol blue, 20% (v/v) glycerol, 5% (v/v) 2-mercaptoethanol). Samples were boiled at 100°C for 5min and clarified by centrifugation at 16,000 x *g* for 10min. The supernatants containing soluble proteins were transferred to new tubes for immunoblot analysis.

Proteins from human cells or yeast were separated via gel electrophoresis on a 4-20% gradient SDS-PAGE gel at 130 V for 90min. Protein was then transferred to a polyvinylidene difluoride Immobilon-FL membrane (Millipore, IPFL00005) using the Bio-Rad quick transfer apparatus, according to manufacturer's protocol. The membranes were then soaked in Casein blocking buffer (Sigma-Aldrich, B6429) for 1h to block nonspecific binding. The membranes were then incubated overnight at 4°C in the primary antibody. Primary antibodies used here are rabbit a-human A3A/B/G antibody 1:1000 (5210-87-13; RRID: AB _2721202)[41], rabbit a-human A3B antibody 1:1000 (Cell Signaling Technology, 41494; RRID: AB_2799203), rabbit a-XPA antibody 1:1000 (GeneTex, GTX103168; RRID: AB_1952594) in T-REx-293-A3Bi-eGFP cells, rabbit a-XPA antibody 1:1000 (Novus Biologicals, NB100-93321; RRID: AB_1237544) in all other cell lines, and mouse a-β-actin 1:5000 (Sigma-Aldrich, A1978; RRID: AB_476692). Following the primary antibody incubation, membranes were washed in PBST three times for 10min each then incubated for 1h with secondary antibodies, goat anti-rabbit HRP 1:5000 (Cell Signaling Technologies, 7074; RRID: AB_2099233) or goat anti-mouse IRDye 680LT 1:5000

(LICOR, 926-68020; RRID: AB_10706161). Following secondary antibody incubation, membranes were washed 3x in PBST, then imaged with Licor Odyssey M imager.

RT-qPCR

SW1573 cells were seeded at densities to either promote active proliferation or induce growth arrest by contact inhibition. Cells were treated with either 5µM NQO or DMSO for 4h, then total RNA was extracted using the RNeasy Mini kit (Qiagen, 74104) following the manufacturer's instructions. Extracted RNA was quantified by Nanodrop and normalized between samples. cDNA was synthesized using the ZymoScript RT PreMix Kit (R3012). Quantitative (q)PCR on DNA or cDNA samples was performed using a LightCycler 480 II instrument (Roche). RT-qPCR primers are listed in **Supplementary Table 2**.

Universal Duplex Sequencing (UDSeq) and data analysis

Duplex sequencing was done using a modified UDSeq workflow[119]. Raw sequencing reads were pre-processed by FastQC and then mapped to the human reference genome GRCh38 using the BWA-MEM aligner[120]. Variant calling was performed using DupCaller2 (https://github.com/AlexandrovLab/DupCaller), an in-house algorithm for error-corrected detection of *de novo* mutations[121]. For cell-line experiments, mutations were called against all other available BAM files used as matched normals in order to identify exposure-specific mutations induced under the respective experimental conditions. For mouse tissue, bulk whole-genome sequencing libraries were prepared and sequenced in parallel, following the protocol described above, to provide a reference for identifying sample-specific mutations. Reported *didyma*, *omikli*, and *kataegis* are all read-coordinated (*e.g.*, each mutation in the cluster is on the same read as all other mutations in the cluster). The number of mutations and *didyma* in each sample were normalized by dividing by the SNV effective coverage for each sample, and then multiplying by the number of sequenceable nucleotides in each genome (human 2,900,077,904 bp; mouse 2,561,435,404 bp; yeast 12,164,119 bp) and the ploidy (assuming human and mouse samples are diploid, and yeast are haploid).

Yeast viability assays following NQO treatment

WT and *rad14Δ* yeast cells were transformed, cultured, and induced for A3B expression as described above. At 24h after galactose induction, cultures were aliquoted and treated with NQO at the indicated concentrations. Cells were then incubated for an additional 24h at 30°C with agitation. The resulting cultures were diluted 100-fold to a final concentration of ~$4 \times 10^6$ cells/mL. Aliquots (3 µL) of the diluted cultures were spotted onto YEPD agar plates and incubated at 30 °C for 48h. Total viable cell counts were determined immediately prior to NQO addition and 24h post-treatment by serial dilution and plating onto YEPD plates, as described above. Colony-forming units were quantified after incubation, and cell viability determined as the fraction of surviving colonies relative to those before treatment.

*Didyma* assays in growth-arrested yeast

WT (*RAD14*) and *rad14Δ* yeast cells were initially cultured in 5 mL YEPD supplemented with 0.5g/L hygromycin with agitation for 24h to a density of approximately $5 \times 10^7$ cells/mL. The cultures were then expanded into 100 mL YEPD supplemented with 0.5g/L hygromycin and grown for an additional 48h to saturation, at which point cells were arrested at G1 (confirmed by no visible buds during microscopic examination). Cells were harvested by centrifugation (1,000 x *g*, 2min), washed once with sterile water, pelleted again (1,000 x *g*, 2min), and resuspended in 100 mL 1× PBS. A3B expression was induced by addition of galactose (Sigma-Aldrich, G5388) to a final concentration of 2% (w/v), followed by incubation at 30°C with agitation for 24 h. To quantify cell viability and mutation frequency, 1 mL aliquots of culture were collected immediately before and 24h after galactose induction, serially diluted, and plated onto YEPD to determine total viable cell counts. For protein expression analysis, 5 mL aliquots were collected before and after galactose induction for immunoblotting to confirm A3B expression. Following A3B induction, NQO was added to the cultures to a final concentration of 1 µg/mL, and cells were incubated at 30°C with agitation for an additional 24 h. After treatment, 1 mL aliquots were plated on YEPD for viability. The remaining cells were harvested for genomic DNA extraction. UDseq data were aligned and processed based on a custom parental yeast genome, AM3422, which was assembled by prior iterative in-house sequencing. Genome annotation was performed using the Companion genome annotation server[122] with default settings for *S. cerevisiae*, from which transcribed loci were also obtained.

Clustered mutation analysis

Sample-dependent IMDs were extrapolated from each sample (mouse and human) using SigProfilerSimulator v1.1.6[109] and SigProfilerClusters v1.2.3[108]. Only SBS mutations were included in these analyses, all indels were discarded. IMDs were calculated as the number of nucleotides separating consecutive mutations (*e.g.*, TCC = 1 IMD; TCTC = 2 IMD). For human samples, APOBEC-specific mutation clusters were extracted by assessing all mutations that fall within the sample-dependent IMD and consist entirely of APOBEC context mutations in a T[C-to-T/G]N context. They were further divided into three groups: *didyma* for strand-coordinated APOBEC-context paired mutations within 32 nucleotides of each other, *omikli* for two to three mutations and at least one IMD greater than 1 (regardless of strand-coordination); and *kataegis* for events with four or more strand-coordinated mutations including at least one IMD greater than 1. To account for potential mechanistic overlap, a simple subtractive approach was used to compare *didyma* and *omikli* numbers by including an "*omikli* minus *didyma*" category in which twin APOBEC mutation *didyma* ≤32bp were subtracted from the broader *omikli* category.

Gaussian mixture modeling of clustered event width distributions

The width of clustered mutation events was defined as the genomic distance between the first and last mutation within each event. We selected clustered *omikli* and *didyma* events consisting of TC-to-TT and TC-to-TG mutations in the TCN context with IMD <1 kb. Event widths were calculated for clustered events in lung cancer samples (LUAD and LUSC) from PCAWG and for larynx and hypopharynx tumor samples from Mutographs containing at least 1 *didyma* event. The event width distribution reflects the extent of individual mutational clusters, each of which is defined by a single underlying mutational process. The width of a cluster arising from a given process is expected to vary around a characteristic event size without extreme skew, resulting in an approximately symmetric distribution. Considering this, the distribution of event widths was modelled using Gaussian mixture modeling implemented in the Flexmix package[123] in R.

To determine the optimal number of components, models containing between one and eight Gaussian components were fitted using a maximum of 200 iterations and a convergence tolerance of $1\times10^{-6}$. The optimal solution was selected based on the Bayesian Information Criterion (BIC). Consistent with a prior report[36], both 3- and 4-component solutions provided good fits; however, we selected the three-component model because the improvement in log-likelihood between the

3- and 4-component solutions was relatively modest.

Finally, Gaussian mixture models with three components were fitted separately for smokers and non-smokers using a maximum of 10,000 iterations and a convergence tolerance of $1\times10^{-6}$. Gaussian components parameters used to model clustered mutation events in the lung cancers and head and neck cancers are as follows:

| Non-smokers | | | |
|---|---|---|---|
| Component (*k*) | 1 | 2 | 3 |
| $\mu_k$ (mean) | 28.83043 | 127.4616 | 439.4905 |
| $\sigma_k$ (standard deviation) | 20.65221 | 61.98422 | 224.1479 |
| $\pi_k$ (weight) | 0.3102 | 0.4248 | 0.2650 |
| 95 Percentile IMD | 62.7999 | 229.408 | 808.1809 |
| $P(X < 32)$ | 0.56098 | 0.061768 | 0.034535 |

| Smokers | | | |
|---|---|---|---|
| Component (*k*) | 1 | 2 | 3 |
| $\mu_k$ (mean) | 14.89932 | 86.67948 | 339.3102 |
| $\sigma_k$ (standard deviation) | 10.61939 | 52.05425 | 208.1131 |
| $\pi_k$ (weight) | 0.3435 | 0.4756 | 0.1809 |
| 95 Percentile IMD | 32.3419 | 172.2846 | 681.6258 |
| $P(X < 32)$ | 0.9463366 | 0.3754781 | 0.06988 |

Gaussian components parameters used to model clustered mutation events in the top 6 cancers by *didyma* contribution are as follows:

| Top six cancers by *didyma* contribution | | | |
|---|---|---|---|
| Component (*k*) | 1 | 2 | 3 |
| $\mu_k$ (mean) | 14.75033 | 75.07538 | 299.23514 |
| $\sigma_k$ (standard deviation) | 8.74282 | 39.18882 | 175.10552 |
| $\pi_k$ (weight) | 0.62416 | 0.17625 | 0.19959 |
| 95 Percentile IMD | 29.13098 | 139.53524 | 587.25808 |
| $P(X < 32)$ | 0.97575 | 0.13585 | 0.06249 |

<u>Gamma mixture model of IMDs</u>

Lung cancer samples (LUAD and LUSC) from the PCAWG cohort and larynx and hypopharynx tumor samples from Mutographs with at least 1 *didyma* event were included in this analysis. IMDs were computed between mutations belonging to the same clustered mutational event. The resulting IMD distribution was modeled using a mixture of Gamma distributions. We selected clustered *omikli* events consisting of TC-to-TT or TC-to-TG mutations with IMD <1 kb and TCW coordination, following criteria described previously[36].

Gamma mixture modeling was performed using the gammamixEM function from the mixtools package[124] in R to identify distinct Gamma components within the IMD distribution. Direct modeling of the raw IMD distribution frequently resulted in unstable estimates, with the algorithm collapsing to invalid solutions (shape parameter $\alpha < 0$). To improve numerical stability, IMDs were log10-transformed prior to modeling. The expectation–maximization algorithm was run with initial parameters $\alpha = 0.2$, a maximum of 10,000 iterations, and a convergence threshold $\varepsilon = 0.001$. Mixture models containing between 1 and 8 components were separately fitted to the IMD distributions of smokers and non-smokers. Model comparison based on log-likelihood indicated that both two- and three-component solutions yield acceptable fits. We elected to use a two-component model to restrict the comparison to what we assume to be *didyma* and *omikli*. Component densities were then calculated using the estimated parameters and mixture weights. Gamma component parameters were as follows:

| Non-smokers | | |
|---|---|---|
| Component (*k*) | 1 | 2 |
| $\alpha_k$ (shape) | 4.5277 | 25.3835 |
| $\beta_k$ (scale) | 0.2787 | 0.0847 |
| $\lambda_k$ (weight) | 0.1924 | 0.8076 |
| $P(X < \log10(32))$ | 0.75941 | 0.05179 |

| Smokers | | |
|---|---|---|
| Component (*k*) | 1 | 2 |
| $\alpha_k$ (shape) | 3.327 | 16.52 |
| $\beta_k$ (scale) | 0.299 | 0.115 |
| $\lambda_k$ (weight) | 0.2244 | 0.7756 |
| $P(X < \log10(32))$ | 0.83857 | 0.20399 |

<u>Duplex sequencing precancer mouse tongues</u>

*Rosa26* and *Rosa26::CAG-L-A3Bi* mice (above) were enrolled randomly at 8-weeks of age. Water containing 50 µg/mL NQO was administered to animals from weeks 9 to 16. At week 16, mice were sacrificed by $CO_2$ asphyxiation and surgically dissected for the collection of the tongue and lung. The left lateral side of the tongue and the superior, middle, and inferior lobes were snap-frozen for subsequent DNA extraction. Tissues were homogenized using Qiashredder columns, and gDNA was extracted using the DNeasy Blood and Tissue Kit. Genomic DNA was then

prepared for sequencing using UDSeq as described above.

MSK-IMPACT analyses

To investigate the association between *didyma* and established cancer-related genes, we leveraged a cohort of 79,898 patients and 93,282 tumors profiled using an FDA-authorized, tumor-normal targeted sequencing assay that profiles up to 505 cancer-associated genes (MSK-IMPACT). For downstream analysis, filtering criteria included a median sequencing depth of >100X, availability of raw BAM files for read-backed didyma calling, and samples exhibiting at least $n$=10 single-nucleotide variants, resulting in a final cohort encompassing 52,902 samples from 46,073 patients. Somatic mutations including synonymous and nonsynonymous substitutions, insertions and deletions were identified using a clinically validated pipeline[125]. Somatic mutations were deemed to be drivers if they were annotated as 'oncogenic' or 'likely oncogenic' in the FDA-recognized precision oncology knowledgebase, OncoKB[126]. To maximize the recall of *didyma*, substitutions originating from off-target reads were included in the analysis. To minimize the false positive rate, both silent mutations and off-target reads were filtered for those carrying at least $n$=100 total reads, with at least 5% of the reads supporting the alternate variant (VAF of 5%). The same criteria for calling *didyma* mutations were applied to the MSK-IMPACT sequencing cohort, and particularly at least $n$=2 strand-specific APOBEC context substitutions (TC-to-TT and TC-to-TG) falling inside a 32-base pair trinucleotide span, with a VAF difference between mutations below 10%, and substitutions supported by a read-backed variant calling (*i.e.*, didyma pairs detected in the same sequencing read). Mutational signatures were inferred using a deep-learning algorithm internally developed at Memorial Sloan Kettering (available at https://github.com/mskcc/DeepSig). A mutational signature was considered present according to a ternary call corresponding to a minimum precision of 0.9 in predicting the presence of that signature.

Statistical analyses

Comparisons were conducted using non-parametric statistical tests, namely two-tailed Mann-Whitney U-tests for comparisons, Spearman's correlation for association, and Poisson regression as noted for comparison across two independent experiments. Fisher's exact tests were used to test the number of observed mutation types to the expected, with Benjamini-Hochberg correction for each context tested where appropriate. Nonlinear regression analyses for colony formation and CellTiter-Glo assays were performed by using a four-parameter variable slope model to determine the response to compound concentrations. The extra sum-of-squares F-test was used to compare the curve fits. Details and statistical values are provided in each figure legend. Analyses were conducted using Prism 10.3.0 and SAS 9.4 (Cary, NC).

<u>Data availability</u>

All murine tumor genomic DNA sequences reported here are available in the Sequence Read Archive under accession number PRJNA1226241 (reviewer link: https://dataview.ncbi.nlm.nih.gov/object/PRJNA1226241?reviewer=nn21h3i6nq24ga1n8lg0b8vff0). Human lung cancer data were gathered from TCGA and PCAWG, which are publicly available, with somatic mutation data downloaded from ref.[66]. Similarly, somatic mutation data were publicly available for bronchial epithelial tissue[68] and human head & neck cancers[44], and these data sets were downloaded from the respective publications for new analyses here. Data for the pan-cancer analyses were gathered from PCAWG[65], Mutographs (esophageal)[69], or Hartwig (metastatic urothelial)[70].

<u>Code availability</u>

Details on all software and algorithms used in this study, including references as appropriate, can be found in individual Methods sections.

**Methods References**

86 Hu, J., Adar, S., Selby, C. P., Lieb, J. D. & Sancar, A. Genome-wide analysis of human global and transcription-coupled excision repair of UV damage at single-nucleotide resolution. *Genes Dev* **29**, 948-960 (2015). https://doi.org/10.1101/gad.261271.115

87 Liu, L. *et al.* Genome-wide screen reveals dependence of break induced replication on several distinct checkpoints. *Nat Commun* **17**, 494 (2025). https://doi.org/10.1038/s41467-025-67182-w

88 Wach, A. PCR-synthesis of marker cassettes with long flanking homology regions for gene disruptions in S. cerevisiae. *Yeast* **12**, 259-265 (1996). https://doi.org/10.1002/(SICI)1097-0061(19960315)12:3%3C259::AID-

YEA901%3E3.0.CO;2-C

89 Mertz, T. M. *et al.* Defining APOBEC-induced mutation signatures and modifying activities in yeast. *Methods Enzymol* **713**, 115-161 (2025). https://doi.org/10.1016/bs.mie.2024.11.041

90 Argyris, P. P. *et al.* Primary mucosal melanomas of the head and neck are characterised by overexpression of the DNA mutating enzyme APOBEC3B. *Histopathology* **82**, 608-621 (2023). https://doi.org/10.1111/his.14843

91 Argyris, P. P. *et al.* Endogenous APOBEC3B overexpression characterizes HPV-positive and HPV-negative oral epithelial dysplasias and head and neck cancers. *Modern Pathology* **34**, 280-290 (2021). https://doi.org/10.1038/s41379-020-0617-x

92 Bankhead, P. *et al.* QuPath: Open source software for digital pathology image analysis. *Sci Rep* **7**, 16878 (2017). https://doi.org/10.1038/s41598-017-17204-5

93 Bolger, A. M., Lohse, M. & Usadel, B. Trimmomatic: a flexible trimmer for Illumina sequence data. *Bioinformatics* **30**, 2114-2120 (2014). https://doi.org/10.1093/bioinformatics/btu170

94 Li, H. & Durbin, R. Fast and accurate short read alignment with Burrows-Wheeler transform. *Bioinformatics* **25**, 1754-1760 (2009). https://doi.org/10.1093/bioinformatics/btp324

95 McKenna, A. *et al.* The Genome Analysis Toolkit: a MapReduce framework for analyzing next-generation DNA sequencing data. *Genome Res* **20**, 1297-1303 (2010). https://doi.org/10.1101/gr.107524.110

96 Benjamin, D. *et al.* Calling Somatic SNVs and Indels with Mutect2. *bioRxiv* (2019). https://doi.org/https://doi.org/10.1101/861054

97 Fan, Y. *et al.* MuSE: accounting for tumor heterogeneity using a sample-specific error model improves sensitivity and specificity in mutation calling from sequencing data. *Genome Biol* **17**, 178 (2016). https://doi.org/10.1186/s13059-016-1029-6

98 Kim, S. *et al.* Strelka2: fast and accurate calling of germline and somatic variants. *Nat Methods* **15**, 591-594 (2018). https://doi.org/10.1038/s41592-018-0051-x

99 Koboldt, D. C. *et al.* VarScan 2: somatic mutation and copy number alteration discovery in cancer by exome sequencing. *Genome Res* **22**, 568-576 (2012). https://doi.org/10.1101/gr.129684.111

100 Cingolani, P. *et al.* A program for annotating and predicting the effects of single nucleotide

polymorphisms, SnpEff: SNPs in the genome of Drosophila melanogaster strain w1118; iso-2; iso-3. *Fly* **6**, 80-92 (2012). https://doi.org/10.4161/fly.19695

101 Chen, X. *et al.* Manta: rapid detection of structural variants and indels for germline and cancer sequencing applications. *Bioinformatics* **32**, 1220-1222 (2016). https://doi.org/10.1093/bioinformatics/btv710

102 Wala, J. A. *et al.* SvABA: genome-wide detection of structural variants and indels by local assembly. *Genome Res* **28**, 581-591 (2018). https://doi.org/10.1101/gr.221028.117

103 Rausch, T. *et al.* DELLY: structural variant discovery by integrated paired-end and split-read analysis. *Bioinformatics* **28**, i333-i339 (2012). https://doi.org/10.1093/bioinformatics/bts378

104 Cameron, D. L. *et al.* GRIDSS2: comprehensive characterisation of somatic structural variation using single breakend variants and structural variant phasing. *Genome Biol* **22**, 202 (2021). https://doi.org/10.1186/s13059-021-02423-x

105 Rasche, H. & Hiltemann, S. Galactic Circos: User-friendly Circos plots within the Galaxy platform. *Gigascience* **9** (2020). https://doi.org/10.1093/gigascience/giaa065

106 Manders, F. *et al.* MutationalPatterns: the one stop shop for the analysis of mutational processes. *BMC Genomics* **23**, 134 (2022). https://doi.org/10.1186/s12864-022-08357-3

107 Amemiya, H. M., Kundaje, A. & Boyle, A. P. The ENCODE blacklist: Identification of problematic regions of the genome. *Sci Rep* **9**, 9354 (2019). https://doi.org/10.1038/s41598-019-45839-z

108 Bergstrom, E. N., Kundu, M., Tbeileh, N. & Alexandrov, L. B. Examining clustered somatic mutations with SigProfilerClusters. *Bioinformatics* **38**, 3470-3473 (2022). https://doi.org/10.1093/bioinformatics/btac335

109 Bergstrom, E. N., Barnes, M., Martincorena, I. & Alexandrov, L. B. Generating realistic null hypothesis of cancer mutational landscapes using SigProfilerSimulator. *BMC Bioinformatics* **21**, 438 (2020). https://doi.org/10.1186/s12859-020-03772-3

110 Lee, Y. M. *et al.* Genomic and transcriptomic landscape of an oral squamous cell carcinoma mouse model for immunotherapy. *Cancer Immunol Res* **11**, 1553-1567 (2023). https://doi.org/10.1158/2326-6066.CIR-23-0133

111 Otlu, B. *et al.* Topography of mutational signatures in human cancer. *Cell Rep* **42**, 112930

(2023). https://doi.org/10.1016/j.celrep.2023.112930
112 Li, C. L. *et al.* Tripartite DNA lesion recognition and verification by XPC, TFIIH, and XPA in nucleotide excision repair. *Mol Cell* **59**, 1025-1034 (2015). https://doi.org/10.1016/j.molcel.2015.08.012
113 Kim, J. *et al.* Lesion recognition by XPC, TFIIH and XPA in DNA excision repair. *Nature* **617**, 170-175 (2023). https://doi.org/10.1038/s41586-023-05959-z
114 Belica, C. A. *et al.* RADD: A real-time FRET-based biochemical assay for DNA deaminase studies. *Methods Enzymol* **705**, 311-345 (2024). https://doi.org/10.1016/bs.mie.2024.08.001
115 Kose, C. *et al.* Cross-species investigation into the requirement of XPA for nucleotide excision repair. *Nucleic Acids Res* **52**, 677-689 (2024). https://doi.org/10.1093/nar/gkad1104
116 Ikeda, T. *et al.* HIV-1 restriction by endogenous APOBEC3G in the myeloid cell line THP-1. *J Gen Virol* **100**, 1140-1152 (2019). https://doi.org/10.1099/jgv.0.001276
117 Mittelhauser, A. *et al.* APOBEC3 knockout in lentiviral packaging cells improves therapeutic transgene sequence fidelity. *Mol Ther* (2026). https://doi.org/10.1016/j.ymthe.2026.09.030
118 Mullally, C. D. *et al.* Collapsing retroviruses for efficient delivery of viro-toxic cargoes. *bioRxiv* (2026). https://doi.org/10.64898/2026.06.28.735095
119 Nandi, S. P. *et al.* A universal duplex sequencing approach for accurate detection of somatic mutations. *bioRxiv* (2025). https://doi.org/10.1101/2025.09.14.676103
120 Li, H. Aligning sequence reads, clone sequences and assembly contigs with BWA-MEM. *arXiv:1303.3997v2* (2013). https://doi.org/https://doi.org/10.48550/arXiv.1303.3997
121 Cheng, Y. *et al.* Improved mutation detection in duplex sequencing data with sample-specific error profiles. *bioRxiv* (2025). https://doi.org/10.1101/2025.07.13.664565
122 Haese-Hill, W., Crouch, K. & Otto, T. D. Annotation and visualization of parasite, fungi and arthropod genomes with Companion. *Nucleic Acids Res* **52**, W39-W44 (2024). https://doi.org/10.1093/nar/gkae378
123 Grün, B. & Leisch, F. FlexMix version 2: Finite mixtures with concomitant variables and varying and constant parameters. *Journal of Statistical Software* **28**, 1-35 (2008). https://doi.org/10.18637/jss.v028.i04

124 Benaglia, T., Chauveau, D., Hunter, D. R. & Young, D. S. mixtools: An R package for analyzing mixture models. *Journal of Statistical Software* **32**, 1-29 (2009). https://doi.org/10.18637/jss.v032.i06

125 Zehir, A. *et al.* Mutational landscape of metastatic cancer revealed from prospective clinical sequencing of 10,000 patients. *Nat Med* **23**, 703-713 (2017). https://doi.org/10.1038/nm.4333

126 Chakravarty, D. *et al.* OncoKB: A precision oncology knowledge base. *JCO Precis Oncol* **2017** (2017). https://doi.org/10.1200/PO.17.00011

#### Acknowledgements

We thank S. Chun, S. Ishbib, and Z. Seeman for assistance with mouse colony maintenance, F.N. Papavasiliou for Greek language consultation, and O. Shärer and all members of the Harris, Alexandrov, and Díaz-Gay labs for thoughtful discussions. We are also grateful for Zhao Lai at the UT Health San Antonio Genome Sequencing Facility for assistance with DNA library preparation and sequencing and also to the South Texas Research Histology Laboratory for help with mouse specimen processing. This publication and the underlying study have been made possible partly based on data that Hartwig Medical Foundation has made available to the study through the Hartwig Medical Database.

Cancer studies in the Harris lab are supported by NCI P01-CA234228, NCI P50-CA247749, and a Recruitment of Established Investigators Award from the Cancer Prevention and Research Institute of Texas (CPRIT RR220053). N.A.T., H.B.G., and R.I.V. are also supported in part by NCI P01-CA234228. C.D. was supported by a CPRIT Research Training Award (RP 170345). Partial salary support for C.D.M. was provided by the South Texas Medical Scientist Training Program (National Institute of General and Medical Sciences T32-GM113896 and T32-GM145432) and the Epigenetics, DNA Repair, Genomics (EDGe) Training Program (NCI T32-CA279363). Research in the Malkova lab is supported by NIGMS R35-GM127006 and CPRIT RR240030. R.S.H. is an Investigator of the Howard Hughes Medical Institute, a CPRIT Scholar, and the Ewing Halsell President's Council Distinguished Chair at University of Texas Health San Antonio. The UT Health San Antonio Genome Sequencing Facility is supported by NCI P30-CA054174 (Mays Cancer Center at UT Health San Antonio) and NIH Shared Instrument grant S10-OD030311 (S10 grant to NovaSeq 6000 System), and CPRIT Core Facility Award

(RP220662). Research in the Alexandrov lab is supported by the US National Institutes of Health grants R01-ES032547, R01-CA269919, P01-CA281819, and U01-CA290479 as well as by a Packard Fellowship for Science and Engineering. The research in this study was also supported by UC San Diego Sanford Stem Cell Institute. The computational development reported in this manuscript have utilized the Triton Shared Computing Cluster at the San Diego Supercomputer Center of UC San Diego. R.I.V. and N.A.T. are supported in part by P30-CA77598 from the National Institutes of Health to the Masonic Cancer Center, University of Minnesota. A.S. and L.A.L.-B. are supported by NIH R35-GM118102. Research in the Díaz-Gay lab is supported by the project PID2024-158474NA-I00, funded by the Agencia Estatal de Investigación (AEI/10.13039/501100011033), Ministerio de Ciencia, Innovación y Universidades and cofunded by the European Regional Development Fund (ERDF-EU). J.C.-C. and M.D.-G. were awarded fellowships within the "Generación D" initiative, Red.es, Ministerio para la Transformación Digital y de la Función Pública, for talent attraction (C005/24-ED CV1), funded by the European Union NextGenerationEU funds, through PRTR. J.C.-C. and M.D.-G. are hosted by the Centro Nacional de Investigaciones Oncológicas (CNIO), which is supported by the Instituto de Salud Carlos III and recognized as a "Severo Ochoa" Centre of Excellence (CEX2024-001442-S) by the Ministerio de Ciencia, Innovación y Universidades (MCIN/AEI/10.13039/501100011033). The funders had no roles in study design, data collection and analysis, decision to publish, or preparation of the manuscript.

**Author contributions**

C.D., P.P.A., L.B.A., and R.S.H. conceptualized the project. C.D., J.P., A.J.Y., Y.W., and P.P.A. conducted experiments with mice. P.P.A. graded murine oral lesions. C.D., J.C-C., L.B.B., E.N.B., Y.Z., R.W., N.A.T., S.P.N., X.L., C.D.S., C.B., D.M., J.W., M.T.A.D., J.M.W., S.C., M.D-G., and L.B.A. performed and/or supervised computational analyses. M.A.C. did biochemical experiments with A3B, and A.S. and L.A.L-B. with NER excision. C.D., B.S., M.A.I., B.T., E.B.D., H.B.G., I.S., and C.D.M. performed experiments with human cells. L.L. and A.M. did yeast experiments. R.I.V., E.N.B., M.D-G., and L.B.A. provided statistical analysis support. C.D. and R.S.H. drafted the manuscript with all authors contributing to manuscript proofing and revision.

**Competing interests**

L.B.A. is a co-founder, CSO, scientific advisory member, and consultant for *io9* (now known as Acurion), has equity and receives income. E.N.B. is a consultant for *io9*, has equity, and receives income. The terms of these arrangements have been reviewed and approved by University of California, San Diego in accordance with its conflict of interest policies. L.B.A. is a compensated member of the scientific advisory board of Inocras. L.B.A.'s spouse is an employee of Hologic, Inc. L.B.A. and E.N.B. declare U.S. provisional applications with serial numbers: 63/289,601 and 63/269,033. L.B.A. also declares U.S. provisional applications with serial numbers: 63/366,392; 63/412,835 as well as international patent application PCT/US2023/010679. L.B.A. is also an inventor of US Patent 10,776,718 for source identification by non-negative matrix factorization. L.B.A. and M.D-G. further declare a European patent application with application number EP25305077.7. S.C. has received institutional grant/funding from Daiichi-Sankyo, Pathos.ai, and AstraZeneca, and consultation/Ad board/Honoraria from Daiichi-Sankyo, AstraZeneca, Lilly, Casdin Capital, Nuvalent, Revelio, Merck, HighTop, and Pathos AI. All other authors declare that they have no competing interests.
