## Supplemental Figs 1-6 for "Carcinogen-induced DNA adducts amplify APOBEC mutagenesis through nucleotide excision repair"

**Supplementary Figs. 1-6** for

### Carcinogen-induced DNA adducts amplify APOBEC mutagenesis through nucleotide excision repair

Cameron Durfee[1,*,†], Bojana Stefanovska[1,2,*], Rongjie Wu[3,4,5,*], José Córdoba-Caballero[6,*], Mahmoud A. Ibrahim[1,*], Michael A. Carpenter[1,2,#], Liping Liu[1,#], Shuvro P. Nandi[3,4,5,#], Prokopios P. Argyris[7,#], Luca Boscolo-Bielo[8,#], Eric C. L. Li[9], Nuri Alpay Temiz[10,11], Yufan Zhou[1], Erik N. Bergstrom[3,4,5], Christopher D. Mullally[1], Yaxi Wang[1], Xingyu Liu[1], Emilia Barreto Duran[1,2], Benjamin Troness[1], Joshua Proehl[1], Harshita B. Gupta[1], Allen J. York[1,2], Rachel I. Vogel[11,12], Jacob M. Wells[1], Christopher D. Steele[3,4,5], Isabella Stuewe[3,4,5], Daniel Muldoon[8], Jun Woo[8], Mark T. A. Donoghue[8], Laura A. Lindsey-Boltz[13], Aziz Sancar[13], Chaitanya Bandlamudi[8], Sarat Chandarlapaty[8], Wei Yang[9], Anna Malkova[1], Marcos Díaz-Gay[3,4,5,6], Ludmil B. Alexandrov[3,4,5], Reuben S. Harris[1,2]

[1] *Department of Biochemistry and Structural Biology, University of Texas Health San Antonio, San Antonio, TX, USA, 78229*
[2] *Howard Hughes Medical Institute, University of Texas Health San Antonio, San Antonio, TX, USA, 78229*
[3] *Department of Cellular and Molecular Medicine, University of California San Diego, La Jolla, CA, USA, 92093*
[4] *Department of Bioengineering, University of California San Diego, La Jolla, CA, USA, 92093*
[5] *Moores Cancer Center, University of California San Diego, La Jolla, CA, USA, 92093*
[6] *Digital Genomics Group, Cancer Genomics Program, Spanish National Cancer Research Center (CNIO), Madrid, Spain, 28029*
[7] *Department of Oral and Maxillofacial Diagnostic Sciences, Division of Oral Pathology, University of Florida College of Dentistry, Gainesville, FL, USA, 32610*
[8] *Memorial Sloan Kettering Cancer Center, New York, NY, USA, 10065*
[9] *Laboratory of Molecular Biology, National Institute of Diabetes and Digestive and Kidney Diseases, National Institutes of Health, Bethesda, MD, USA, 20892*
[10] *Institute for Health Informatics, University of Minnesota, Minneapolis, MN, USA, 55455*
[11] *Masonic Cancer Center, University of Minnesota, Minneapolis, MN, USA, 55455*
[12] *Department of Obstetrics, Gynecology, and Women's Health, University of Minnesota, Minneapolis, MN, USA, 55455*
[13] *Department of Biochemistry and Biophysics, University of North Carolina, Chapel Hill, NC, USA, 27599*
† *Present address: Stanford Cancer Institute, Stanford, CA, USA, 94305*

[*] Equal primary contributions
[#] Equal secondary contributions

**Content:** Supplementary Figs. 1-6

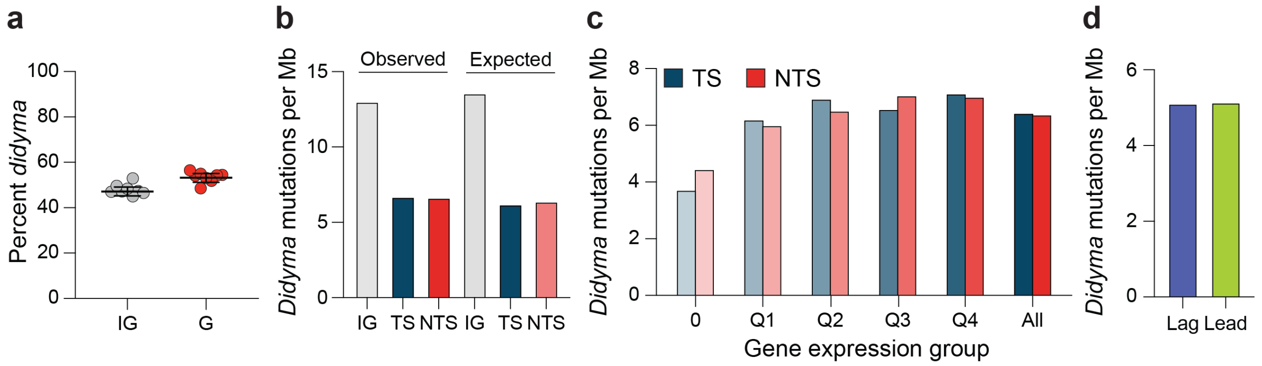


**Supplementary Fig. 1. Topography of *didyma* in murine oral SCC.**

(**a**) Percentage of APOBEC signature *didyma* (C-to-T) per megabase in intergenic (IG) versus genic (G) regions of tumors from A3B/NQO animals (*n*=8). Horizontal lines represent means, and whiskers represent 95% confidence intervals.

(**b**) Quantification of *didyma* in the indicated genomic regions in A3B/NQO tumors with SBS2 (*n*=8; IG, intergenic; TS, transcribed strand; non-transcribed strand, NTS). Observed mutations are from tumor data, and expected are from simulations (*q*-value, Fisher's exact test with Benjamini-Hochberg correction for each context tested).

(**c**) Quantification of *didyma* per Mb for the indicated gene expression groups in A3B/NQO tumors (*n*=8; 0, no expression; Q1-Q4, expression quartile; All, all groups combined).

(**d**) Quantification of *didyma* in A3B/NQO tumors (*n*=8) with respect to DNA replication leading or lagging strand (lead and lag, respectively).

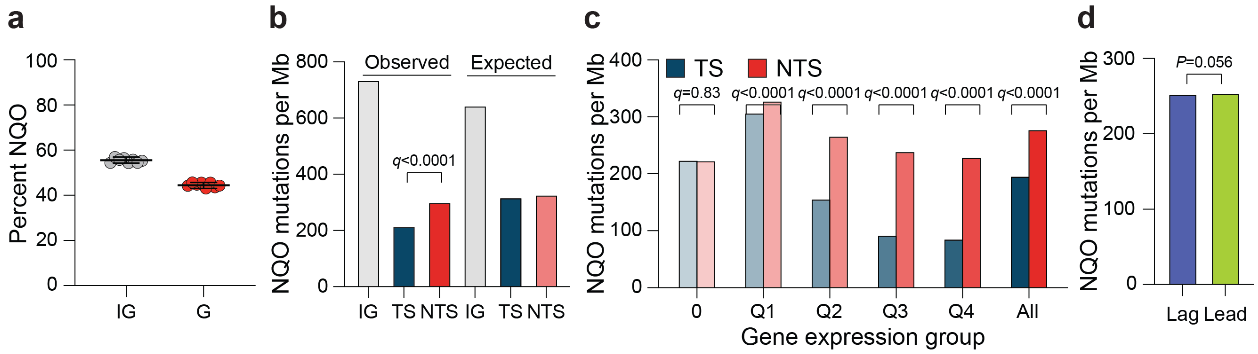


**Supplementary Fig. 2. Topography of NQO mutations in murine oral SCC.**

(**a**) Percentage of NQO mutations (G-to-Y) per megabase in intergenic (IG) versus genic (G) regions of tumors from A3B/NQO animals (*n*=8). Horizontal lines represent means, and whiskers represent 95% confidence intervals.

(**b**) Quantification of NQO mutations in the indicated regions in A3B/NQO tumors with SBS2 (*n*=8; IG, intergenic; TS, transcribed strand; non-transcribed strand, NTS). Observed mutations are from tumor data, and expected are from simulations (*q*-value, Fisher's exact test with Benjamini-Hochberg correction for each context tested).

(**c**) Quantification of NQO mutations per Mb for the indicated gene expression groups in A3B/NQO tumors (*n*=8; 0, no expression; Q1-Q4, expression quartile; All, all groups combined).

(**d**) Quantification of NQO mutations in A3B/NQO tumors (*n*=8) with respect to DNA replication leading or lagging strand (lead and lag, respectively).

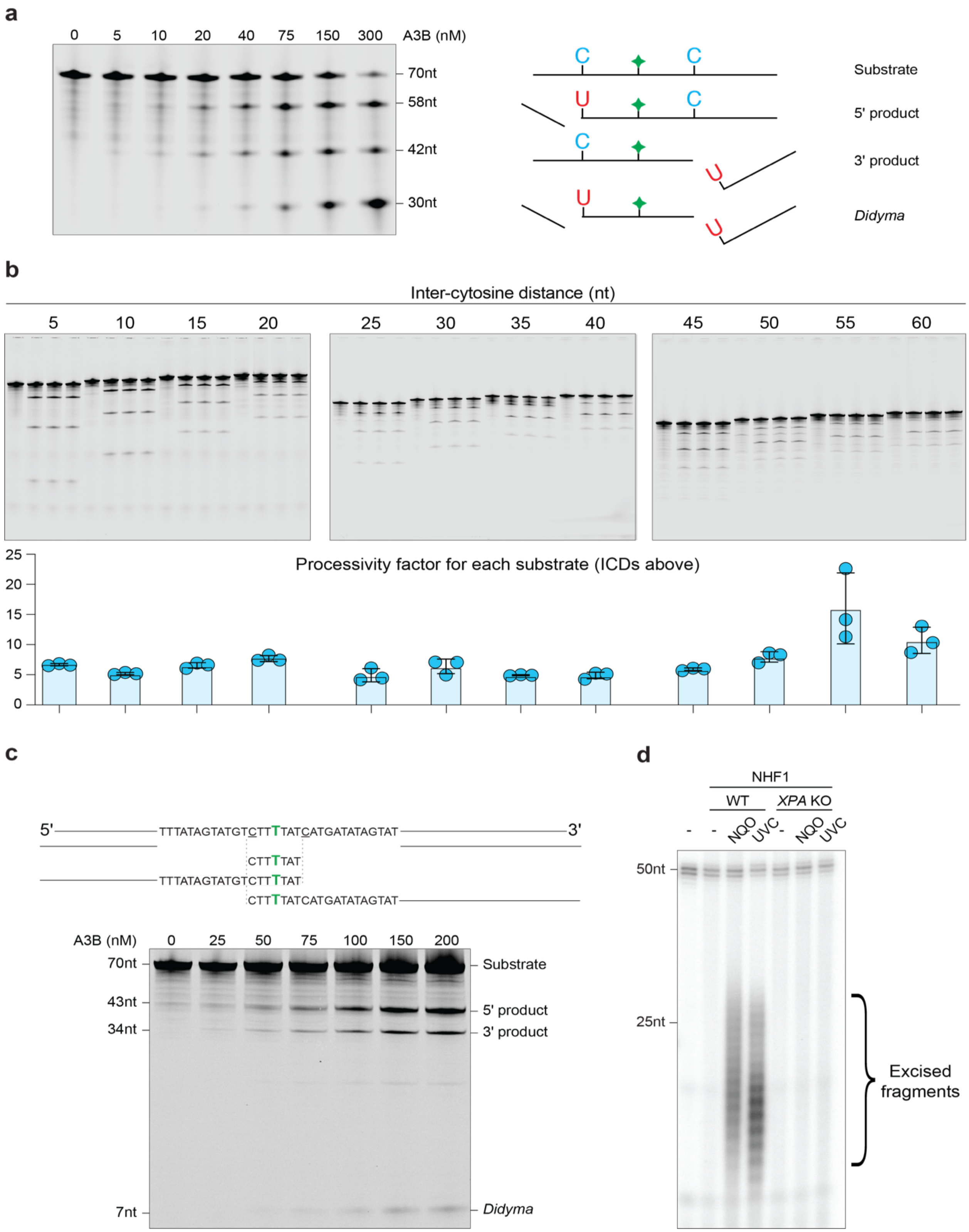


**Supplementary Fig. 3. Additional biochemical experiments.**

(**a**) Gel image of A3B-catalyzed ssDNA deamination reactions (enzyme concentrations indicated,

800nM substrate, 15min incubation). Schematics of the ssDNA substrate and each of the unique deamination products are shown to the right of the gel image (green diamond=thymine with a fluorescein attached for visualization). Each reaction includes a molar excess of EndoQ, which cleaves near-instantaneously on the 5'-side of each nascent deoxy-uracil.

(**b**) Gel images showing A3B activity on ssDNA substrates with 2 target cytosines separated by the indicated number of nucleotides (5 to 60nt). The left lane of each group of 4 reactions has no A3B to assess background signal (slightly different for each substrate). The next 3 lanes of each group show 3 independent reactions of 25nM A3B, 800nM substrate, and 15min incubation (and excess EndoQ as above). The inter-cytosine mutation distances are indicated above each substrate group. A3B processivity factors for each inter-cytosine distance are plotted below (mean with 95% confidence intervals from 3 independent reactions). The A3B processivity factor over all conditions is 7.3 +/- 0.6 (mean +/- SEM).

(**c**) A3B-catalyzed deamination of twin TCW motifs in a 32nt-gapped region. Schematics of the gapped substrate and 3 visible deamination products are shown above the gel image (polarity indicated; green T has a fluorescein label; target cytosines are underlined).

(**d**) Phosphoimage of $\alpha$-$^{32}$P-labeled low-molecular-weight DNA extracted 2h after exposing WT or *XPA* KO NHF1 cells to 5µM NQO or 20J/m$^2$ UVC as indicated. Each labeling reaction included 2fmol of a 50nt ssDNA as an internal control.

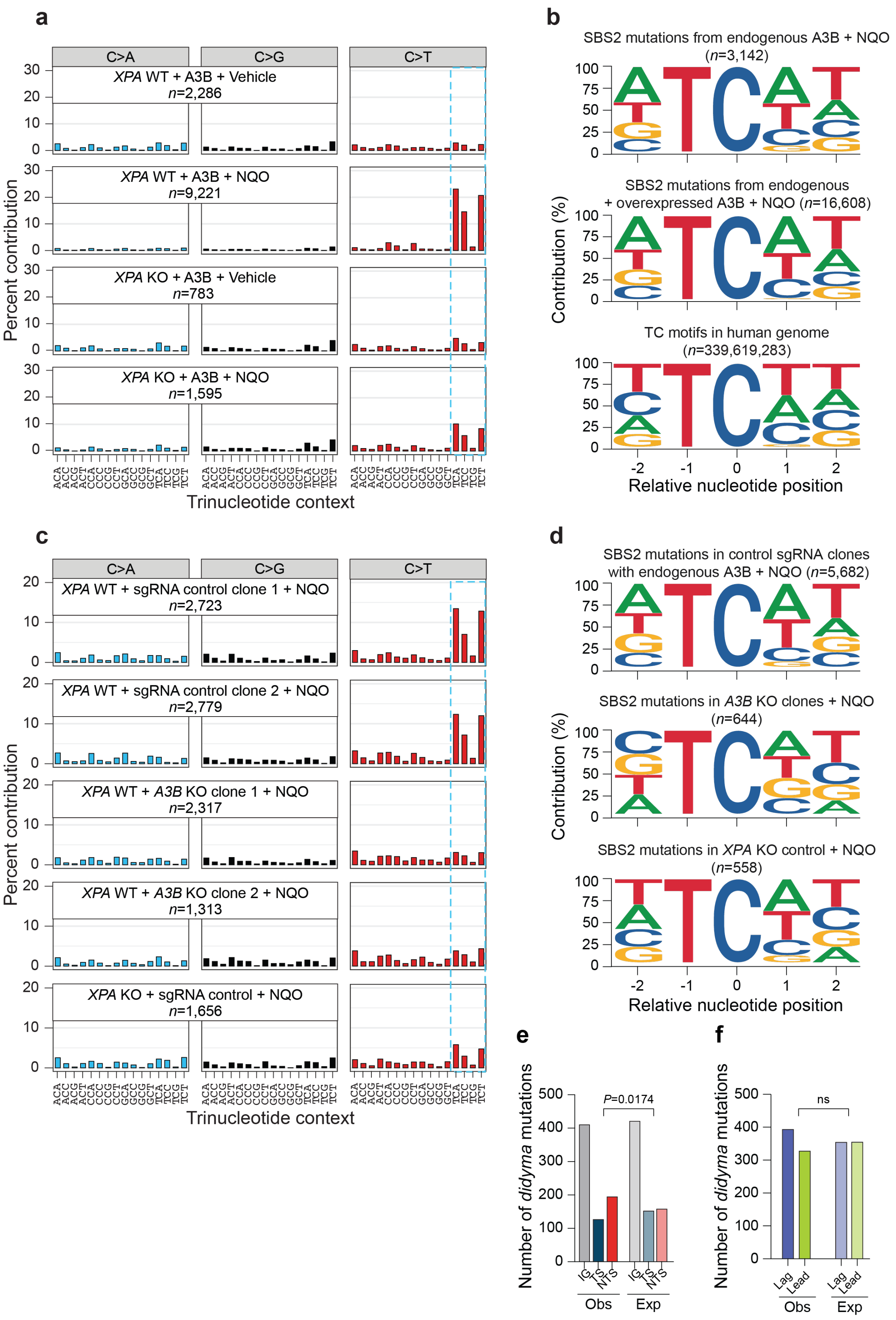
a
C>A
C>G
C>T
XPA WT + A3B + Vehicle
n=2,286
XPA WT + A3B + NQO
n=9,221
XPA KO + A3B + Vehicle
n=783
XPA KO + A3B + NQO
n=1,595
Percent contribution
Trinucleotide context
b
SBS2 mutations from endogenous A3B + NQO
(n=3,142)
SBS2 mutations from endogenous
+ overexpressed A3B + NQO (n=16,608)
TC motifs in human genome
(n=339,619,283)
Contribution (%)
Relative nucleotide position
c
XPA WT + sgRNA control clone 1 + NQO
n=2,723
XPA WT + sgRNA control clone 2 + NQO
n=2,779
XPA WT + A3B KO clone 1 + NQO
n=2,317
XPA WT + A3B KO clone 2 + NQO
n=1,313
XPA KO + sgRNA control + NQO
n=1,656
Percent contribution
Trinucleotide context
d
SBS2 mutations in control sgRNA clones
with endogenous A3B + NQO (n=5,682)
SBS2 mutations in A3B KO clones + NQO
(n=644)
SBS2 mutations in XPA KO control + NQO
(n=558)
Contribution (%)
Relative nucleotide position
e
Number of didyma mutations
P=0.0174
IG
TS
NTS
Obs
Exp
f
Number of didyma mutations
ns
Lag
Lead
Obs
Exp

2145

**Supplementary Fig. 4. Context of A3B mutations following NQO treatment.**

(**a**) Trinucleotide SBS mutation distributions of SW1573 cells from the indicated conditions. *n*-values represent the C/G base substitution mutation total per diploid genome.

(**b**) Pentanucleotide context of TC-to-TT and TC-to-TG mutations in NQO-treated SW1573 cells expressing endogenous A3B (top) or a combination of endogenous and exogenous A3B (middle), in comparison to the local pentanucleotide context of all TC motifs in the human genome (bottom).

(**c**) Trinucleotide SBS mutation distributions of SW1573 cells from the indicated conditions. *n*-values represent the C/G base substitution mutation total per diploid genome.

(**d**) Pentanucleotide context of TC-to-TT and TC-to-TG mutations in NQO-treated SW1573 cells from control clones expressing endogenous A3B (top) in comparison to those in two *A3B* knockout clones (middle) and one *XPA* knockout clone (bottom).

(**e**) Quantification of APOBEC signature *didyma* mutations in the indicated regions from NQO-treated SW1573 cells expressing endogenous A3B (IG, intergenic; TS, transcribed strand; non-transcribed strand, NTS). To have sufficient numbers of *didyma* for this transcription association analysis, duplex sequencing *didyma* mutations were pooled from four conditions (eGFP and catalytic mutant controls in Fig. 5a-d and two sgRNA control clones in Fig. 5e-h).

(**f**) APOBEC signature *didyma* mutations from NQO-treated SW1573 cells expressing endogenous A3B do not associate with DNA replication leading or lagging strand (lead and lag, respectively). *Didyma* mutation data pooled for this analysis as in panel-e.

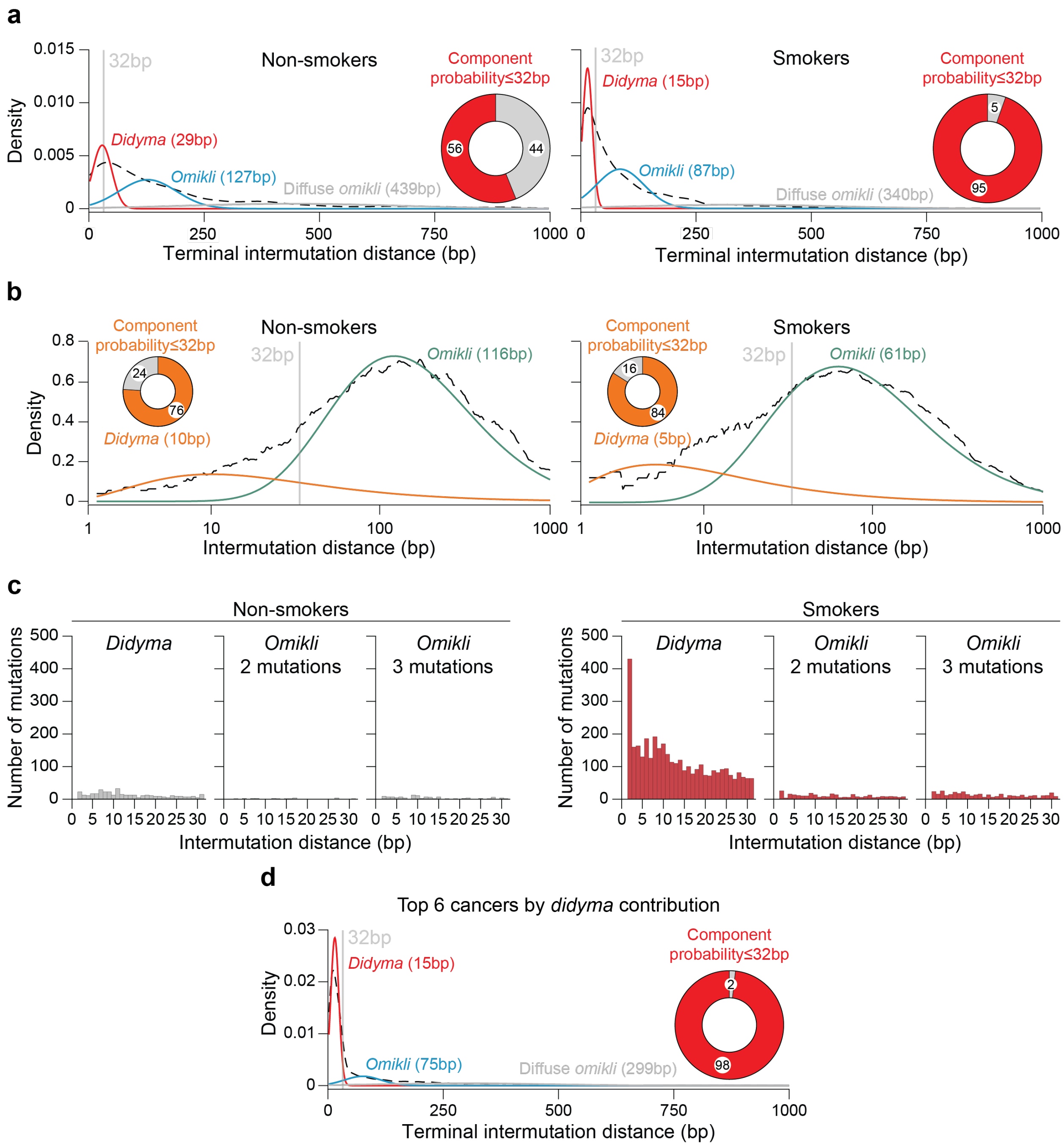
a
Non-smokers
Smokers
Density
0.015
0.010
0.005
0
32bp
Didyma (29bp)
Omikli (127bp)
Diffuse omikli (439bp)
Component probability≤32bp
56
44
Didyma (15bp)
Omikli (87bp)
Diffuse omikli (340bp)
5
95
0
250
500
750
1000
Terminal intermutation distance (bp)
b
Component probability≤32bp
24
76
Didyma (10bp)
Omikli (116bp)
16
84
Didyma (5bp)
Omikli (61bp)
0.8
0.6
0.4
0.2
1
10
100
1000
Intermutation distance (bp)
c
Number of mutations
Didyma
Omikli 2 mutations
Omikli 3 mutations
500
400
300
200
100
0 5 10 15 20 25 30
d
Top 6 cancers by didyma contribution
0.03
0.02
0.01
Didyma (15bp)
Omikli (75bp)
Diffuse omikli (299bp)
2
98

**Supplementary Fig. 5. Probabilistic models of clustered APOBEC signature mutations.** (**a**) Gaussian mixture model of *didyma* and *omikli* distance distributions in non-smokers and smokers using a 3-component-based model, representing terminal IMDs up to 1,000bp. The 3 components (red, blue, and gray) are overlaid on top of the observed distribution of all *didyma* and *omikli* combined (black dashed line) from the lung and head & neck cancers in Fig. 6 (*n*=144, smokers; *n*=32, non-smokers). Inset pie charts represent the probability of the shortest component having a width ≤32bp. Components are labeled with inferred clustered mutational event and peak value. These plots are the same as those in Fig. 6i, except the x-axis is expanded to 1,000bp to include diffuse *omikli*.

(**b**) Gamma mixture model of the densities of combined *didyma* and *omikli* IMD distributions in non-smokers (left) and smokers (right) using two components, representing all IMDs up to 1,000bp. The two-component models (orange and green) are overlaid on top of all *didyma* and *omikli* IMDs (black) from the lung and head & neck cancers in Fig. 6 (*n*=144, smokers; *n*=32, non-smokers). Inset proportions represent the probability of the shortest component having an IMD ≤32bp. Components are labeled with their inferred mutagenic process, along with their IMD peak.

(**c**) Number of mutations that occur at each IMD from 2-31bp in indicated clustered mutation groups in non-smokers (top) and smokers (bottom).

(**d**) Gaussian mixture model of the densities of combined *didyma* and *omikli* terminal IMD distributions in the top-6 tumor types with the highest number of *didyma* events (according to Fig. 7a (*i.e.*, metastatic urothelial, primary bladder-TCC, LUAD from smokers, LUAD from non-smokers, cervix-SCC, and biliary adenocarcinoma) using three components, representing all terminal IMDs up to 1,000bp. Color scheme and labels as in panel-a.

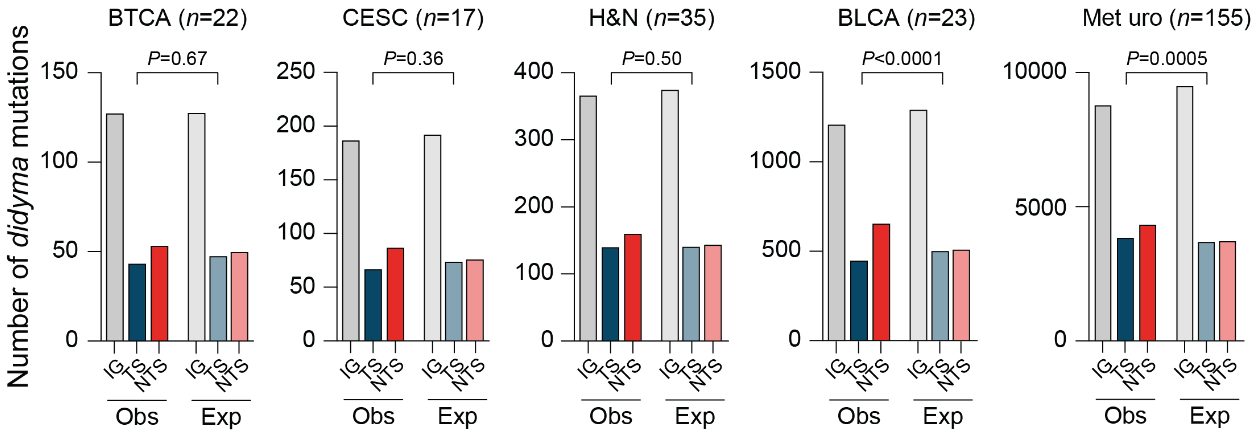


**Supplementary Fig. 6. *Didyma* enrichment on the non-transcribed strand of genes.** Quantification of *didyma* in the indicated regions across the cancer cohorts with the most *didyma* observed in Fig. 7, along with larynx and hypopharynx head & neck cancers from mutographs as in Fig. 6. Each tumor cohort is indicated above the relevant plot and ordered left-to-right from lowest to highest number of *didyma* (BTCA, biliary tract carcinoma; CESC, cervical squamous cell carcinoma; H&N, head and neck; BLCA, bladder carcinoma; Met uro, metastatic urothelial cancer). Observed mutations are from tumor data and expected from simulations (*n*-values represent the number of tumors analyzed in each cohort; *P*-values, two-sided Fisher's exact test).
